# Redox-active cysteines in TGACG-BINDING FACTOR 1 (TGA1) do not play a role in salicylic acid- or pathogen-induced expression of TGA1-regulated target genes in *Arabidopsis thaliana*

**DOI:** 10.1101/2020.01.30.926758

**Authors:** Jelena Budimir, Katrin Treffon, Aswin Nair, Corinna Thurow, Christiane Gatz

**Affiliations:** Albrecht-von-Haller-Institut für Pflanzenwissenschaften, Georg-August-Universität Göttingen, Julia-Lermontowa-Weg 3, D-37077 Göttingen, Germany

**Keywords:** defense responses, NPR1, redox regulation, salicylic acid, TGA transcription factors

## Abstract

Salicylic acid (SA) is an important signaling molecule of the plant immune system.
SA biosynthesis is indirectly modulated by the closely related transcription factors TGA1 (TGACG-BINDING FACTOR 1) and TGA4. They activate expression of *SARD1* (*SYSTEMIC ACQUIRED RESISTANCE DEFICIENT1*), the gene product of which regulates the key SA biosynthesis gene *ICS1* (*ISOCHORISMATE SYNTHASE 1*).
Since TGA1 interacts with the SA receptor NPR1 (NON EXPRESSOR OF PATHOGENESIS-RELATED GENES 1) in a redox-dependent manner and since the redox state of TGA1 is altered in SA-treated plants, TGA1 was assumed to play a role in the NPR1-dependent signaling cascade. Here we identified 193 out of 2090 SA-induced genes that require TGA1/TGA4 for maximal expression after SA treatment. One robustly TGA1/TGA4-dependent gene encodes for the SA hydroxylase DLO1 (DOWNY MILDEW RESISTANT 6-LIKE OXYGENASE 1) suggesting an additional regulatory role of TGA1/TGA4 in SA catabolism.
Expression of TGA1/TGA4-dependent genes in mock/SA-treated or Pseudomonas-infected plants was rescued in the *tga1 tga4* double mutant after introduction of a mutant genomic *TGA1* fragment encoding a TGA1 protein without any cysteines. Thus, the functional significance of the observed redox modification of TGA1 in SA-treated tissues has remained enigmatic.

**SIGNIFICANCE STATEMENT:** Previous findings demonstrating a redox-dependent interaction between transcription factor TGA1 and NPR1 attracted considerable attention. Here we show that TGA1 can act in the NPR1- and SA-dependent signaling cascade, but that its SA-regulated redox-active cysteines do not affect its function in this process.

## Introduction

Redox reactions drive all energy-converting processes in living organisms. To adjust metabolic and regulatory processes to the prevailing redox state, proteins possess reactive cysteines which can be subject to various oxidative modifications. Prominent examples of proteins regulated by these so called thiol switches are enzymes of the Calvin Cycle, which become inactivated during the night when less reducing power is available in the chloroplast (Michelet *et al.*, 2013). Conversely, oxidation of yeast transcription factor yAP1 leads to its accumulation in the nucleus, where it activates genes of the anti-oxidative system (Delaunay *et al.*, 2000).

Plant immune responses are associated with complex changes in the cellular redox state. The defence hormone salicylic acid (SA), for instance, promotes the production of reactive oxygen or nitrogen species, while on the other hand inducing genes of the anti-oxidative system, like e.g. oxidoreductases or glutathione biosynthesis genes (Herrera-Vasquez *et al.*, 2015). Redox signals affect the activity of the important regulatory protein NON EXPRESSOR OF PATHOGENESIS-RELATED GENE1 (NPR1). NPR1 controls many processes that are induced by elevated SA levels. In SA-treated tissues, NPR1 becomes first nitrosylated, which is a prerequisite for the formation of intermolecular disulfide bonds. These force the protein into the inactive oligomeric form, which resides in the cytosol (Mou *et al.*, 2003; Tada *et al.*, 2008). On the other hand, transcription of the small oxidoreductase *THIOREDOXIN h5* is activated, which in turn reduces the disulfide bonds resulting in monomerization and nuclear translocation of NPR1 (Spoel *et al.*, 2009). In the nucleus, NPR1 protein levels are regulated by NPR3 and NPR4 in an SA-dependent manner (Fu *et al.*, 2012). All three NPR1 proteins bind SA, which is essential for their regulatory function (Fu *et al.*, 2012; Ding *et al.*, 2018). NPR1 interacts with TGACG-binding (TGA) transcription factors TGA2, TGA3, TGA5 and TGA6 to induce the expression of defence genes (Zhang *et al.*, 2003; Saleh *et al.*, 2015), while NPR3 and NPR4 function as repressors (Ding *et al.*, 2018).

TGA factors form a family of ten members which are grouped into five clades (Gatz, 2013). The partially redundant clade-II TGAs (TGA2, TGA5 and TGA6) function together with NPR1 in the context of the immune response “systemic acquired resistance” (SAR) (Zhang *et al.*, 2003). TGA3 is – like NPR1 – required for basal resistance against the bacterial pathogen *Pseudomonas syringae* pv. *maculicola* ES4356 (*Psm*) (Kesarwani *et al.*, 2007). Since NPR1 is sumoylated after SA treatment and since TGA3 only interacts with sumoylated NPR1, it has been concluded that TGA3 and NPR1 functionally interact *in vivo* (Saleh *et al.*, 2015). The SA marker gene *PR1* (*PATHOGENESIS-RELATED 1*) has been used as an example to provide evidence that the described NPR1/TGA interactions occur at TGA binding sites in SA-responsive promoters.

Initial studies also suggested that NPR1 and clade-I TGAsc (TGA1 and TGA4) act in the same pathway. First, TGA1/TGA and NPR1 are required for basal resistance against *Psm*; second, TGA1 interacts with NPR1 only if an inhibitory internal disulfide bridge between cysteine residues 260 and 266 of TGA1 is not formed; third, the interaction between NPR1 and TGA1 promotes its binding to DNA and fourth, TGA1 is partially oxidized in untreated leaves and becomes reduced after SA treatment. Based on these circumstantial pieces of evidence, models presenting redox-modulated TGA1 interacting with NPR1 at SA-responsive promoters were published in numerous reviews and book chapters (Eckardt, 2003; Pieterse & Van Loon, 2004; Li & Zachgo, 2009; Moore *et al.*, 2011; Chi *et al.*, 2013; Li & Loake, 2016; Gullner *et al.*, 2017).

However, the functional significance of TGA1 for the expression of SA/NPR1-regulated genes and the role of the often cited redox modulation has not yet been conclusively demonstrated. Using microarray analysis of SA-treated plants, Shearer *et al.* observed that expression of 584 of the 629 SA-induced NPR1-dependent genes were independent from TGA1/TGA4 and that basal levels of the remaining 45 genes including *PR1* were up-regulated in *tga1 tga4* (Shearer et al., 2012). This implicated that oxidized TGA1/TGA4, which has a low DNA binding activity at least *in vitro*, would repress these genes and repression would be released upon the interaction of reduced TGA1/TGA4 with NPR1. To explain the susceptibility of the *tga1 tga4* mutant, an NPR1-independent defence mechanism was postulated and confirmed by the higher susceptibility of the *npr1 tga1 tga4* mutant as compared to *npr1* and *tga1 tga4* mutants. A very recent study explained the susceptibility of the *tga1 tga4* mutant by lower SA and pipecolic acid levels after *Psm* infections. These are due to the reduced expression of the master regulator of SA and pipecolic acid biosynthesis, SARD1 (SAR DEFICIENT 1) in *Psm*-infected *tga1 tga4* plants. Chromatin immunoprecipitation experiments confirmed *SARD1* as a direct target gene of TGA1/TGA4 (Sun *et al.*, 2018).

We decided to re-address the question whether the redox-regulated cysteines in TGA1 play a regulatory role. Since in our hands, basal expression of *PR1* was not enhanced in *tga1 tga4*, we performed again transcriptome analysis to identify TGA1/TGA4-regulated genes. RNAseq analysis provided a number of SA-induced NPR1-dependent genes that were less expressed in *tga1 tga4*, with *SA-3-HYDROXYLASE (S3H)/DOWNY MILDEW RESISTANT 6-LIKE OXYGENASE1 (DLO1)* (Zhang *et al.*, 2013; Zeilmaker *et al.*, 2015) being the most robust TGA1/TGA-dependent gene. Under the conditions tested so far, no evidence for a function of the previously postulated redox switch of TGA1 for the regulation of *DLO1* and other genes was obtained.

## Materials and Methods

### Plant material and cultivation

All plants used in this study are in the *Arabidopsis thaliana* Columbia background. Plants were cultivated in individual pots containing steamed soil (Archut, Fruhstorfer Erde, T25, Str1fein, soaked twice with 0.2% Wuxal Super (Manna, Ammerbuch-Pfäffingen, Germany)) in a growth cabinet at 22°C with a 12-h-day/12h-night cycle and a photon flux density of 100-120 µmol photons m^−2^ s^−1^ and 60% relative humidity. Genotypes used in the study and corresponding references are: *npr1*-*1* (Cao *et al.*, 1997), *sard1-1 cbp60g-1* (Zhang et al., 2010), *sid2-2* (Wildermuth et al., 2001), *tga1 tga4* (Kesarwani et al., 2007), *tga2 tga5 tga6* (Zhang *et al.*, 2003). The *sid2 tga1 tga4* triple mutant and the *tga1 tga2 tga4 tga5 tga6* pentuple mutants were generated through crossings of the respective above mentioned genotypes. Mutants were either obtained from the Nottingham Arabidopsis Stock Center or from Prof. Dr. Yuelin Zhang (UBC Vancouver, Canada; *sard1 cbp60g*; *tga1 tga4*; *tga2 tga5 tga6*).

### Salicylic acid treatment of soil-grown plants

Four-week-old plants were sprayed either with water or with a freshly prepared 1 mM solution of sodium salicylate (Sigma) until the whole rosette was equally moist. Treatment was conducted 1 hour after the subjective dawn and samples were collected at eight hours after treatment.

### Pathogen infection assays

*Pseudomonas syringae* pv. *maculicola* ES4356 (*Psm*) was cultivated at 28°C in King’s B medium. Overnight cultures were diluted in 10 mM MgCl_2_ to the final optical density at 600 nm (OD_600_) of 0.005. 10 mM MgCl_2_ (mock) or the diluted bacteria were hand-infiltrated into three leaves of five-week-old plants. Two days after this primary infection, three younger upper leaves were infiltrated again with a *Psm* solution (OD_600_ of 0.005) in 10 mM MgCl_2._ These leaves were harvested for RNA extraction after eight hours post infection. Pathogen infiltrations were generally conducted at one hour after the subjective dawn.

### Other methods

Construction of recombinant plasmids, transcriptome analysis, quantitative reverse transcription (qRT)-PCR, transient expression analysis in Arabidopsis protoplasts, Western blot analysis, and accession numbers can be found in Methods S1. Primer sequences are depicted in Table S1. Maps and sequences of plasmids can be found in Notes S1.

## Results

### TGA1/TGA4 positively regulate a subgroup of salicylic acid-induced genes

In order to address the question whether the disulfide bridge-forming cysteines of TGA1 that become reduced in SA-treated plants indeed play a role for accurate transcription of SA-responsive genes, we first tested SA-induced expression of *SARD1*, which has been recently identified as a target gene of TGA1/TGA4 (Sun et al., 2018). However, in contrast to *Psm* infections, spraying with 1 mM SA resulted in TGA1/TGA4-independent *SARD1* expression (Fig. 1). Under these conditions, *SARD1* was controlled by the well-established SA-responsive regulatory module which consists of NPR1 and clade-II TGAs TGA2/TGA5/TGA6. Still, it has to be noted that basal levels of *SARD1* were lower in the *tga1 tga4* and the *npr1* mutants than in wild-type plants suggesting that residual basal levels of TGA1/TGA4 and NPR1 stimulate basal *SARD1* expression.

**Fig. 1.**
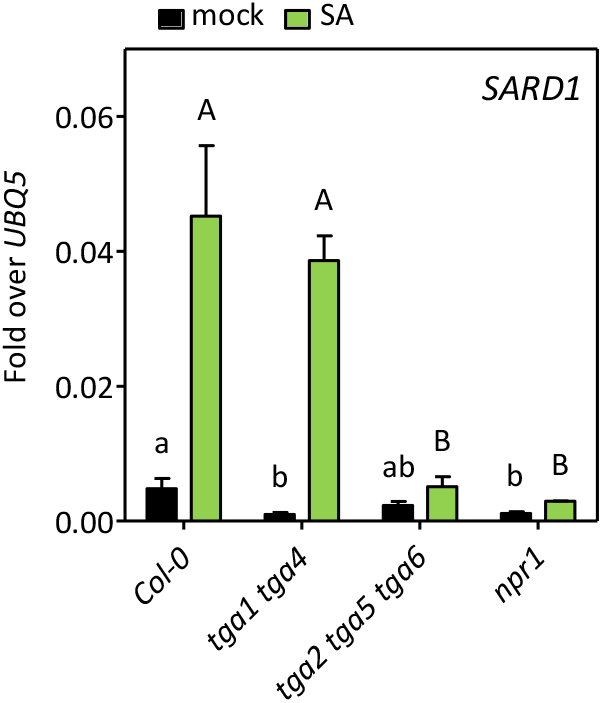
Salicylic acid (SA)-induced *SARD1* expression is independent from TGA1/TGA4. qRT-PCR analysis of *SARD1* transcript levels in wild-type (Col-0), *tga1 tga4*, *tga2 tga5 tga6* and *npr1* plants. Four-week-old soil-grown plants were sprayed with water (mock) or 1 mM SA and tissue was harvested after 8 hours. Transcript levels were normalized to the transcript levels of *UBQ5*. Bars represent the average ± SEM of four to five biological replicates, each replicate representing three leaves from one plant. Statistical analysis was performed using one-way ANOVA followed by Tukey’s post hoc test for mock- and SA-treated samples separately. Lowercase letters indicate significant differences (*P* < 0.05) between mock-treated samples; uppercase letters indicate significant differences (*P* < 0.05) between SA-treated samples.

In contrast to previously published observations (Lindermayr *et al.*, 2010; Shearer *et al.*, 2012), basal *PR1* expression was not enhanced in the *tga1 tga4* mutant and SA-induced *PR1* transcript levels were only slightly affected (see below, Fig. 4). Therefore, we performed transcriptome analysis of RNA harvested from leaves of mock- and SA-treated plants. We compared the expression pattern of *sid2* and *sid2 tga1 tga4* rather than that of wild-type and *tga1 tga4* because we wanted to avoid any possible influence of TGA1/TGA4 on endogenous SA biosynthesis. Moreover, we aimed to reduce fluctuations in gene expression due to environmental factors affecting endogenous SA levels in different experiments. Four-week-old plants were sprayed either with water or with 1 mM SA. Eight hours after treatment, three leaves of five individual plants were collected and total RNA was isolated. The experiment was repeated four times with batches of independently grown plants. Thus, the RNA from 15 leaves of five plants served as one replicate and replicates originated from four independent experiments.

Principal component analysis (PCA) results in clusters of samples with a similar expression pattern and thus yields a first impression of the global structure of the data set. The clusters from *sid2* and *sid2 tga1 tga4* plants treated with water showed a clear separation (Fig. S1) indicating that the transcriptomes of both genotypes are different even in the absence of ICS1-derived metabolites. The clusters from SA-treated plants indicate that both genotypes respond to SA. Since our aim was to identify target genes of TGA1/TGA4 after SA treatment, we focused on those 2090 genes that were induced (log2 fold change (FC) >1) by SA in *sid2* (Table S2).

Fourtyone % (864 genes) of the 2090 SA-induced genes showed a differential expression pattern in *sid2 tga1 tga4*. These 864 genes fall in two major groups (Fig. 2a). Genes with lower expression values in the *sid2 tga1 tga4* plants as compared to *sid2* establish the “green” group (346 genes). Three major subgroups were identified based on reduced gene expression in *sid2 tga1 tga4* either after SA treatment (119), mock and SA treatment (71), or only after mock treatment (153). Only three genes were less expressed in SA-treated leaves while background levels were elevated. The two major subgroups of the “red” group, which comprises 518 genes that are higher expressed in *sid2 tga1 tga4* as compared to *sid*2, contain genes that have higher expression values only in the mock situation (401) and genes that had elevated transcript levels in mock- and SA-treated plants (114). Three genes were hyper-induced upon SA treatment and have wild-type transcript levels upon mock treatment.

**Fig. 2.**
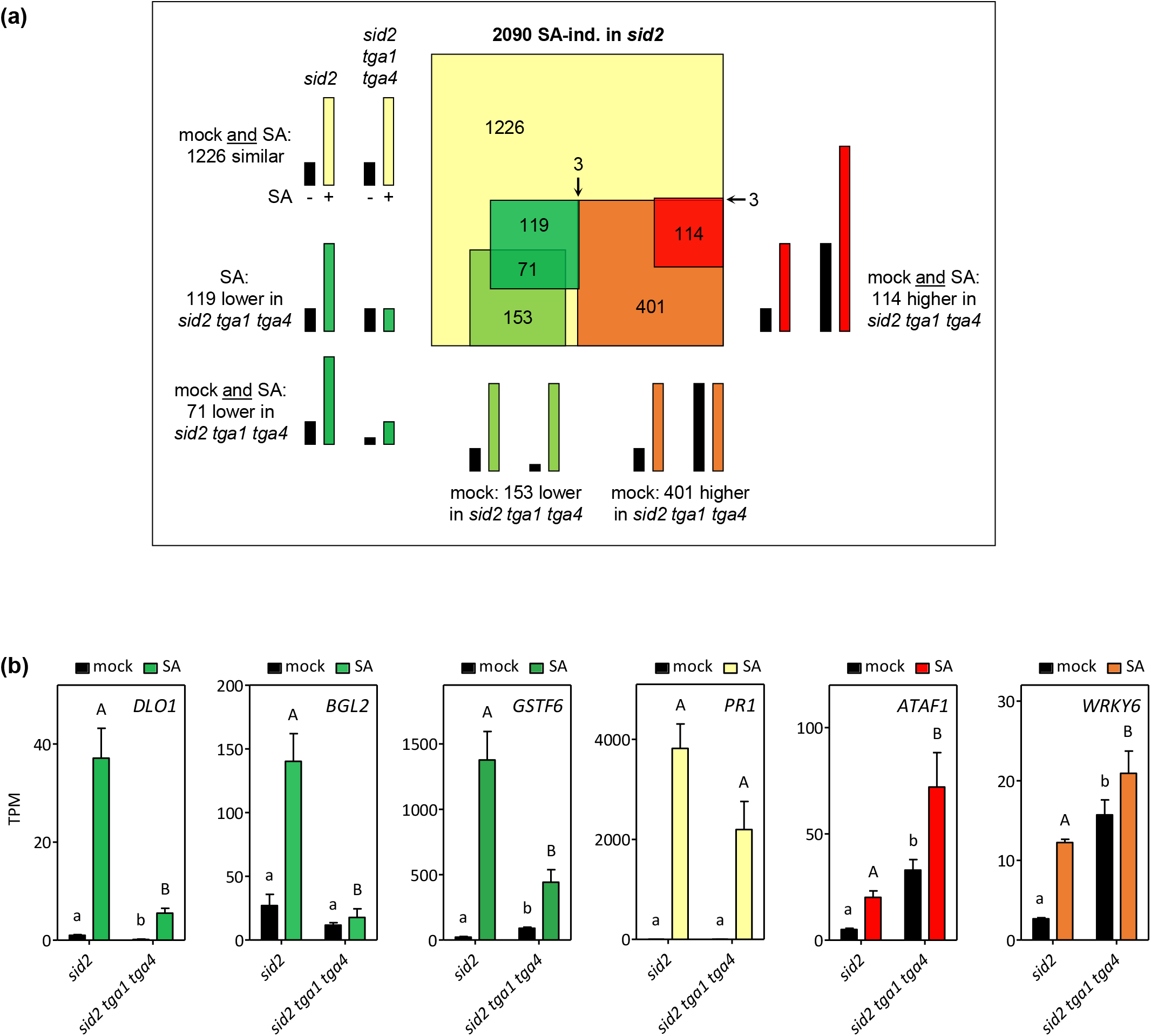
Almost half of the salicylic acid (SA)-induced transcripts are modulated by TGA1/TGA4 under basal or induced conditions. (a) Euler diagram of 2090 SA-inducible genes identified at 8 h after spraying with 1 mM SA as compared to water (mock) treatment in *sid2*. Different square sizes represent the number of genes with significantly different (log2 fold change (FC) ≥1 or log2 FC ≤ −1, *P* < 0.05) transcript levels in *sid2 tga1 tga4* under either mock, SA-induced- or under both conditions. Sketches are drawn to visualize the expression pattern in the respective groups. (b) Relative expression of selected genes as identified by RNAseq analysis of four-week old Arabidopsis *sid2* and *sid2 tga1 tga4* plants treated with water (mock) or 1 mM SA. Bars represent the average of Transcripts Per Kilobase Million (TPM) ± SEM of four biological replicates of each genotype, with each replicate representing three leaves of five plants of one independent experiment. Statistical analysis was performed using unpaired Student’s t-test (two-tailed) for mock- and SA-treated samples separately. Lowercase letters indicate significant differences (*P* < 0.05) between mock-treated samples; uppercase letters indicate significant differences (*P* < 0.05) between SA-treated samples.

Figure 2b shows relative expression levels of representative genes of the green and the red group. *DLO1* encodes for an SA hydroxylase that is involved in dampening the immune response by inactivating SA (Zhang *et al.*, 2013; Zeilmaker *et al.*, 2015). Its expression responded strongly to SA (30-fold) and we observed a 7-fold reduction of expression in the *tga1 tga4* mutant, both under basal conditions and after SA treatment. Thus, the induction factor after SA treatment was not changed, suggesting that TGA1/TGA4 act as amplifiers under both conditions. In contrast, induction factors were lower for other genes in *sid2 tga1 tga4* as compared to *sid2* (Table S3). *ß-1,3 GLUCANASE* (*BGL2*) for example was induced by a factor of 4.3 in SA-treated *sid2*, and by a factor of 1.4 in SA-treated *sid2 tga1 tga4*. Another example is *GLUTATHIONE S-TRANSFERASE F 6* (*GSTF6*), which was induced by a factor of 52 in *sid2* and by a factor of 4.3 in *sid2 tga1 tga4*. The well-known gene of the NPR1-dependent SA response gene *PR1* barely missed the cut-off for being differentially expressed in *sid2 tga1 tga4* versus *sid2* in the RNAseq analysis, but its expression was still modulated by TGA1/TGA4. Two genes with elevated expression levels (*ATAF1* and *WKRY6*) in *sid2 tga1 tga4* as compared to *sid2* are displayed as well. These genes code for transcription factors.

### The TGACGTCA motif is specifically enriched in SA-induced genes that are positively regulated by TGA1/TGA4

The ideal binding site for TGA factors is the palindromic sequence TGAC/GTCA, which is an extended C-box (GAC/GTC) (Izawa *et al.*, 1993; Qin *et al.*, 1994). However, the pentamer TGAC/G is sufficient for binding. Moreover, at least TGA1 binds to the A-box (TAC/GTA) *in vivo* (Wang *et al.*, 2019) and tobacco TGA1a binds to A- and G- (CAC/GTG) boxes *in vitro* (Izawa *et al.*, 1993). Therefore, we tested whether any of these potential binding sites is specifically enriched in promoters that are either lower or higher expressed in *sid2 tga1 tga4* as compared to *sid2*. To this end, the 1-kb sequences upstream of the predicted transcriptional start sites were scanned using the Motif Mapper *cis* element analysis tool (Berendzen *et al.*, 2012). As displayed in Fig. 3, all potential TGA1 binding sites are enriched in the 2090 SA-inducible promoters as compared to promoters arbitrarily selected from the whole genome. At least the enrichment of the extended C-boxes was expected since most of the 2090 genes are likely to be regulated by NPR1 acting in concert with TGA2/TGA5/TGA6 or TGA3 (Wang *et al.*, 2006). However, when comparing the relative frequency of these motifs in TGA1/TGA4-regulated promoters with their relative frequency in the 2090 SA-regulated genes, an enrichment of the TGACGTCA motif was detected in the group of those 346 genes that required TGA1/TGA4 for maximal expression. *DLO1*, for instance, contains the TGACGTCA palindrome at position −72 bps with respect to the transcriptional start site. Likewise, *SARD1*, which was slightly but significantly activated by TGA1/TGA4 in the absence of SA (Fig. 1) and after *Psm* infections (Sun *et al*., 2018), contains a TGACGTCA motif at position - 212 bps. The G-box is significantly depleted in promoters being less activated in *sid2 tga1 tga4* plants. Conversely, the G-box and the A-box are enriched in promoters being de-repressed in this mutant. It might well be that more efficient transcriptional activators compete with TGA1 for A- and G-boxes.

**Fig. 3.**
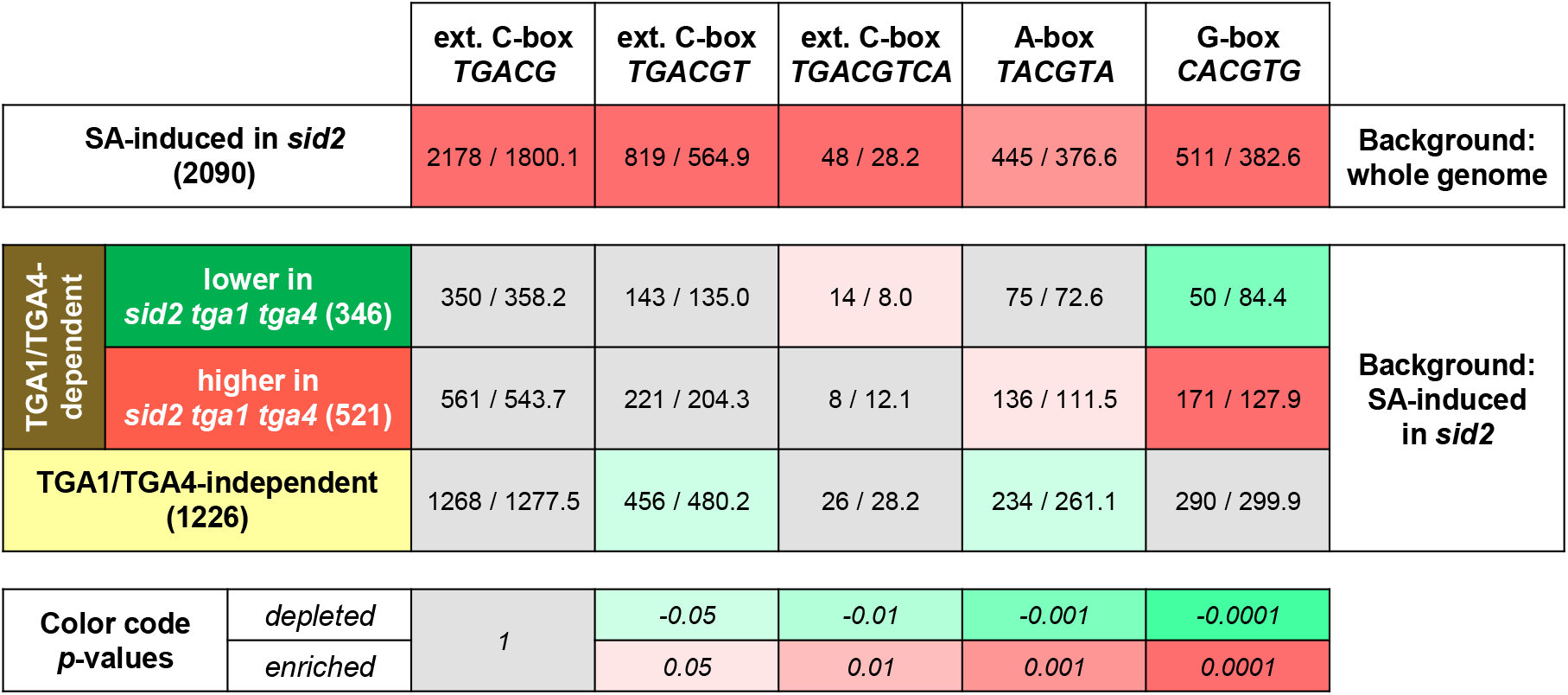
TGA1/TGA4-modulated salicylic acid (SA)-induced genes are characterized by a higher incidence of the *TGACGTCA* motif. The occurrence of enriched motifs was determined in the 1-kb sequences upstream of the 5’-untranslated regions. Numbers before the slash represent the total number of occurrences of the given motif within the indicated set. Numbers behind the slash represent the expected number of occurrences in a set of randomly chosen promoters from either the whole genome (upper panel) or the set of 2090 SA-induced genes (lower panel). The corresponding enrichment *P*-values are color coded (grey: not significant, green: significantly depleted, red: significantly enriched). ext. – extended.

In conclusion, the enrichment analysis indicates that promoters containing the TGACGTCA motif might be preferred targets of TGA1/TGA4.

### NPR1 and clade-II TGA factors are required for expression of selected TGA1/TGA4-dependent genes

Since the SA-induced redox-modification of TGA1 alters the interaction with NPR1, we tested, whether expression of the selected TGA1/TGA4-dependent genes (Fig. 2b) requires NPR1. We also included the *tga2 tga5 tga6* mutant, since NPR1 has been functionally associated with clade-II TGAs (Zhang *et al.*, 2003). Col-0 wild-type and *tga1 tga4* mutant plants were included in the experiment. As observed before for *sid2 tga1 tga4* vs. *sid2*, the four marker genes that are positively regulated by TGA1/TGA4, were less expressed in *tga1 tga4* vs. Col-0 (Fig. 4). However, the elevated background levels of *WRKY6* and *ATAF1* were not observed in the presence of a functional *SID2* allele. Expression of all genes with the exception of *ATAF1* was significantly regulated by NPR1 (Fig. 4). Since expression also depended on clade-II TGAs, a functional connection between TGA1/TGA4 and NPR1 could not be inferred.

**Fig. 4.**
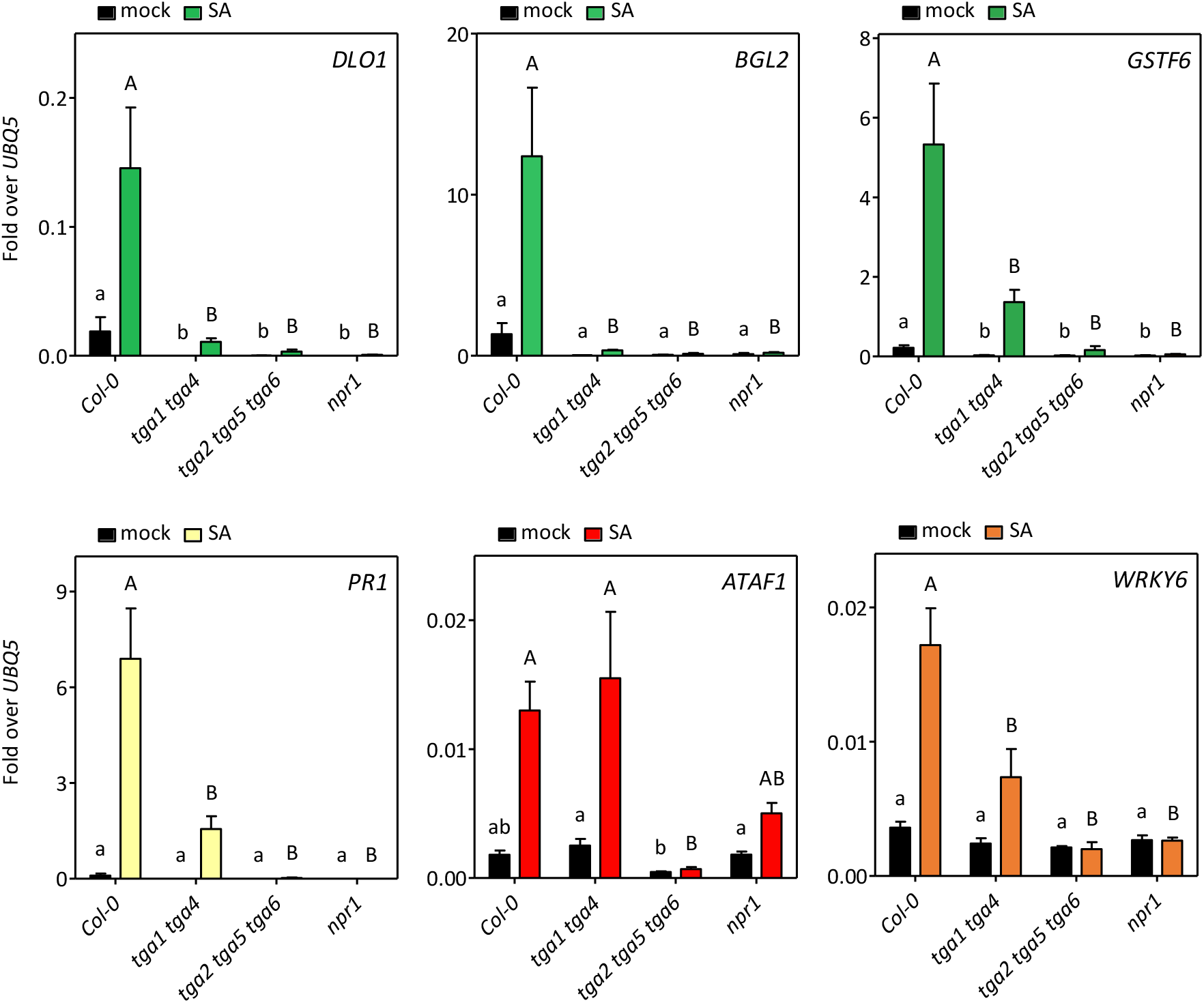
TGA1/TGA4-modulated genes are stringently controlled by NPR1 and clade II TGAs. qRT-PCR analysis of transcript levels of six TGA1/TGA4-modulated genes in wild-type (Col-0), *tga1 tga4*, *tga2 tga5 tga6* and *npr1* plants. Four-week-old plants were sprayed either with water (mock) or 1 mM salicylic acid (SA) and further incubated for 8 h. Transcript levels were normalized to transcript levels of *UBQ5*. Bars represent the average ± SEM of four to five biological replicates, each replicate representing three leaves from one plant. Statistical analysis was performed using one-way ANOVA followed by Tukey’s post hoc test for mock- and SA-treated samples separately. Lowercase letters indicate significant differences (*P* < 0.05) between mock-treated samples; uppercase letters indicate significant differences (*P* < 0.05) between SA-treated samples.

### The TGACGTCA motif in the *DLO1* promoter is a target site for TGA1 and for TGA2

The *DLO1* promoter contains only one of the typical TGA binding sites (TGACGTCA) and one A-box within 2000 bps upstream of the transcriptional start site. In order to investigate, whether representatives of both clades of TGAs might be accommodated at the promoter, we analyzed the effect of transiently expressed TGA1 and TGA2 on *DLO1* promoter activity. The *DLO1* regulatory region from −1777 bps (with respect to the transcriptional start site) to the ATG start codon was fused to the open reading frame of the firefly *luciferase* gene (*fLUC*), while *TGA1* and *TGA2* were expressed under the control of the *UBIQUITIN10* (*UBQ10*) promoter. Proteins were tagged at their C-terminal ends with a triple HA and a streptavidin tag. An “empty” vector expressing only the triple HA tag under the control of the *UBQ10* promoter was used as a control of background promoter activity and adjustment of equal amounts of DNA in the transfection mixture. *Renilla luciferase* (*rLUC*) served to normalize for transfection efficiency.

In order to avoid background activation by endogenous clade-I and clade-II TGAs, we used protoplasts of the *tga1 tga2 tga4 tga5 tga6* mutant that was previously obtained by crossing the respective genotypes. In this assay, only TGA1, but not TGA2 activated the promoter. Co-expression of TGA1 with TGA2 slightly enhanced promoter activity (Fig. 5a and Fig. S2). Since TGA2 - in contrast to TGA1 - does not contain an extended N-terminal domain with transactivation capacities, a potential binding of TGA2 to the promoter might have been missed in this assay. SA treatment, which leads to association of the transcriptional co-activator NPR1 with clade-II TGAs in differentiated leaf cells, did not specifically increase TGA-enhanced expression of the *Prom_DLO1_:fLUC* construct in protoplasts (Fig. S2). Thus, the SA signal transduction chain does not operate in the same manner in protoplasts as in mesophyll cells. To compensate for this, the coding regions of *TGA1* and *TGA2* were fused C-terminally to the activation domain of the *Herpes simplex* viral protein (VP) 16. Strong trans-activation of the *DLO1* promoter by TGA1-VP and TGA2-VP (Fig. 5b) allowed to address the importance of the TGA binding sites. Activation was abolished when the TGACGTCA motif at position −72 bps relative to the transcriptional start site was mutated while mutation of the A-box at position −1731 bps did not affect the responsiveness of the promoter to both TGAs. It is concluded that the single TGA binding site of the *DLO1* promoter is the only site to recruit either clade-I or clade-II TGAs.

**Fig. 5.**
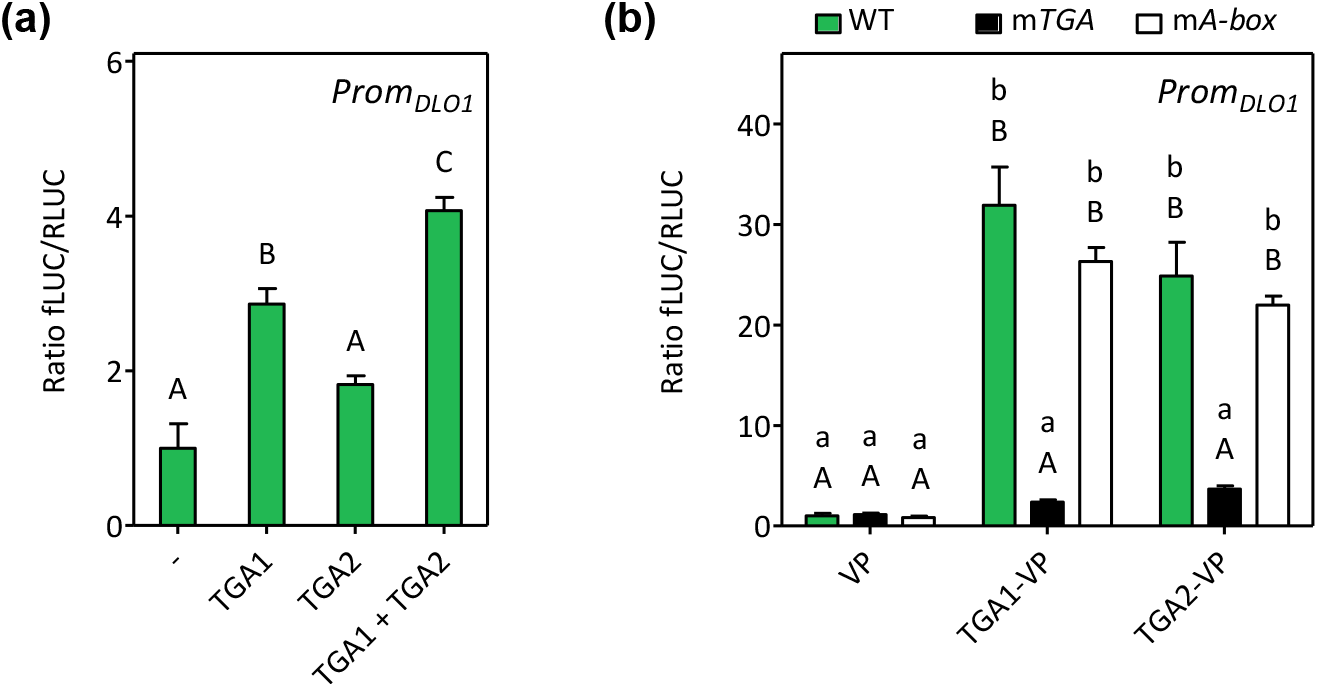
TGA1 and TGA2 both recognize the single *TGACGTCA* motif in the *DLO1* promoter. (a) Relative luciferase (LUC) activities yielded by the *DLO1* promoter as a function of co-expressed TGA1, TGA2 or TGA1 together with TGA2. The *Prom_DLO1_:fLUC* reporter plasmid was transformed into Arabidopsis *tga1 tga2 tga4 tga5 tga6* mesophyll protoplasts with either an “empty” effector plasmid or effector plasmids encoding TGA1 and/or TGA2 under the control of the *UBQ10* promoter. (b) Relative LUC activities yielded by the TGA-VP-activated *DLO1* promoter as a function of the presence of the *TGACGTCA* element or the A-box. Reporter plasmids were transformed into Arabidopsis Col-0 protoplasts with either an effector plasmid encoding TGA1 or TGA2 fused to the activation domain of viral protein 16 (VP) under the control of the *UBQ10* promoter, respectively, or a control plasmid encoding non-fused VP. WT – wild-type *Prom_DLO1_:fLUC* reporter sequence, *mTGA* – reporter plasmid with mutated *TGACGTCA* motif, *mA-box* – reporter plasmid with mutated *A-box*. Firefly LUC activities were normalized to *Renilla* LUC activities. LUC activity obtained from the wild-type *DLO1* promoter in the presence of the respective control vector plasmids was set to 1. Values are means of four independently transfected batches of protoplasts (+/− SEM). Different letters indicate significant differences at *P* < 0.05 (one-way ANOVA followed by Tukey’s post hoc test) for the various transfections in (a). In (b), statistical analysis was done using two-way ANOVA followed by Bonferroni’s post hoc test: lowercase letters indicate significant differences (P < 0.05) between various reporter constructs combined with the same type of effector plasmid; uppercase letters indicate significant differences (P < 0.05) between different effectors combined with the same reporter variant.

### *DLO1* and *BGL2* expression depends on SARD1/CBP60g

On the one hand, the requirement of clade-I and clade-II TGAs for expression of a marker gene with only one TGACG motif in the promoter might be explained by a heterodimer being active at this promoter. This scenario seems feasible since heterodimerization between *in vitro* co-translated tobacco TGA1a and TGA2 has been shown before (Niggeweg *et al.*, 2000). Alternatively, they might bind as homodimers not only at the final target gene, but also at genes encoding for regulators acting upstream in the SA-dependent signaling cascade. This hypothesis fits to the expression pattern of *SARD1*, which is regulated by the well-established NPR1-TGA2/TGA5/TGA6 module, but not by TGA1/TGA4 (Fig. 1). As shown in Fig. 6, expression of *DLO1* and *BGL2* was strongly reduced in the *sard1 cbp60g* double mutant, which lacks not only SARD1 but also the related and sometimes redundantly acting factor CBP60g (CALCIUM BINDING PROTEIN 60g (Wang *et al.*, 2011)). This effect was less pronounced for *PR1* and absent for *GSTF6* expression. At least for *DLO1* and *BGL2*, which are more affected by TGA1/TGA4 than *PR1* and *GSTF6*, the concept of indirect regulation by clade-II TGA-activated SARD1 and direct regulation by clade-I TGAs seems plausible. Consistently, both promoters contain *SARD1* binding sites. No *SARD1* binding sites were found in the other two promoters.

**Fig. 6.**
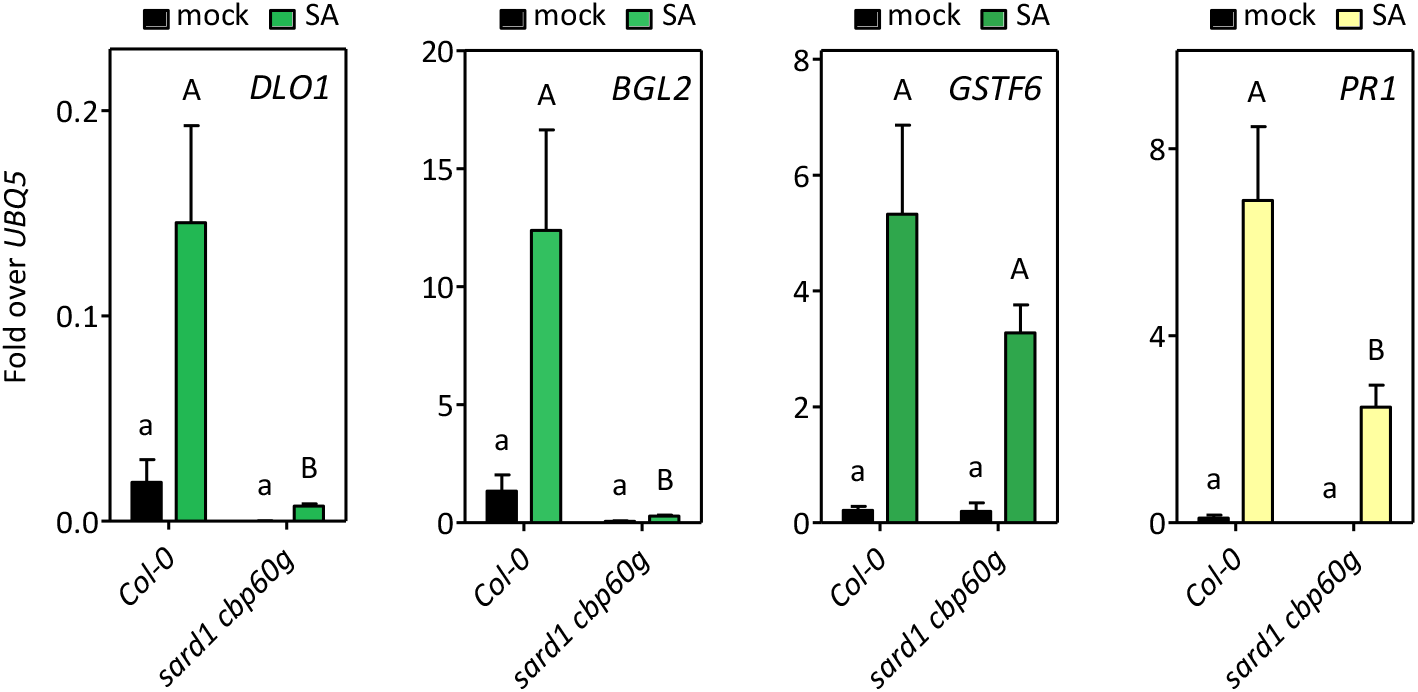
*DLO1* and *BGL2* expression strongly depends on SARD1/CBP60g. qRT-PCR analysis of transcript levels of four representative TGA1/TGA4-dependent genes in wild-type (Col-0) and *sard1 cbp60g* plants. Four-week-old plants were sprayed either with water (mock) or 1 mM salicylic acid (SA) and further incubated for 8 h. The experiment is part of the experiment shown in Figure 4 with Col-0 used as a common control. Transcript levels were normalized to transcript levels of *UBQ5*. Bars represent the average ± SEM of four to five plants of each genotype. Statistical analysis was performed using unpaired Student’s t-test (two-tailed) for mock- and SA-treated samples separately. Lowercase letters indicate significant differences (*P* < 0.05) between mock-treated samples; uppercase letters indicate significant differences (*P* < 0.05) between SA-treated samples.

### Mutation of the redox-active cysteines does not alter the expression pattern of selected marker genes in SA-treated leaves

As mentioned above, a disulfide bridge between C260 and C266 was detected in 50% of the TGA1 proteins in untreated tissue, while 100% of the TGA1 pool is reduced in SA-treated tissue resulting in a larger amount of TGA1 being able to interact with NPR1, which in turn leads to increased DNA binding (Despres *et al.*, 2003). Having identified SA-induced TGA1/TGA4-dependent genes we were now able address the importance of these “SA-switchable” cysteines. Since the two flanking cysteines C172 and C287 are also prone to redox modifications (Lindermayr *et al.*, 2010), all four cysteines were mutated (C172N C260N C266S C287S). The first three cysteines were changed into residues found in TGA2 at the corresponding positions, while the last cysteine was changed to serine, which is found at the corresponding positions in TGA3, TGA4, TGA7 and TGA9. Mutations were introduced into a genomic clone that consisted of 2671 bps upstream of the translational start site, exons and introns and 217 bps downstream of the transcribed region. These lines, along with a transgenic line transformed with the empty vector, were treated with water or with SA and the expression of TGA1/TGA4-dependent marker genes was monitored. As observed before, expression of *DLO1*, *BGL2, GSTF6* and *PR1* was reduced in the absence of TGA1/TGA4. Both, the wild-type and the mutated TGA1 protein rescued SA-induced expression of the four TGA1/TGA4-dependent marker genes to the same degree (Fig. 7a). Also, background levels were not differentially affected in plants expressing either TGA1 or mutated TGA1 (for background *SARD1* transcript levels, see Fig. S3). Although the proteins accumulated to higher levels than endogenous TGA1 in the untransformed Col-0 wild-type plants (Fig. 7b), only partial complementation was observed for *GSTF6* and *PR1*. This might be due to the absence of TGA4, to the N-terminal HA-tag that was introduced upstream of the ATG start codon, or the history of the untransformed Col-0 seeds. Since the non-mutated and the mutated proteins were equally effective, it is concluded that the SA-mediated redox switch in TGA1 does not contribute to the proper expression of TGA1/TGA4-dependent target genes at eight hours after mock- or SA treatment.

**Fig. 7.**
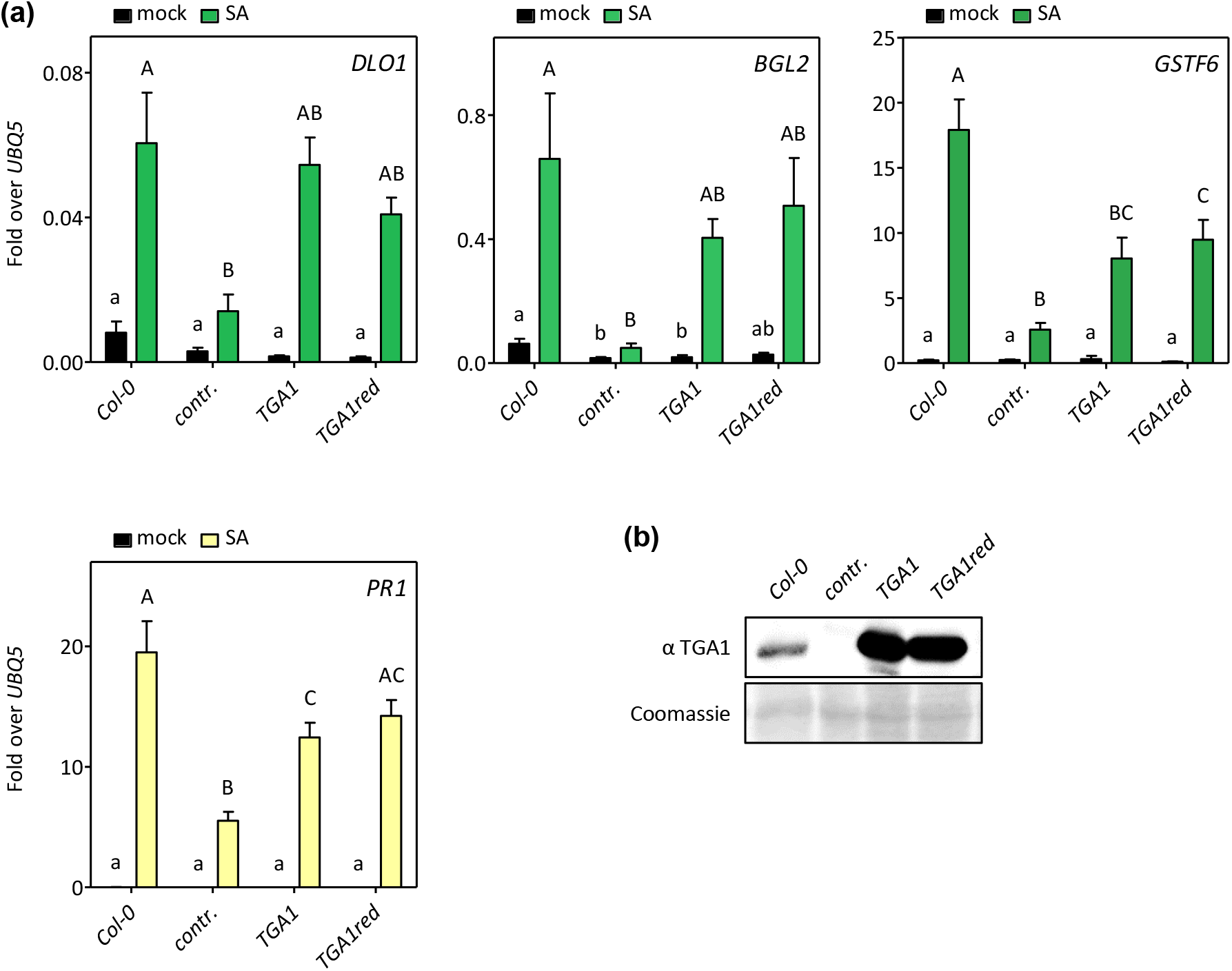
Cysteines in TGA1 are not important for wildtype-like expression of salicylic acid (SA)-induced target genes. (a) qRT-PCR analysis of transcript levels of four TGA1/TGA4-dependent genes after SA treatment of wild-type (Col-0) and *tga1 tga4* plants complemented either with a control vector *(contr.)*, a wildtype *TGA1* genomic construct (*TGA1*) or a mutated *TGA1* genomic construct carrying mutations in four critical cysteine residues (*TGA1red*). Four-week-old plants were sprayed either with water (mock) or 1 mM SA at 1 h after the subjective dawn and further incubated for 8 h. Transcript levels were normalized to transcript levels of *UBQ5*. Bars represent the average ± SEM of four to six plants of each genotype. Statistical analysis was performed using one-way ANOVA followed by Tukey’s post hoc test for mock- and SA-treated samples separately. Lowercase letters indicate significant differences (*P* < 0.05) between mock-treated samples; uppercase letters indicate significant differences (*P* < 0.05) between SA-treated samples. (b) Western blot analysis of protein extracts obtained from roots of the different plant genotypes as indicated in (a). TGA1 protein levels were detected using an anti-TGA1 antibody. Coomassie blue staining served as a loading control.

### TGA1/TGA4 might act in concert with NPR1 when regulating *SARD1* after pathogen infection

Since *SARD1* expression is modulated by TGA1/TGA4 upon pathogen infection (Sun *et al.*, 2018), we questioned whether under these conditions TGA1/TGA4 might act in concert with NPR1 and SA. We tested *SARD1* expression eight hours after infection with *Psm* in the *tga1 tga4*, *tga2 tga5 tga6, npr1* and the *sid2* mutants (Fig. 8a). Indeed, *Psm*-induced expression of *SARD1* was reduced to approximately the same levels in *sid2*, *npr1* and *tga1 tga4* (Fig. 8a). Importantly, expression levels were unaffected in *tga2 tga5 tga6*. This suggests that an SA-mediated feedforward loop using NPR1 and TGA1/TGA4 enhances *SARD1* expression independently from TGA2/TGA5/TGA6. Likewise, in *Psm*-infected SAR leaves, NPR1 and TGA1/TGA4, but not TGA2/TGA5/TGA6, were important for *SARD1* expression (Fig. 8b).

**Fig. 8.**
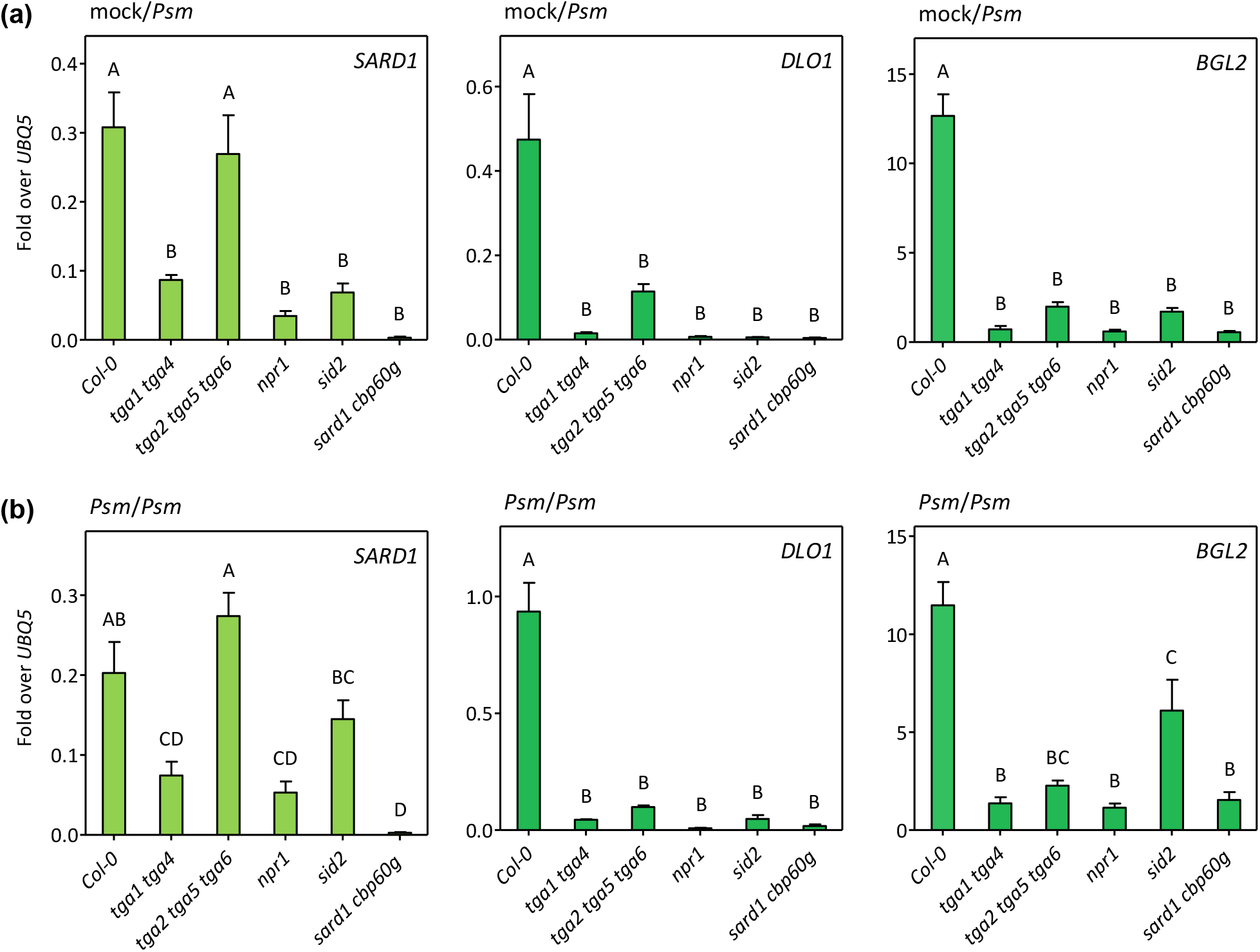
Clade-II TGAs are not important for *SARD1* but for *DLO1* and *BGL2* expression after infection with *Pseudomonas syringae* pv. *maculicola* ES4356 (*Psm*). qRT-PCR analysis of transcript levels of *SARD1* and *DLO1* in wild-type (Col-0), *tga1 tga4, tga2 tga5 tga6, npr1, sid2* and *sard1 cpb60g* plants. Three leaves of five-week-old plants were either MgCl_2_ (mock)-infiltrated (a) or infiltrated with *Psm* (*O*D_600_ of 0.005) (b) at 1 h after the subjective dawn. Two days later, three younger upper leaves were infiltrated with *Psm* (OD_600_ of 0.005). After 8 hours, these were harvested for RNA extraction. Transcript levels were normalized to transcript levels of *UBQ5*. Bars represent the average ± SEM of three to four plants of each treatment. Statistical analysis was performed using one-way ANOVA followed by Tukey’s post hoc test. Letters indicate significant differences (*P* < 0.05) between the different genotypes.

As observed in SA-treated tissue, SARD1/CBP60g, NPR1 and clade-I and clade-II TGAs were important for *DLO1* and *BGL2* expression in *Psm*-infected leaves, independent of whether they had been pre-treated with *Psm* or with MgCl_2_. Since clade-II TGAs were not required for *SARD1* expression and thus did not influence *ICS1* transcript levels (Fig. S4), we consider it likely that SA levels were not reduced in *tga2 tga5 tga6*. It is concluded that these factors activate *DLO1* directly, while the effect of the clade-I TGAs and NPR1 can be partially explained by reduced SA levels due to reduced *SARD1* and *ICS1* expression.

*BGL2* transcript levels followed a similar trend. However, it has to be noted that in *Psm*-infected SAR leaves, *SARD1* and *BGL2* were not as stringently dependent on SA as in *Psm*-infected leaves from plants that had be pre-treated with MgCl_2_. NPR1 remained to be necessary even when SA levels were not as critical for induction. This suggests that a signaling molecule different from SA can activate the NPR1/TGA1/TGA4 regulatory module in *Psm*-infected SAR leaves. A similar phenomenon has been observed very recently in the auto-immune mutant *camta123* showing that *SARD1* transcript levels were higher in *sid2* as compared to *npr1* (Kim *et al.*, 2019).

### Mutation of the redox-active cysteines of TGA1 does not change the expression pattern of selected marker genes after pathogen infection

Having identified *SARD1* as an SA/NPR1/TGA1/TGA4-dependent target gene after pathogen infection we analysed its expression in the complementation lines (Fig. 9). As expected, *Psm*-induced *SARD1* expression was reduced in *tga1 tga4* plants transformed with the “empty vector”. Importantly, both *TGA1* constructs (i.e. *TGA1* and *TGA1red*) complemented the phenotype to the same extent. Elevated expression after *Psm* pre-infections as compared to mock pre-treatments and the contribution of TGA1/TGA4 to gene expression was more pronounced for *DLO1* and *BGL2.* Again, TGA1 lacking all four cysteines complemented the phenotype to a similar extent as the wildtype protein, supporting the notion that the lack of potential oxidative modifications does not alter the regulatory properties of the protein under these conditions.

**Fig. 9.**
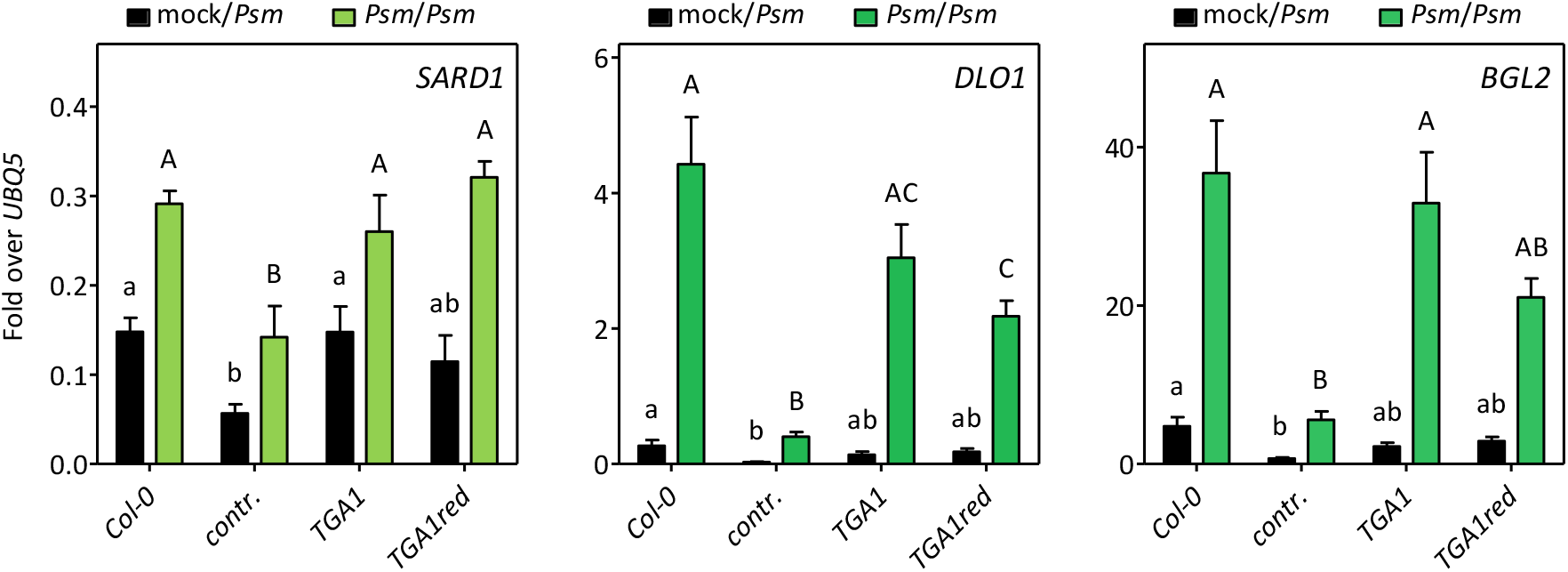
Cysteines in TGA1 are not important for wildtype-like expression of target genes in pathogen-infected leaves. qRT-PCR analysis of transcript levels of TGA1/TGA4 target genes in wild-type (Col-0) and *tga1 tga4* plants complemented either with a control vector (*contr.*), a wildtype *TGA1* genomic construct (*TGA1*) or a mutated *TGA1* genomic construct carrying mutations in four critical cysteine residues (*TGA1red*). Three leaves of five-week-old plants were MgCl_2_ (mock)- or *Psm*-infiltrated (OD_600_ of 0.005) at 1 h after the subjective dawn. Two days later, three younger upper leaves were infiltrated with *Psm* (OD_600_ of 0.005). After 8 hours, these were harvested for RNA extraction. Transcript levels were normalized to transcript levels of *UBQ5*. Bars represent the average ± SEM of five to six plants of each treatment. Statistical analysis was performed using one-way ANOVA followed by Tukey’s post hoc test for mock- and *Psm-*pretreated samples separately. Lowercase letters indicate significant differences (*P* < 0.05) between mock-pretreated samples; uppercase letters indicate significant differences (*P* < 0.05) between *Psm*-pretreated samples.

## DISCUSSION

Arabidopsis TGA transcription factors TGA1 and TGA4 interact with the SA-activated transcriptional co-activator NPR1 in a redox-dependent manner (Despres *et al.*, 2003; Lindermayr *et al.*, 2010). Here, using TGA1 mutants with point mutations in all four cysteines, we show that these cysteines do not play a role in SA- or pathogen-induced NPR1-dependent expression of TGA1/TGA4-regulated marker genes. We identified TGA1/TGA4 as a positive regulator of the SA catabolizing gene *DLO1*. Finally, we found that the relative influence of clade-I and clade-II TGAs on *SARD1* expression depends on whether plants are treated with SA or with *Psm*.

In order to address the functional importance of redox-modulated cysteines in TGA1, we first identified SA-induced TGA1/TGA4-dependent genes by RNAseq analysis. Since it was known that SA synthesis is controlled by TGA1/TGA4 (Sun *et al.*, 2018), we performed the analysis in the SA biosynthesis mutant *sid2*. This strategy guaranteed that genes affected in the SA-treated *sid2 tga1 tga4* mutant as compared to *sid2* would require TGA1/TGA4 downstream of SA, while any effects upstream of SA were excluded. Only 193 out of the 2090 genes that were higher expressed at eight hours after SA treatment as compared to mock treatment showed reduced expression in SA-treated *sid2 tga1 tga4* plants. It is likely that this number would be even lower in the wild-type background since we observed more fluctuations in the presence of endogenous amounts of SA. The low frequency might be due to the expression pattern of TGA1/TGA4, the promoters of which are mainly active in the vascular tissue (Song *et al.*, 2008; Wang *et al.*, 2019). This correlates well with the expression pattern of the two robustly regulated target genes: *DLO1* is expressed near the vascular tissue in *Hpa*-infected leaves (Zeilmaker *et al.*, 2015), while *BGL2* is expressed near the vascular tissue in SA-treated leaves (Spoel *et al.*, 2009). We assume that the other regulatory components influencing *DLO1* and *BGL2* expression (NPR1, clade-II TGAs, SARD1) are also present in this tissue.

Still, the discrepancy to previously published gene expression patterns of the SA-treated *tga1 tga4* mutant has to be pointed out. Shearer *et al*. performed a similar study by analyzing the transcriptomes of soil-grown *tga1 tga4* and *npr1* mutants after one and eight hours after SA treatment (Shearer *et al.*, 2012). *DLO1*, *BGL2* and *PR1*, which were robustly less expressed in *tga1 tga4* in this study, showed increased basal levels in untreated *tga1 tga4*, while expression values were similar to wild-type levels after SA treatment. Likewise, Lindermayr et al. (2010) reported enhanced *PR1* transcript levels in the *tga1 tga4* mutant. We did not detect increased transcript levels of our marker genes in the *tga1 tga4* mutant after mock treatment. In the presence of SA, expression levels were clearly lower in the *tga1 tga4* mutant in our hands. Different growth conditions might affect how TGA1/TGA4 contribute to the SA signaling cascade.

Whatever the reason for these differences is, we were able to identify SA-induced NPR1-dependent genes that required TGA1/TGA4 for maximal expression. Due to the limited expression domain of TGA1/TGA4 we failed to prove direct binding to e.g the promoters of *DLO1* or *BGL2* by chromatin immunoprecipitation (ChIP) experiments. Similar problems were encountered before: binding of TGA1/TGA4 to the *SARD1* promoter was only shown in protoplasts ectopically expressing *TGA1* under the *Cauliflower Mosaic Virus (CaMV) 35S* promoter (Sun *et al.*, 2018), while binding of clade-II TGAs could be documented by ChIP in wild-type plants (Ding *et al.*, 2018).

To answer our primary research question, whether the redox-modulated NPR1-dependent DNA-binding activity of TGA1 influences the expression of SA-dependent target genes, we had to make sure that expression of the identified target genes are regulated by the interplay between SA, TGA1 and NPR1. However, the analysis was complicated since SA-induced expression of all four tested TGA1/TGA4-dependent target genes also depended on clade-II TGAs, which can recognize the same binding site as TGA1/TGA4. Given the fact that at least the *DLO1* promoter contains only one TGA binding site, we postulate for SA-treated tissues that SA activates NPR1 to stimulate expression of *SARD1* in concert with clade-II TGAs. Subsequently, SARD1 acts at the *DLO1* and the *BGL2* promoters, the expression of which is further enhanced by TGA1/TGA4. Thus, in SA-treated tissues, we could not clearly establish that *DLO1* or *BGL2* are regulated by a mechanism that is controlled by TGA1/TGA4 interacting with NPR1.

Interestingly, the functions of clade-I and clade-II TGAs in the SA-dependent regulatory network were changed in *Psm*-infected leaves. Here, the *SARD1* promoter remained to be responsive to NPR1, but was regulated by TGA1/TGA4 while TGA2/TGA5/TGA6 became dispensable. Thus, in this tissue, at least *SARD1* was the candidate gene we were looking for to address the functional importance of the redox-regulated cysteines. However, the redox-regulated cysteines did not play a role for *SARD1* expression, at least after eight hours after pathogen infection of naïve or SAR leaves. Under these conditions, endogenous SA levels might have already led to full reduction of the wild-type protein.

According to previously published data, interfering with the internal disulfide bridge formation should lead to a protein that constitutively interacts with NPR1 and subsequently binds to DNA with a higher affinity (Despres *et al.*, 2003; Lindermayr *et al.*, 2010). Thus, higher background activity of at least *SARD1* and thus its downstream genes might have been the expected consequence of the mutations. This was not observed, most likely due to other inhibitory mechanisms including the repressive effects of NPR3 and NPR4 (Ding *et al.*, 2018). A phenotype might be expected if oxidation and thus inactivation of TGA1 would happen under certain conditions. Our complementation lines in combination with the TGA1/TGA4-dependent marker genes might provide useful tools to analyse whether potential antagonistic effects of e.g. reactive oxygen species-generating abiotic stresses that interfere with the SA pathway are less pronounced in plants expressing a mutant TGA1 protein that cannot be oxidized.

## Supporting Information

Supporting Figures, Supporting Tables, Supporting Methods, Supporting Notes

## Acknowledgements

J.B. and A.N. were funded by the German Research Foundation DFG (IRTG 2172 “PRoTECT”), K.T. was funded by a Dorothea Schlözer fellowship granted by the University of Göttingen. We thank Anna Herrmann for excellent technical assistance. We also thank the Transcriptome and Genome Analysis Laboratory (TAL) of the University Medical Center Göttingen (UMG) for performing the RNAseq analysis.

## Author contribution

J.B. designed, acquired, analyzed, and interpreted most of the data, K.T. generated the *TGA* complementation lines, A.N. contributed to the data obtained with *Psm-* infected plants, C.T. performed the bioinformatic analysis of the RNAseq data, C.G. and C.T. participated in the design of the study and wrote the manuscript. All authors read and approved the final manuscript.

## Supporting Information

Additional Supporting Information may be found in the online version of this article.

**Fig. S1** Principal component analysis of the normalized transcriptome data obtained from RNAseq analysis.

**Fig. S2** Salicylic acid (SA) treatment does not increase TGA-dependent activation of the *DLO1* promoter in mesophyll protoplasts.

**Fig. S3** TGA1 with mutated cysteines does not lead to increased basal *SARD1* transcript levels.

**Fig. S4** Clade-II TGAs are not important for *ICS1* expression after infection with *Pseudomonas syringae* pv. *maculicola* ES4356 (*Psm*).

**Table S1** Primers used for qRT-PCR.

**Table S2** Expression Data of 2090 salicylic acid-inducible genes.

**Table S3** Fold change in selected transcripts as identified by RNAseq analysis of four-week old Arabidopsis *sid2* and *sid2 tga1 tga4* treated with water (mock) or 1 mM salicylic acid (SA) for 8 hours.

**Methods S1** Detailed description of methods.

**Notes S1** Maps and sequences of plasmids used in this work.

