## Supplementary material for "Redox-active cysteines in TGACG-BINDING FACTOR 1 (TGA1) do not play a role in salicylic acid- or pathogen-induced expression of TGA1-regulated target genes in *Arabidopsis thaliana*": Figures and Methods

***New Phytologist* Supporting Information**

The following Supporting Information is available for this article:

**Fig. S1** Principal component analysis of the normalized transcriptome data obtained from RNAseq analysis.

**Fig. S2** Salicylic acid (SA) treatment does not increase TGA-dependent activation of the *DLO1* promoter in mesophyll protoplasts.

**Fig. S3** TGA1 with mutated cysteines does not lead to increased basal *SARD1* transcript levels.

**Fig. S4** Clade-II TGAs are not important for *ICS1* expression after infection with *Pseudomonas syringae* pv. *maculicola* ES4356 (*Psm*).

**Table S1** Primers used for qRT-PCR.

**Table S2** Expression Data of 2090 salicylic acid-inducible genes.

**Table S3** Fold change in selected transcripts as identified by RNAseq analysis of four-week old Arabidopsis *sid2* and *sid2 tga1 tga4* treated with water (mock) or 1 mM salicylic acid (SA) for 8 hours.

**Methods S1** Detailed description of methods.

**Notes S1** Maps and sequences of plasmids used in this work.

**Fig. S1** Principal component analysis of the normalized transcriptome data obtained from RNAseq analysis.

Symbols represent four independent biological replicates of *sid2* and *sid2 tga1 tga4* at 8 hours after salicylic acid (SA) or control (water) treatments.

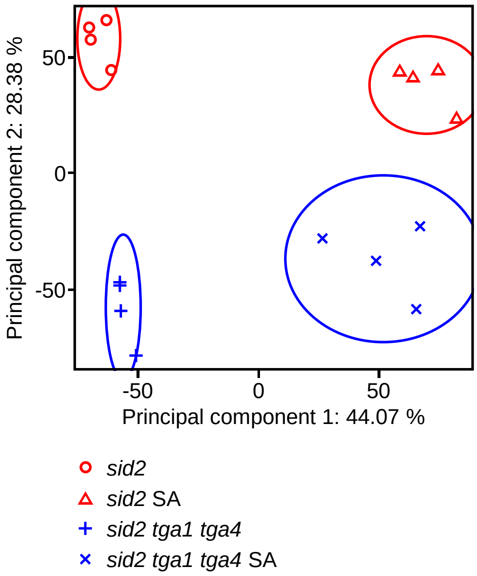

**Fig. S2** Salicylic acid (SA) treatment does not increase TGA-dependent activation of the *DLO1* promoter in mesophyll protoplasts.

Relative luciferase (LUC) activities yielded by the *DLO1* promoter as a function of co-expressed TGA1, TGA2 or TGA1 together with TGA2. The *Prom*$_{DLO1}$*:fLUC* reporter plasmid was transformed into Arabidopsis *tga1 tga2 tga4 tga5 tga6* mesophyll protoplasts with either an empty effector plasmid or an effector plasmid encoding TGA1 or TGA2 under the control of the *UBQ10* promoter. After transfection, protoplasts were incubated overnight in WI buffer with or without (mock) 2 µM SA. Firefly LUC activities were normalized to *Renilla* LUC activities. LUC activity obtained from the *DLO1* promoter in the presence of an "empty" effector plasmid was set to 1. Values are means of three independently transfected batches of protoplasts (+/- SEM). Lowercase letters indicate significant differences ($P < 0.05$) between the treatments performed with the same type of transfection; uppercase letters indicate significant differences ($P < 0.05$) between various transfections subjected to the same treatment. Statistical analysis was done using two-way ANOVA followed by Bonferroni's post hoc test.

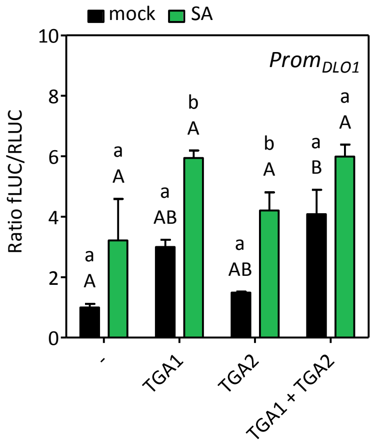

**Fig. S3** TGA1 with mutated cysteines does not lead to increased basal *SARD1* transcript levels.

qRT-PCR analysis of *SARD1* transcript levels in wild-type (Col-0) and *tga1 tga4* plants complemented either with a control vector (*contr.*), a wildtype *TGA1* genomic construct (*TGA1*) or a mutated *TGA1* genomic construct carrying mutations in four critical cysteine residues (*TGA1red*). Four-week-old plants were sprayed either with water (mock) or 1 mM SA at 1 h after the subjective dawn and further incubated for 8 h. Transcript levels were normalized to transcript levels of *UBQ5*. Bars represent the average ± SEM of four to five plants of each genotype. Statistical analysis was performed using one-way ANOVA followed by Tukey's post hoc test for mock- and SA-treated samples separately. Lowercase letters indicate significant differences ($P < 0.05$) between mock-treated samples; uppercase letters indicate significant differences ($P < 0.05$) between SA-treated samples.

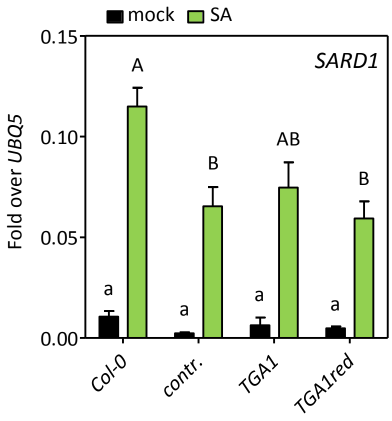

**Fig. S4** Clade-II TGAs are not important for *ICS1* expression after infection with *Pseudomonas syringae* pv. *maculicola* ES4356 (*Psm*).

qRT-PCR analysis of *ICS1* transcript levels in wild-type (Col-0) and *tga2 tga5 tga6* plants. Three leaves of five-week-old plants were either $MgCl_2$ (mock)-infiltrated or infiltrated with *Psm* ($OD_{600}$ of 0.005) at 1 h after the subjective dawn. After two days, three younger upper leaves were infiltrated with *Psm* ($OD_{600}$ of 0.005). After 8 hours, these were harvested for RNA extraction. The same samples were analysed in Fig. 8. Transcript levels were normalized to transcript levels of *UBQ5*. Bars represent the average ± SEM of three to four plants of each genotype. Unpaired Student's t-test (two-tailed) revealed no significant differences between the genotypes.

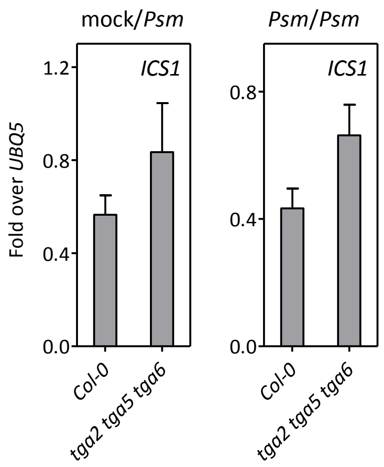

**Table S1** Primers used for qRT-PCR.

| Primer | Sequences (5'-3') |
|---|---|
| ATAF1 | QuantiTect QT00866439 (Qiagen, Hilden, Germany) |
| BGL2 | QuantiTect QT00793730 (Qiagen, Hilden, Germany) |
| DLO1 qRT For | AATATCGGCGACCAAATGC |
| DLO1 qRT Rev | CGCTCGTTCTCGGTGTTTAC |
| GSTF6 | QuantiTect QT00868147 (Qiagen, Hilden, Germany) |
| ICS1 | QuantiTect QT00893473 (Qiagen, Hilden, Germany) |
| PR1 qRT For | CTGACTTTCTCCAAACAACTTG |
| PR1 qRT Rev | GCGAGAAGGCTAACTACAACTAC |
| SARD1 qRT For | TCAAGGCGTTGTGGTTTGTG |
| SARD1 qRT Rev | CGTCAACGACGGTATGTTTC |
| UBQ5 qRT For | GACGCTTCATCTCGTCC |
| UBQ5 qRT Rev | GTAAACGTAGGTGAGTCCA |
| WRKY6 | QuantiTect QT00884884 (Qiagen, Hilden, Germany) |

**Table S3** Fold change in selected transcripts as identified by RNAseq analysis of four-week old Arabidopsis *sid2* and *sid2 tga1 tga4* treated with water (mock) or 1 mM salicylic acid (SA) for 8 hours.

| | **Fold change** | | |
|---|---|---|---|
| | **SA / mock (*sid2*)** | **SA / mock (*sid2 tga1 tga4*)** | ***sid2* / *sid2 tga1 tga4* (SA)** |
| ***DLO1*** | 30.4 | 31.6 | 7.0 |
| ***BGL2*** | 4.3 | 1.4 | 8.2 |
| ***GSTF6*** | 52.2 | 4.3 | 3.3 |
| ***PR1*** | 825.4 | 555.9 | 1.9 |
| ***ATAF1*** | 3.4 | 1.9 | 0.3 |
| ***WRKY6*** | 3.8 | 1.2 | 0.6 |

**Methods S1.** Detailed description of methods.

#### Construction of recombinant plasmids

The genomic *TGA1* complementation construct with an N-terminal HA tag-encoding sequence was cut out from pG0229HAgTGA1 (Li et al., 2019) via *Acc*65I/*Stu*I and ligated into the binary vector pBGW (http://www.psb.ugent.be/gateway/) digested with *Acc*65I/*Eco*RV. The map of the resulting pB-HAgTGA1 is shown in Supporting Information Notes S1. The C172N and C287S cysteine mutations were introduced through overlapping PCR using the plasmid pG0229HAgTGA1(C260N/C266S) (Li et al., 2019) as template DNA. A comparison between wild-type *TGA1* and mutated DNA regions in pB-HAgTGA1(C172N/C260N/C266S/C287S) is given in Supporting Information Notes S1. The control plasmid pB-HA was obtained by deletion of the *TGA1* coding region (start codon until last amino acid codon, see Supporting Information Notes S1). All constructs were confirmed by sequencing and transformed into *tga1 tga4* plants.

Generation of plasmids for transient transformation of Arabidopsis mesophyll protoplasts was done using GATEWAY technology (Invitrogen, Karlsruhe, Germany). For expression of C-terminally fused TGAs, pPZP200-based binary vectors (Hajdukiewicz et al., 1994) were modified by various cloning steps yielding pUBQ10GW3HAstrepII7 and pUBQ10GWVP7 (Supporting Information Notes S1). *TGA1* (genomic sequence with introns)- and *TGA2* (cDNA)-encoding regions were amplified as PCR fragments from the particular start codon to the last amino acid codon, respectively, using primers that add GATEWAY recombination sites. Fragments were inserted into pDONR207 and subsequently recombined into the respective destination vectors. Control plasmids contain coding regions for 3xHA-tag or 3xHA-tag fused to the VP16 activation domain, respectively. The reporter plasmid pB-DLO1promLUC (see Supporting Information Notes S1) was constructed by recombining a *DLO1* promoter PCR fragment (-1777 to +77 relative to transcriptional start site) via pDONR207 into the destination vector pBGWL7 (http://www.psb.ugent.be/gateway/). Mutations of the *TGACGTCA* motif and the *A-box* were introduced via overlapping PCR, respectively (nucleotide exchanges are shown in Supporting Information Notes S1).

#### Transcriptome analyses

For each sample, three leaves of five individual plants were collected. The experiment was repeated four times with batches of independently grown plants.

RNA was extracted by the Trizol method and RNA quality control was performed using the AGILENT BIOANALYZER 2100. Single-end 50 bp raw reads from mRNA sequencing were generated by the Illumina HiSeq 2000 platform. Sequence images were transformed with the Illumina BaseCaller software to BCL files, which were demultiplexed to FASTQ files with the bcl2fastq v2.17.1.14 Conversion Software (Illumina). Subsequently, reads were mapped to the genome reference sequence of *Arabidopsis thaliana* (TAIR10 release-38, http://plants.ensembl.org/Arabidopsis_thaliana/Info/Index) using the STAR aligner version 2.5 alignment software (Dobin et al., 2013) allowing for two mismatches within 50 bases. Final read counting was performed with featureCounts version 1.4.5-p1 (Liao et al., 2013).

For statistical assessment of differential gene expression, read counts were imported into the *R*obi*NA* 1.2.4_buid656 application (Lohse et al., 2012) and analyzed by the DESeq method to obtain log2 fold change and adjusted *P* values (Benjamini-Hochberg-corrected).

Analysis of *cis* element enrichment was done using the Cluster Analysis Real Randomization algorithm incorporated into the Motif Mapper Version 5.2.4.0 (Berendzen et al., 2012) to define significant distribution alterations compared to 1000 randomly composed, equally sized, reference promoter datasets as described in Zander et al. (2014).

#### Quantitative reverse transcription (qRT)-PCR

RNA extraction and qRT-PCR analyses were performed as described (Fode et al., 2008). Calculations were done according to the $2^{-\Delta C_T}$ method (Livak & Schmittgen, 2001) using the *UBQ5* transcript as a reference (Kesarwani et al., 2007). Primers serving to amplify and quantify transcript levels are listed in Table S1.

#### Transient expression analysis in Arabidopsis protoplasts

Protoplasts assays were carried out as described (Yoo et al., 2007). 5 µg of effector plasmids harboring sequences for HA-tagged *TGA1* and *TGA2* or *TGA1-VP* and *TGA2-VP* under the control of the *Arabidopsis thaliana UBQ10* promoter and 5 µg of the reporter plasmid containing the firefly *LUCIFERASE* coding region (*fLUC*) under the control of the *DLO1* promoter and its variants were co-transfected. To normalize for the experimental variability, 1 µg of the plasmid pUBQ10rLUC encoding the *Renilla LUCIFERASE* (*rLUC*) gene under the control of the *UBQ10* promoter, was added to each sample. After transformation, protoplast were incubated for 16 hours in WI buffer (0.5 M mannitol, 4 mM 2-(N-morpholino)ethanesulfonic acid, 20 mM KCl) with or without 2 µM salicylic acid. LUC activities were measured using the Dual-Luciferase® Reporter Assay System (Promega, Mannheim, Germany) and the Centro XS3 LB 960 Microplate Luminometer (Berthold Technologies, Wildbad, Germany).

#### Western blot analysis

TGA1 protein levels in transgenic plants containing the genomic *TGA1* complementation constructs in the *tga1 tga4* mutant background were detected by Western blot analysis. Protein extracts were prepared by mixing 100 mg ground frozen root tissue with 150 µL of extraction buffer (4 M urea, 16.6 % glycerol, 5 % SDS, 0.5 % 2-mercaptoethanol). Protein concentrations were determined using the Pierce 660nm Protein Assay Reagent in combination with the Ionic Detergent Compatibility Reagent (Thermo Fisher Scientific Inc., Rockford, IL USA). 10 µg of root proteins were loaded on a 12 % SDS polyacrylamide gel and transferred to a polyvinylidene difluoride membrane by semi-dry electroblotting. Proteins were detected using an anti-TGA1 antibody (#AS16 3208, Agrisera, Vännäs, Sweden) and the SuperSignal™West Femto Maximum Sensitivity Substrate kit (Thermo Fisher Scientific Inc., Rockford, IL USA).

**Accession numbers**

Sequence data from this article can be found in the Arabidopsis Genome Initiative or GenBank/EMBL databases under the following accession numbers:

ATAF1 - At1g01720; BGL2 - At3g57260; DLO1 - At4g10500; GSTF6 - At1g02930; ICS1 - AT1G74710; PR1 - At2g14610; SARD1 - At1g73805; UBQ5 - At3g62250; WRKY6 - At1g62300.

**Berendzen KW, Weiste C, Wanke D, Kilian J, Harter K, Droge-Laser W. 2012.** Bioinformatic *cis*-element analyses performed in Arabidopsis and rice disclose bZIP- and MYB-related binding sites as potential AuxRE-coupling elements in auxin-mediated transcription. *BMC Plant Biol* **12**: 125.

**Dobin A, Davis CA, Schlesinger F, Drenkow J, Zaleski C, Jha S, Batut P, Chaisson M, Gingeras TR. 2013.** STAR: ultrafast universal RNA-seq aligner. *Bioinformatics* **29**(1): 15-21.

**Fode B, Siemsen T, Thurow C, Weigel R, Gatz C. 2008.** The Arabidopsis GRAS protein SCL14 interacts with class II TGA transcription factors and is essential for the activation of stress-inducible promoters. *Plant Cell* **20**(11): 3122-3135.

**Hajdukiewicz P, Svab Z, Maliga P. 1994.** The small, versatile pPZP family of Agrobacterium binary vectors for plant transformation. *Plant Mol Biol* **25**(6): 989-994.

**Kesarwani M, Yoo J, Dong X. 2007.** Genetic interactions of TGA transcription factors in the regulation of pathogenesis-related genes and disease resistance in Arabidopsis. *Plant Physiol* **144**(1): 336-346.

**Li N, Muthreich M, Huang LJ, Thurow C, Sun T, Zhang Y, Gatz C. 2019.** TGACG-BINDING FACTORs (TGAs) and TGA-interacting CC-type glutaredoxins modulate hyponastic growth in Arabidopsis thaliana. *New Phytol.* **121**(4): 1906-1918.

**Liao Y, Smyth GK, Shi W. 2013.** featureCounts: an efficient general purpose program for assigning sequence reads to genomic features. *Bioinformatics* **30**(7): 923-930.

**Livak KJ, Schmittgen TD. 2001.** Analysis of relative gene expression data using real-time quantitative PCR and the 2(-Delta Delta C(T)) Method. *Methods* **25**(4): 402-408.

**Lohse M, Bolger AM, Nagel A, Fernie AR, Lunn JE, Stitt M, Usadel B. 2012.** *R*obi*NA*: a user-friendly, integrated software solution for RNA-Seq-based transcriptomics. *Nuleic Acids Res* **40**(Web Server issue):W622-7.

**Yoo SD, Cho YH, Sheen J. 2007.** Arabidopsis mesophyll protoplasts: a versatile cell system for transient gene expression analysis. *Nat Protoc* **2**(7): 1565-1572.

**Zander M, Thurow C, Gatz C. 2014.** TGA transcription factors activate the salicylic acid-suppressible branch of the ethylene-induced defense program by regulating *ORA59* expression. *Plant Physiol* **165**(4): 1671-1683.

**Notes S1** Maps and sequences of plasmids used in this work.

### Index

### pUBQ10GW3HAstrepII7

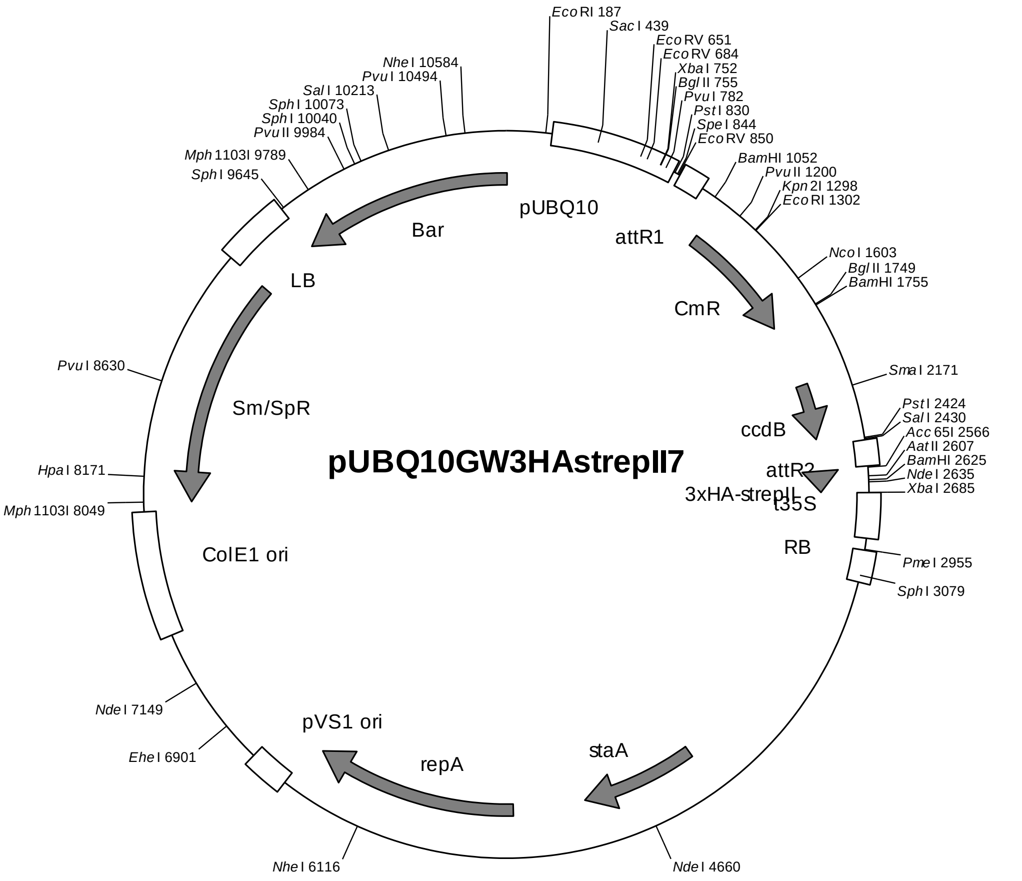

*UBQ10* promoter with 5'-UTR and 1. intron: 218-823
Gateway cassette (attR1, *Cm*R, *ccd*B, attR2): 856-2560
3xHA-strepII-tag: 2568-2684
35S terminator: 2686-2902
Right border: 2959-3112
pVS1 replicon (*sta*A, *rep*A, ori): 4328-6712
ColE1 ori: 8003-7404
Spectinomycin resistance cassette: 9295-8046
Left border: 9301-9633
Basta resistance cassette: 9637-4

sequence:
GATAATTCGAGCTGGCACGACAGGTTTCCCGACTGGAAAGCGGGCAGTGAGCGCAACGCAATTAATGTGAGTTAG
CTCACTCATTAGGCACCCCAGGCTTTACACTTTATGCTTCCGGCTCGTATGTTGTGTGGAATTGTGAGCGGATAA
CAATTTCACACAGGAAACAGCTATGACCATGATTACGAATTCTCATGTTTGACAGCTTATCATCGATGTGGTTGC
AGCCGGCACACACGAGTCGTGTTTATCAACTCAAAGCACAAATACTTTTCCTCAACCTAAAAATAAGGCAATTAG
CCAAAAACAACTTTGCGTGTAAACAACGCTCAATACACGTGTCATTTTATTATTAGCTATTGCTTCACCGCCTTA
GCTTTCTCGTGACCTAGTCGTCCTCGTCTTTTCTTCTTCTTCTTCTATAAAACAATACCCAAAGAGCTCTTCTTC
TTCACAATTCAGATTTCAATTTCTCAAAATCTTAAAAACTTTCTCTCAATTCTCTCTACCGTGATCAAGGTAAAT
TTCTGTGTTCCTTATTCTCTCAAAATCTTCGATTTTGTTTTCGTTCGATCCCAATTTCGTATATGTTCTTTGGTT
TAGATTCTGTTAATCTTAGATCGAAGACGATTTTCTGGGTTTGATCGTTAGATATCATCTTAATTCTCGATTAGG
GTTTCATAGATATCATCCGATTTGTTCAAATAATTTGAGTTTTGTCGAATAATTACTCTTCGATTTGTGATTTCT
ATCTAGATCTGGTGTTAGTTTCTAGTTTGTGCGATCGAATTTGTCGATTAATCTGAGTTTTTCTGATTAACAGCT
CGACCTGCAGGCGGCCGCACTAGTGATATCACAAGTTTGTACAAAAAAGCTGAACGAGAAACGTAAAATGATATA
AATATCAATATATTAAATTAGATTTTGCATAAAAAACAGACTACATAATACTGTAAAACACAACATATCCAGTCA

CTATGGCGGCCGCATTAGGCACCCCAGGCTTTACACTTTATGCTTCCGGCTCGTATAATGTGTGGATTTTGAGTT
AGGATCCGTCGAGATTTTCAGGAGCTAAGGAAGCTAAAATGGAGAAAAAAATCACTGGATATACCACCGTTGATA
TATCCCAATGGCATCGTAAAGAACATTTTGAGGCATTTCAGTCAGTTGCTCAATGTACCTATAACCAGACCGTTC
AGCTGGATATTACGGCCTTTTTAAAGACCGTAAAGAAAAATAAGCACAAGTTTTATCCGGCCTTTATTCACATTC
TTGCCCGCCTGATGAATGCTCATCCGGAATTCCGTATGGCAATGAAAGACGGTGAGCTGGTGATATGGGATAGTG
TTCACCCTTGTTACACCGTTTTCCATGAGCAAACTGAAACGTTTTCATCGCTCTGGAGTGAATACCACGACGATT
TCCGGCAGTTTCTACACATATATTCGCAAGATGTGGCGTGTTACGGTGAAAACCTGGCCTATTTCCCTAAAGGGT
TTATTGAGAATATGTTTTTCGTCTCAGCCAATCCCTGGGTGAGTTTCACCAGTTTTGATTTAAACGTGGCCAATA
TGGACAACTTCTTCGCCCCCGTTTTCACCATGGGCAAATATTATACGCAAGGCGACAAGGTGCTGATGCCGCTGG
CGATTCAGGTTCATCATGCCGTCTGTGATGGCTTCCATGTCGGCAGAATGCTTAATGAATTACAACAGTACTGCG
ATGAGTGGCAGGGCGGGGCGTAAAGATCTGGATCCGGCTTACTAAAAGCCAGATAACAGTATGCGTATTTGCGCG
CTGATTTTTGCGGTATAAGAATATATACTGATATGTATACCCGAAGTATGTCAAAAAGAGGTGTGCTATGAAGCA
GCGTATTACAGTGACAGTTGACAGCGACAGCTATCAGTTGCTCAAGGCATATATGATGTCAATATCTCCGGTCTG
GTAAGCACAACCATGCAGAATGAAGCCCGTCGTCTGCGTGCCGAACGCTGGAAAGCGGAAAATCAGGAAGGGATG
GCTGAGGTCGCCCGGTTTATTGAAATGAACGGCTCTTTTGCTGACGAGAACAGGGACTGGTGAAATGCAGTTTAA
GGTTTACACCTATAAAAGAGAGAGCCGTTATCGTCTGTTTGTGGATGTACAGAGTGATATTATTGACACGCCCGG
GCGACGGATGGTGATCCCCCTGGCCAGTGCACGTCTGCTGTCAGATAAAGTCTCCCGTGAACTTTACCCGGTGGT
GCATATCGGGGATGAAAGCTGGCGCATGATGACCACCGATATGGCCAGTGTGCCGGTCTCCGTTATCGGGGAAGA
AGTGGCTGATCTCAGCCACCGCGAAAATGACATCAAAAACGCCATTAACCTGATGTTCTGGGGAATATAAATGTC
AGGCTCCCTTATACACAGCCAGTCTGCAGGTCGACCATAGTGACTGGATATGTTGTGTTTTACAGTATTATGTAG
TCTGTTTTTTATGCAAAATCTAATTTAATATATTGATATTTATATCATTTTACGTTTCTCGTTCAGCTTTCTTGT
ACAAAGTGGTTGATGGGTACCCATACGATGTTCCTGACTATGCGGGCTATCCCTATGACGTCCCGGACTATGCAG
GATCCTATCCATATGACGTTCCAGATTACGCTTGGTCTCATCCTCAATTTGAAAAATAATCTAGAGTCCGCAAAA
ATCACCAGTCTCTCTCTACAAATCTATCTCTCTCTATTTTTCTCCAGAATAATGTGTGAGTAGTTCCCAGATAAG
GGAATTAGGGTTCTTATAGGGTTTCGCTCATGTGTTGAGCATATAAGAAACCCTTAGTATGTATTTGTATTTGTA
AAATACTTCTATCAATAAAATTTCTAATTCCTAAAACCAAAATCCAGTGACCGGGCGGCCGCCACCGCGGTGGAG
GGGGATCAGATTGTCGTTTCCCGCCTTCAGTTTAAACTATCAGTGTTTGACAGGATATATTGGCGGGTAAACCTA
AGAGAAAAGAGCGTTTATTAGAATAATCGGATATTTAAAAGGGCGTGAAAAGGTTTATCCGTTCGTCCATTTGTA
TGTGCATGCCAACCACAGGGTTCCCCTCGGGATCAAAGTACTTTAAAGTACTTTAAAGTACTTTAAAGTACTTTG
ATCCAACCCCTCCGCTGCTATAGTGCAGTCGGCTTCTGACGTTCAGTGCAGCCGTCTTCTGAAAACGACATGTCG
CACAAGTCCTAAGTTACGCGACAGGCTGCCGCCCTGCCCTTTTCCTGGCGTTTTCTTGTCGCGTGTTTTAGTCGC
ATAAAGTAGAATACTTGCGACTAGAACCGGAGACATTACGCCATGAACAAGAGCGCCGCCGCTGGCCTGCTGGGC
TATGCCCGCGTCAGCACCGACGACCAGGACTTGACCAACCAACGGGCCGAACTGCACGCGGCCGGCTGCACCAAG
CTGTTTTCCGAGAAGATCACCGGCACCAGGCGCGACCGCCCGGAGCTGGCCAGGATGCTTGACCACCTACGCCCT
GGCGACGTTGTGACAGTGACCAGGCTAGACCGCCTGGCCCGCAGCACCCGCGACCTACTGGACATTGCCGAGCGC
ATCCAGGAGGCCGGCGCGGGCCTGCGTAGCCTGGCAGAGCCGTGGGCCGACACCACCACGCCGGCCGGCCGCATG
GTGTTGACCGTGTTCGCCGGCATTGCCGAGTTCGAGCGTTCCCTAATCATCGACCGCACCCGGAGCGGGCGCGAG
GCCGCCAAGGCCCGAGGCGTGAAGTTTGGCCCCCGCCCTACCCTCACCCCGGCACAGATCGCGCACGCCCGCGAG
CTGATCGACCAGGAAGGCCGCACCGTGAAAGAGGCGGCTGCACTGCTTGGCGTGCATCGCTCGACCCTGTACCGC
GCACTTGAGCGCAGCGAGGAAGTGACGCCCACCGAGGCCAGGCGGCGCGGTGCCTTCCGTGAGGACGCATTGACC
GAGGCCGACGCCCTGGCGGCCGCCGAGAATGAACGCCAAGAGGAACAAGCATGAAACCGCACCAGGACGGCCAGG
ACGAACCGTTTTTCATTACCGAAGAGATCGAGGCGGAGATGATCGCGGCCGGGTACGTGTTCGAGCCGCCCGCGC
ACGTCTCAACCGTGCGGCTGCATGAAATCCTGGCCGGTTTGTCTGATGCCAAGCTGGCGGCCTGGCCGGCCAGCT
TGGCCGCTGAAGAAACCGAGCGCCGCCGTCTAAAAAGGTGATGTGTATTTGAGTAAAACAGCTTGCGTCATGCGG
TCGCTGCGTATATGATGCGATGAGTAAATAAACAAATACGCAAGGGGAACGCATGAAGGTTATCGCTGTACTTAA
CCAGAAAGGCGGGTCAGGCAAGACGACCATCGCAACCCATCTAGCCCGCGCCCTGCAACTCGCCGGGGCCGATGT
TCTGTTAGTCGATTCCGATCCCCAGGGCAGTGCCCGCGATTGGGCGGCCGTGCGGGAAGATCAACCGCTAACCGT
TGTCGGCATCGACCGCCCGACGATTGACCGCGACGTGAAGGCCATCGGCCGGCGCGACTTCGTAGTGATCGACGG
AGCGCCCCAGGCGGCGGACTTGGCTGTGTCCGCGATCAAGGCAGCCGACTTCGTGCTGATTCCGGTGCAGCCAAG
CCCTTACGACATATGGGCCACCGCCGACCTGGTGGAGCTGGTTAAGCAGCGCATTGAGGTCACGGATGGAAGGCT
ACAAGCGGCCTTTGTCGTGTCGCGGGCGATCAAAGGCACGCGCATCGGCGGTGAGGTTGCCGAGGCGCTGGCCGG
GTACGAGCTGCCCATTCTTGAGTCCCGTATCACGCAGCGCGTGAGCTACCCAGGCACTGCCGCCGCCGGCACAAC
CGTTCTTGAATCAGAACCCGAGGGCGACGCTGCCCGCGAGGTCCAGGCGCTGGCCGCTGAAATTAAATCAAAACT
CATTTGAGTTAATGAGGTAAAGAGAAAATGAGCAAAAGCACAAACACGCTAAGTGCCGGCCGTCCGAGCGCACGC
AGCAGCAAGGCTGCAACGTTGGCCAGCCTGGCAGACACGCCAGCCATGAAGCGGGTCAACTTTCAGTTGCCGGCG
GAGGATCACACCAAGCTGAAGATGTACGCGGTACGCCAAGGCAAGACCATTACCGAGCTGCTATCTGAATACATC
GCGCAGCTACCAGAGTAAATGAGCAAATGAATAAATGAGTAGATGAATTTTAGCGGCTAAAGGAGGCGGCATGGA
AAATCAAGAACAACCAGGCACCGACGCCGTGGAATGCCCCATGTGTGGAGGAACGGGCGGTTGGCCAGGCGTAAG
CGGCTGGGTTGTCTGCCGGCCCTGCAATGGCACTGGAACCCCCAAGCCCGAGGAATCGGCGTGACGGTCGCAAAC
CATCCGGCCCGGTACAAATCGGCGCGGCGCTGGGTGATGACCTGGTGGAGAAGTTGAAGGCCGCGCAGGCCGCCC
AGCGGCAACGCATCGAGGCAGAAGCACGCCCCGGTGAATCGTGGCAAGCGGCCGCTGATCGAATCCGCAAAGAAT
CCCGGCAACCGCCGGCAGCCGGTGCGCCGTCGATTAGGAAGCCGCCCAAGGGCGACGAGCAACCAGATTTTTTCG
TTCCGATGCTCTATGACGTGGGCACCCGCGATAGTCGCAGCATCATGGACGTGGCCGTTTTCCGTCTGTCGAAGC

GTGACCGACGAGCTGGCGAGGTGATCCGCTACGAGCTTCCAGACGGGCACGTAGAGGTTTCCGCAGGGCCGGCCG
GCATGGCCAGTGTGTGGGATTACGACCTGGTACTGATGGCGGTTTCCCATCTAACCGAATCCATGAACCGATACC
GGGAAGGGAAGGGAGACAAGCCCGGCCGCGTGTTCCGTCCACACGTTGCGGACGTACTCAAGTTCTGCCGGCGAG
CCGATGGCGGAAAGCAGAAAGACGACCTGGTAGAAACCTGCATTCGGTTAAACACCACGCACGTTGCCATGCAGC
GTACGAAGAAGGCCAAGAACGGCCGCCTGGTGACGGTATCCGAGGGTGAAGCCTTGATTAGCCGCTACAAGATCG
TAAAGAGCGAAACCGGGCGGCCGGAGTACATCGAGATCGAGCTAGCTGATTGGATGTACCGCGAGATCACAGAAG
GCAAGAACCCGGACGTGCTGACGGTTCACCCCGATTACTTTTTGATCGATCCCGGCATCGGCCGTTTTCTCTACC
GCCTGGCACGCCGCGCCGCAGGCAAGGCAGAAGCCAGATGGTTGTTCAAGACGATCTACGAACGCAGTGGCAGCG
CCGGAGAGTTCAAGAAGTTCTGTTTCACCGTGCGCAAGCTGATCGGGTCAAATGACCTGCCGGAGTACGATTTGA
AGGAGGAGGCGGGGCAGGCTGGCCCGATCCTAGTCATGCGCTACCGCAACCTGATCGAGGGCGAAGCATCCGCCG
GTTCCTAATGTACGGAGCAGATGCTAGGGCAAATTGCCCTAGCAGGGGAAAAAGGTCGAAAAGGTCTCTTTCCTG
TGGATAGCACGTACATTGGGAACCCAAAGCCGTACATTGGGAACCGGAACCCGTACATTGGGAACCCAAAGCCGT
ACATTGGGAACCGGTCACACATGTAAGTGACTGATATAAAAGAGAAAAAAGGCGATTTTTCCGCCTAAAACTCTT
TAAAACTTATTAAAACTCTTAAAACCCGCCTGGCCTGTGCATAACTGTCTGGCCAGCGCACAGCCGAAGAGCTGC
AAAAAGCGCCTACCCTTCGGTCGCTGCGCTCCCTACGCCCCGCCGCTTCGCGTCGGCCTATCGCGGCCGCTGGCC
GCTCAAAAATGGCTGGCCTACGGCCAGGCAATCTACCAGGGCGCGGACAAGCCGCGCCGTCGCCACTCGACCGCC
GGCGCCCACATCAAGGCACCCTGCCTCGCGCGTTTCGGTGATGACGGTGAAAACCTCTGACACATGCAGCTCCCG
GAGACGGTCACAGCTTGTCTGTAAGCGGATGCCGGGAGCAGACAAGCCCGTCAGGGCGCGTCAGCGGGTGTTGGC
GGGTGTCGGGGCGCAGCCATGACCCAGTCACGTAGCGATAGCGGAGTGTATACTGGCTTAACTATGCGGCATCAG
AGCAGATTGTACTGAGAGTGCACCATATGCGGTGTGAAATACCGCACAGATGCGTAAGGAGAAAATACCGCATCA
GGCGCTCTTCCGCTTCCTCGCTCACTGACTCGCTGCGCTCGGTCGTTCGGCTGCGGCGAGCGGTATCAGCTCACT
CAAAGGCGGTAATACGGTTATCCACAGAATCAGGGGATAACGCAGGAAAGAACATGTGAGCAAAAGGCCAGCAAA
AGGCCAGGAACCGTAAAAAGGCCGCGTTGCTGGCGTTTTTCCATAGGCTCCGCCCCCCTGACGAGCATCACAAAA
ATCGACGCTCAAGTCAGAGGTGGCGAAACCCGACAGGACTATAAAGATACCAGGCGTTTCCCCCTGGAAGCTCCC
TCGTGCGCTCTCCTGTTCCGACCCTGCCGCTTACCGGATACCTGTCCGCCTTTCTCCCTTCGGGAAGCGTGGCGC
TTTCTCATAGCTCACGCTGTAGGTATCTCAGTTCGGTGTAGGTCGTTCGCTCCAAGCTGGGCTGTGTGCACGAAC
CCCCCGTTCAGCCCGACCGCTGCGCCTTATCCGGTAACTATCGTCTTGAGTCCAACCCGGTAAGACACGACTTAT
CGCCACTGGCAGCAGCCACTGGTAACAGGATTAGCAGAGCGAGGTATGTAGGCGGTGCTACAGAGTTCTTGAAGT
GGTGGCCTAACTACGGCTACACTAGAAGGACAGTATTTGGTATCTGCGCTCTGCTGAAGCCAGTTACCTTCGGAA
AAAGAGTTGGTAGCTCTTGATCCGGCAAACAAACCACCGCTGGTAGCGGTGGTTTTTTTGTTTGCAAGCAGCAGA
TTACGCGCAGAAAAAAAGGATCTCAAGAAGATCCTTTGATCTTTTCTACGGGGTCTGACGCTCAGTGGAACGAAA
ACTCACGTTAAGGGATTTTGGTCATGCATGATATATCTCCCAATTTGTGTAGGGCTTATTATGCACGCTTAAAAA
TAATAAAAGCAGACTTGACCTGATAGTTTGGCTGTGAGCAATTATGTGCTTAGTGCATCTAATCGCTTGAGTTAA
CGCCGGCGAAGCGGCGTCGGCTTGAACGAATTTCTAGCTAGACATTATTTGCCGACTACCTTGGTGATCTCGCCT
TTCACGTAGTGGACAAATTCTTCCAACTGATCTGCGCGCGAGGCCAAGCGATCTTCTTCTTGTCCAAGATAAGCC
TGTCTAGCTTCAAGTATGACGGGCTGATACTGGGCCGGCAGGCGCTCCATTGCCCAGTCGGCAGCGACATCCTTC
GGCGCGATTTTGCCGGTTACTGCGCTGTACCAAATGCGGGACAACGTAAGCACTACATTTCGCTCATCGCCAGCC
CAGTCGGGCGGCGAGTTCCATAGCGTTAAGGTTTCATTTAGCGCCTCAAATAGATCCTGTTCAGGAACCGGATCA
AAGAGTTCCTCCGCCGCTGGACCTACCAAGGCAACGCTATGTTCTCTTGCTTTTGTCAGCAAGATAGCCAGATCA
ATGTCGATCGTGGCTGGCTCGAAGATACCTGCAAGAATGTCATTGCGCTGCCATTCTCCAAATTGCAGTTCGCGC
TTAGCTGGATAACGCCACGGAATGATGTCGTCGTGCACAACAATGGTGACTTCTACAGCGCGGAGAATCTCGCTC
TCTCCAGGGGAAGCCGAAGTTTCCAAAAGGTCGTTGATCAAAGCTCGCCGCGTTGTTTCATCAAGCCTTACGGTC
ACCGTAACCAGCAAATCAATATCACTGTGTGGCTTCAGGCCGCCATCCACTGCGGAGCCGTACAAATGTACGGCC
AGCAACGTCGGTTCGAGATGGCGCTCGATGACGCCAACTACCTCTGATAGTTGAGTCGATACTTCGGCGATCACC
GCTTCCCCCATGATGTTTAACTTTGTTTTAGGGCGACTGCCCTGCTGCGTAACATCGTTGCTGCTCCATAACATC
AAACATCGACCCACGGCGTAACGCGCTTGCTGCTTGGATGCCCGAGGCATAGACTGTACCCCAAAAAAACATGTC
ATAACAAGAAGCCATGAAAACCGCCACTGCGCCGTTACCACCGCTGCGTTCGGTCAAGGTTCTGGACCAGTTGCG
TGACGGCAGTTACGCTACTTGCATTACAGCTTACGAACCGAACGAGGCTTATGTCCACTGGGTTCGTGCCCGAAT
TGATCACAGGCAGCAACGCTCTGTCATCGTTACAATCAACATGCTACCCTCCGCGAGATCATCCGTGTTTCAAAC
CCGGCAGCTTAGTTGCCGTTCTTCCGAATAGCATCGGTAACATGAGCAAAGTCTGCCGCCTTACAACGGCTCTCC
CGCTGACGCCGTCCCGGACTGATGGGCTGCCTGTATCGAGTGGTGATTTTGTGCCGAGCTGCCGGTCGGGGAGCT
GTTGGCTGGCTGGTGGCAGGATATATTGTGGTGTAAACAAATTGACGCTTAGACAACTTAATAACACATTGCGGA
CGTTTTTAATGTACTGAATTAACGCCGAATTGAATTATCAGCTTGCATGCCGGTCGATCTAGTAACATATAGATG
ACACCGCGCGCGATAATTTATCCTAGTTTGCGCGCTATATTTTGTTTTCTATCGCGTATTAAATGTATAATTGCG
GGACTCTAATCATAAAAACCCATCTCATAAATAACGTCATGCATTACATGTTAATTATTACATGCTTAACGTAAT
TCAACAGAAATTATATGATAATCATCGCAAGACCGGCAACAGGATTCAATCTTAAGAAACTTTATTGCCAAATGT
TTGAACGATCTGCTTGACTCTAGGGGTCATCAGATTTCGGTGACGGGCAGGACCGGACGGGGCGGCACCGGCAGG
CTGAAGTCCAGCTGCCAGAAACCCACGTCATGCCAGTTCCCGTGCTTGAAGCCGGCCGCCCGCAGCATGCCGCGG
GGGGCATATCCGAGCGCCTCGTGCATGCGCACGCTCGGGTCGTTGGGCAGCCCGATGACAGCGACCACGCTCTTG
AAGCCCTGTGCCTCCAGGGACTTCAGCAGGTGGGTGTAGAGCGTGGAGCCCAGTCCCGTCCGCTGGTGGCGGGGG
GAGACGTACACGGTCGACTCGGCCGTCCAGTCGTAGGCGTTGCGTGCCTTCCAGGGACCCGCGTAGGCGATGCCG
GCGACCTCGCCGTCCACCTCGGCGACGAGCCAGGGATAGCGCTCCCGCAGACGGACGAGGTCGTCCGTCCACTCC
TGCGGTTCCTGCGGCTCGGTACGGAAGTTGACCGTGCTTGTCTCGATGTAGTGGTTGACGATGGTGCAGACCGCC

GGCATGTCCGCCTCGGTGGCACGGCGGATGTCGGCCGGGCGTCGTTCTGGGCTCATGGTAGATCCCCTCGATCGA
GTTGAGAGTGAATATGAGACTCTAATTGGATACCGAGGGGAATTTATGGAACGTCAGTGGAGCATTTTTGACAAG
AAATATTTGCTAGCTGATAGTGACCTTAGGCGACTTTTGAACGCGCAATAATGGTTTCTGACGTATGTGCTTAGC
TCATTAAACTCCAGAAACCCGCGGCTCAGTGGCTCCTTCAACGTTGCGGTTCTGTCAGTTCCAAACGTAAAACGG
CTTGTCCCGCGTCATCGGCGGGGGTCATAACGTGACTCCCTTAATTCTCATGTAT

### pUBQ10GWVP7

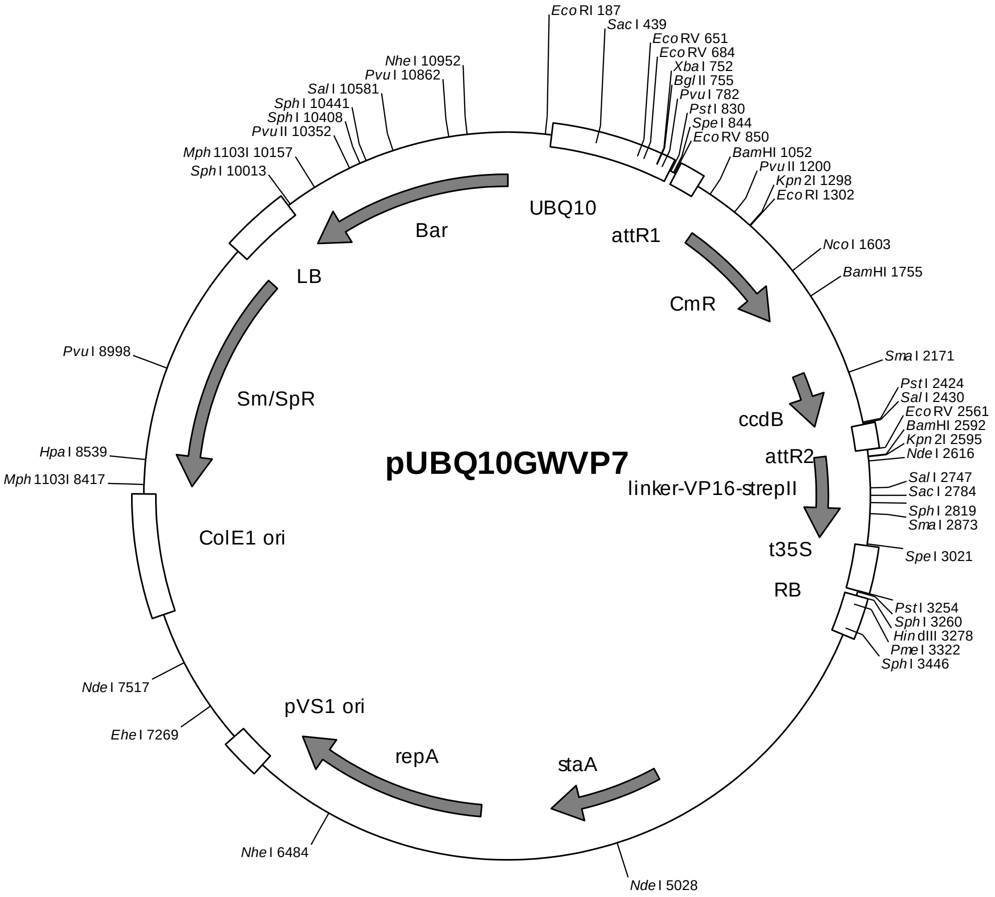

*UBQ10* promoter with 5'-UTR and 1. intron: 218-823
Gateway cassette (attR1, *Cm*R, *ccd*B, attR2): 856-2560
linker/VP16 activation domain coding region/strepII-tag: 2568-3020
35S terminator: 3030-3255
Right border: 3279-3480
pVS1 replicon (*sta*A, *rep*A, ori): 4696-7080
ColE1 ori: 8371-7772
Spectinomycin resistance cassette: 9663-8414
Left border: 9669-10001
Basta resistance cassette: 10005-4

sequence:

GATAATTCGAGCTGGCACGACAGGTTTCCCGACTGGAAAGCGGGCAGTGAGCGCAACGCAATTAATGTGAGTTAG
CTCACTCATTAGGCACCCCAGGCTTTACACTTTATGCTTCCGGCTCGTATGTTGTGTGGAATTGTGAGCGGATAA
CAATTTCACACAGGAAACAGCTATGACCATGATTACGAATTCTCATGTTTGACAGCTTATCATCGATGTGGTTGC
AGCCGGCACACACGAGTCGTGTTTATCAACTCAAAGCACAAATACTTTTCCTCAACCTAAAAATAAGGCAATTAG
CCAAAAACAACTTTGCGTGTAAACAACGCTCAATACACGTGTCATTTTATTATTAGCTATTGCTTCACCGCCTTA
GCTTTCTCGTGACCTAGTCGTCCTCGTCTTTTCTTCTTCTTCTTCTATAAAACAATACCCAAAGAGCTCTTCTTC
TTCACAATTCAGATTTCAATTTCTCAAAATCTTAAAAACTTTCTCTCAATTCTCTCTACCGTGATCAAGGTAAAT
TTCTGTGTTCCTTATTCTCTCAAAATCTTCGATTTTGTTTTCGTTCGATCCCAATTTCGTATATGTTCTTTGGTT
TAGATTCTGTTAATCTTAGATCGAAGACGATTTTCTGGGTTTGATCGTTAGATATCATCTTAATTCTCGATTAGG
GTTTCATAGATATCATCCGATTTGTTCAAATAATTTGAGTTTTGTCGAATAATTACTCTTCGATTTGTGATTTCT
ATCTAGATCTGGTGTTAGTTTCTAGTTTGTGCGATCGAATTTGTCGATTAATCTGAGTTTTTCTGATTAACAGCT

CGACCTGCAGGCGGCCGCACTAGTGATATCACAAGTTTGTACAAAAAAGCTGAACGAGAAACGTAAAATGATATA
AATATCAATATATTAAATTAGATTTTGCATAAAAAACAGACTACATAATACTGTAAAACACAACATATCCAGTCA
CTATGGCGGCCGCATTAGGCACCCCAGGCTTTACACTTTATGCTTCCGGCTCGTATAATGTGTGGATTTTGAGTT
AGGATCCGTCGAGATTTTCAGGAGCTAAGGAAGCTAAAATGGAGAAAAAAATCACTGGATATACCACCGTTGATA
TATCCCAATGGCATCGTAAAGAACATTTTGAGGCATTTCAGTCAGTTGCTCAATGTACCTATAACCAGACCGTTC
AGCTGGATATTACGGCCTTTTTAAAGACCGTAAAGAAAAATAAGCACAAGTTTTATCCGGCCTTTATTCACATTC
TTGCCCGCCTGATGAATGCTCATCCGGAATTCCGTATGGCAATGAAAGACGGTGAGCTGGTGATATGGGATAGTG
TTCACCCTTGTTACACCGTTTTCCATGAGCAAACTGAAACGTTTTCATCGCTCTGGAGTGAATACCACGACGATT
TCCGGCAGTTTCTACACATATATTCGCAAGATGTGGCGTGTTACGGTGAAAACCTGGCCTATTTCCCTAAAGGGT
TTATTGAGAATATGTTTTTCGTCTCAGCCAATCCCTGGGTGAGTTTCACCAGTTTTGATTTAAACGTGGCCAATA
TGGACAACTTCTTCGCCCCCGTTTTCACCATGGGCAAATATTATACGCAAGGCGACAAGGTGCTGATGCCGCTGG
CGATTCAGGTTCATCATGCCGTTTGTGATGGCTTCCATGTCGGCAGAATGCTTAATGAATTACAACAGTACTGCG
ATGAGTGGCAGGGCGGGGCGTAAACGCGTGGATCCGGCTTACTAAAAGCCAGATAACAGTATGCGTATTTGCGCG
CTGATTTTTGCGGTATAAGAATATATACTGATATGTATACCCGAAGTATGTCAAAAAGAGGTATGCTATGAAGCA
GCGTATTACAGTGACAGTTGACAGCGACAGCTATCAGTTGCTCAAGGCATATATGATGTCAATATCTCCGGTCTG
GTAAGCACAACCATGCAGAATGAAGCCCGTCGTCTGCGTGCCGAACGCTGGAAAGCGGAAAATCAGGAAGGGATG
GCTGAGGTCGCCCGGTTTATTGAAATGAACGGCTCTTTTGCTGACGAGAACAGGGGCTGGTGAAATGCAGTTTAA
GGTTTACACCTATAAAAGAGAGAGCCGTTATCGTCTGTTTGTGGATGTACAGAGTGATATTATTGACACGCCCGG
GCGACGGATGGTGATCCCCCTGGCCAGTGCACGTCTGCTGTCAGATAAAGTCTCCCGTGAACTTTACCCGGTGGT
GCATATCGGGGATGAAAGCTGGCGCATGATGACCACCGATATGGCCAGTGTGCCGGTCTCCGTTATCGGGGAAGA
AGTGGCTGATCTCAGCCACCGCGAAAATGACATCAAAAACGCCATTAACCTGATGTTCTGGGGAATATAAATGTC
AGGCTCCCTTATACACAGCCAGTCTGCAGGTCGACCATAGTGACTGGATATGTTGTGTTTTACAGTATTATGTAG
TCTGTTTTTTATGCAAAATCTAATTTAATATATTGATATTTATATCATTTTACGTTTCTCGTTCAGCTTTCTTGT
ACAAAGTGGTGATATCAGGAGGAGGAGGTTCAGGTGGTGGTGGATCCGGAGGAGGTGGTTCAATTCATATGACGA
AAAACAATTACGGGTCTACCATCGAGGGCCTGCTCGATCTCCCGGACGACGACGCCCCCGAAGAGGCGGGGCTGG
CGGCTCCGCGCCTGTCCTTTCTCCCCGCGGACACACGCGCAGACTGTCGACGGCCCCCCCGACCGATGTCAGCC
TGGGGGACGAGCTCCACTTAGACGGCGAGGACGTGGCGATGGCGCATGCCGACGCGCTAGACGATTTCGATCTGG
ACATGTTGGGGGACGGGGATTCCCCGGGTCCGGGATTTACCCCCCACGACTCCGCCCCCTACGGCGCTCTGGATA
TGGCCGACTTCGAGTTTGAGCAGATGTTTACCGATGCCCTTGGAATTGACGAGTACGGTGGGCGTACGTGGAGTC
ATCCGCAGTTCGAAAAGTAGACTAGTCCGCGGCCATGCTAGAGTCCGCAAAAATCACCAGTCTCTCTCTACAAAT
CTATCTCTCTCTATTTTTCTCCAGAATAATGTGTGAGTAGTTCCCAGATAAGGGAATTAGGGTTCTTATAGGGTT
TCGCTCATGTGTTGAGCATATAAGAAACCCTTAGTATGTATTTGTATTTGTAAAATACTTCTATCAATAAAATTT
CTAATTCCTAAAACCAAAATCCAGTGACCTGCAGGCATGCGACGTCGGGCCCAAGCTTAGCTTGAGCTTGGATCA
GATTGTCGTTTCCCGCCTTCAGTTTAAACTATCAGTGTTTGACAGGATATATTGGCGGGTAAACCTAAGAGAAAA
GAGCGTTTATTAGAATAATCGGATATTTAAAAGGGCGTGAAAAGGTTTATCCGTTCGTCCATTTGTATGTGCATG
CCAACCACAGGGTTCCCCTCGGGGATCAAAGTACTTTAAAGTACTTTAAAGTACTTTAAAGTACTTTGATCCAAC
CCCTCCGCTGCTATAGTGCAGTCGGCTTCTGACGTTCAGTGCAGCCGTCTTCTGAAAACGACATGTCGCACAAGT
CCTAAGTTACGCGACAGGCTGCCGCCCTGCCCTTTTCCTGGCGTTTTCTTGTCGCGTGTTTTAGTCGCATAAAGT
AGAATACTTGCGACTAGAACCGGAGACATTACGCCATGAACAAGAGCGCCGCCGCTGGCCTGCTGGGCTATGCCC
GCGTCAGCACCGACGACCAGGACTTGACCAACCAACGGGCCGAACTGCACGCGGCCGGCTGCACCAAGCTGTTTT
CCGAGAAGATCACCGGCACCAGGCGCGACCGCCCGGAGCTGGCCAGGATGCTTGACCACCTACGCCCTGGCGACG
TTGTGACAGTGACCAGGCTAGACCGCCTGGCCCGCAGCACCCGCGACCTACTGGACATTGCCGAGCGCATCCAGG
AGGCCGGCGCGGGCCTGCGTAGCCTGGCAGAGCCGTGGGCCGACACCACCACGCCGGCCGGCCGCATGGTGTTGA
CCGTGTTCGCCGGCATTGCCGAGTTCGAGCGTTCCCTAATCATCGACCGCACCCGGAGCGGGCGCGAGGCCGCCA
AGGCCCGAGGCGTGAAGTTTGGCCCCCGCCCTACCCTCACCCCGGCACAGATCGCGCACGCCCGCGAGCTGATCG
ACCAGGAAGGCCGCACCGTGAAAGAGGCGGCTGCACTGCTTGGCGTGCATCGCTCGACCCTGTACCGCGCACTTG
AGCGCAGCGAGGAAGTGACGCCCACCGAGGCCAGGCGGCGCGGTGCCTTCCGTGAGGACGCATTGACCGAGGCCG
ACGCCCTGGCGGCCGCCGAGAATGAACGCCAAGAGGAACAAGCATGAAACCGCACCAGGACGGCCAGGACGAACC
GTTTTTCATTACCGAAGAGATCGAGGCGGAGATGATCGCGGCCGGGTACGTGTTCGAGCCGCCCGCGCACGTCTC
AACCGTGCGGCTGCATGAAATCCTGGCCGGTTTGTCTGATGCCAAGCTGGCGGCCTGGCCGGCCAGCTTGGCCGC
TGAAGAAACCGAGCGCCGCCGTCTAAAAAGGTGATGTGTATTTGAGTAAAACAGCTTGCGTCATGCGGTCGCTGC
GTATATGATGCGATGAGTAAATAAACAAATACGCAAGGGGAACGCATGAAGGTTATCGCTGTACTTAACCAGAAA
GGCGGGTCAGGCAAGACGACCATCGCAACCCATCTAGCCCGCGCCCTGCAACTCGCCGGGGCCGATGTTCTGTTA
GTCGATTCCGATCCCCAGGGCAGTGCCCGCGATTGGGCGGCCGTGCGGGAAGATCAACCGCTAACCGTTGTCGGC
ATCGACCGCCCGACGATTGACCGCGACGTGAAGGCCATCGGCCGGCGCGACTTCGTAGTGATCGACGGAGCGCCC
CAGGCGGCGGACTTGGCTGTGTCCGCGATCAAGGCAGCCGACTTCGTGCTGATTCCGGTGCAGCCAAGCCCTTAC
GACATATGGGCCACCGCCGACCTGGTGGAGCTGGTTAAGCAGCGCATTGAGGTCACGGATGGAAGGCTACAAGCG
GCCTTTGTCGTGTCGCGGGCGATCAAAGGCACGCGCATCGGCGGTGAGGTTGCCGAGGCGCTGGCCGGGTACGAG
CTGCCCATTCTTGAGTCCCGTATCACGCAGCGCGTGAGCTACCCAGGCACTGCCGCCGCCGGCACAACCGTTCTT
GAATCAGAACCCGAGGGCGACGCTGCCCGCGAGGTCCAGGCGCTGGCCGCTGAAATTAAATCAAAACTCATTTGA
GTTAATGAGGTAAAGAGAAAATGAGCAAAAGCACAAACACGCTAAGTGCCGGCCGTCCGAGCGCACGCAGCAGCA
AGGCTGCAACGTTGGCCAGCCTGGCAGACACGCCAGCCATGAAGCGGGTCAACTTTCAGTTGCCGGCGGAGGATC
ACACCAAGCTGAAGATGTACGCGGTACGCCAAGGCAAGACCATTACCGAGCTGCTATCTGAATACATCGCGCAGC

TACCAGAGTAAATGAGCAAATGAATAAATGAGTAGATGAATTTTAGCGGCTAAAGGAGGCGGCATGGAAAATCAA
GAACAACCAGGCACCGACGCCGTGGAATGCCCCATGTGTGGAGGAACGGGCGGTTGGCCAGGCGTAAGCGGCTGG
GTTGTCTGCCGGCCCTGCAATGGCACTGGAACCCCCAAGCCCGAGGAATCGGCGTGACGGTCGCAAACCATCCGG
CCCGGTACAAATCGGCGCGGCGCTGGGTGATGACCTGGTGGAGAAGTTGAAGGCCGCGCAGGCCGCCCAGCGGCA
ACGCATCGAGGCAGAAGCACGCCCCGGTGAATCGTGGCAAGCGGCCGCTGATCGAATCCGCAAAGAATCCCGGCA
ACCGCCGGCAGCCGGTGCGCCGTCGATTAGGAAGCCGCCCAAGGGCGACGAGCAACCAGATTTTTTCGTTCCGAT
GCTCTATGACGTGGGCACCCGCGATAGTCGCAGCATCATGGACGTGGCCGTTTTCCGTCTGTCGAAGCGTGACCG
ACGAGCTGGCGAGGTGATCCGCTACGAGCTTCCAGACGGGCACGTAGAGGTTTCCGCAGGGCCGGCCGGCATGGC
CAGTGTGTGGGATTACGACCTGGTACTGATGGCGGTTTCCCATCTAACCGAATCCATGAACCGATACCGGGAAGG
GAAGGGAGACAAGCCCGGCCGCGTGTTCCGTCCACACGTTGCGGACGTACTCAAGTTCTGCCGGCGAGCCGATGG
CGGAAAGCAGAAAGACGACCTGGTAGAAACCTGCATTCGGTTAAACACCACGCACGTTGCCATGCAGCGTACGAA
GAAGGCCAAGAACGGCCGCCTGGTGACGGTATCCGAGGGTGAAGCCTTGATTAGCCGCTACAAGATCGTAAAGAG
CGAAACCGGGCGGCCGGAGTACATCGAGATCGAGCTAGCTGATTGGATGTACCGCGAGATCACAGAAGGCAAGAA
CCCGGACGTGCTGACGGTTCACCCCGATTACTTTTTGATCGATCCCGGCATCGGCCGTTTTCTCTACCGCCTGGC
ACGCCGCGCCGCAGGCAAGGCAGAAGCCAGATGGTTGTTCAAGACGATCTACGAACGCAGTGGCAGCGCCGGAGA
GTTCAAGAAGTTCTGTTTCACCGTGCGCAAGCTGATCGGGTCAAATGACCTGCCGGAGTACGATTTGAAGGAGGA
GGCGGGGCAGGCTGGCCCGATCCTAGTCATGCGCTACCGCAACCTGATCGAGGGCGAAGCATCCGCCGGTTCCTA
ATGTACGGAGCAGATGCTAGGGCAAATTGCCCTAGCAGGGGAAAAAGGTCGAAAAGGTCTCTTTCCTGTGGATAG
CACGTACATTGGGAACCCAAAGCCGTACATTGGGAACCGGAACCCGTACATTGGGAACCCAAAGCCGTACATTGG
GAACCGGTCACACATGTAAGTGACTGATATAAAAGAGAAAAAAGGCGATTTTTCCGCCTAAAACTCTTTAAAACT
TATTAAAACTCTTAAAACCCGCCTGGCCTGTGCATAACTGTCTGGCCAGCGCACAGCCGAAGAGCTGCAAAAAGC
GCCTACCCTTCGGTCGCTGCGCTCCCTACGCCCCGCCGCTTCGCGTCGGCCTATCGCGGCCGCTGGCCGCTCAAA
AATGGCTGGCCTACGGCCAGGCAATCTACCAGGGCGCGGACAAGCCGCGCCGTCGCCACTCGACCGCCGGCGCCC
ACATCAAGGCACCCTGCCTCGCGCGTTTCGGTGATGACGGTGAAAACCTCTGACACATGCAGCTCCCGGAGACGG
TCACAGCTTGTCTGTAAGCGGATGCCGGGAGCAGACAAGCCCGTCAGGGCGCGTCAGCGGGTGTTGGCGGGTGTC
GGGGCGCAGCCATGACCCAGTCACGTAGCGATAGCGGAGTGTATACTGGCTTAACTATGCGGCATCAGAGCAGAT
TGTACTGAGAGTGCACCATATGCGGTGTGAAATACCGCACAGATGCGTAAGGAGAAAATACCGCATCAGGCGCTC
TTCCGCTTCCTCGCTCACTGACTCGCTGCGCTCGGTCGTTCGGCTGCGGCGAGCGGTATCAGCTCACTCAAAGGC
GGTAATACGGTTATCCACAGAATCAGGGGATAACGCAGGAAAGAACATGTGAGCAAAAGGCCAGCAAAAGGCCAG
GAACCGTAAAAAGGCCGCGTTGCTGGCGTTTTTCCATAGGCTCCGCCCCCCTGACGAGCATCACAAAAATCGACG
CTCAAGTCAGAGGTGGCGAAACCCGACAGGACTATAAAGATACCAGGCGTTTCCCCCTGGAAGCTCCCTCGTGCG
CTCTCCTGTTCCGACCCTGCCGCTTACCGGATACCTGTCCGCCTTTCTCCCTTCGGGAAGCGTGGCGCTTTCTCA
TAGCTCACGCTGTAGGTATCTCAGTTCGGTGTAGGTCGTTCGCTCCAAGCTGGGCTGTGTGCACGAACCCCCCGT
TCAGCCCGACCGCTGCGCCTTATCCGGTAACTATCGTCTTGAGTCCAACCCGGTAAGACACGACTTATCGCCACT
GGCAGCAGCCACTGGTAACAGGATTAGCAGAGCGAGGTATGTAGGCGGTGCTACAGAGTTCTTGAAGTGGTGGCC
TAACTACGGCTACACTAGAAGGACAGTATTTGGTATCTGCGCTCTGCTGAAGCCAGTTACCTTCGGAAAAAGAGT
TGGTAGCTCTTGATCCGGCAAACAAACCACCGCTGGTAGCGGTGGTTTTTTTGTTTGCAAGCAGCAGATTACGCG
CAGAAAAAAAGGATCTCAAGAAGATCCTTTGATCTTTTCTACGGGGTCTGACGCTCAGTGGAACGAAAACTCACG
TTAAGGGATTTTGGTCATGCATGATATATCTCCCAATTTGTGTAGGGCTTATTATGCACGCTTAAAAATAATAAA
AGCAGACTTGACCTGATAGTTTGGCTGTGAGCAATTATGTGCTTAGTGCATCTAATCGCTTGAGTTAACGCCGGC
GAAGCGGCGTCGGCTTGAACGAATTTCTAGCTAGACATTATTTGCCGACTACCTTGGTGATCTCGCCTTTCACGT
AGTGGACAAATTCTTCCAACTGATCTGCGCGCGAGGCCAAGCGATCTTCTTCTTGTCCAAGATAAGCCTGTCTAG
CTTCAAGTATGACGGGCTGATACTGGGCCGGCAGGCGCTCCATTGCCCAGTCGGCAGCGACATCCTTCGGCGCGA
TTTTGCCGGTTACTGCGCTGTACCAAATGCGGGACAACGTAAGCACTACATTTCGCTCATCGCCAGCCCAGTCGG
GCGGCGAGTTCCATAGCGTTAAGGTTTCATTTAGCGCCTCAAATAGATCCTGTTCAGGAACCGGATCAAAGAGTT
CCTCCGCCGCTGGACCTACCAAGGCAACGCTATGTTCTCTTGCTTTTGTCAGCAAGATAGCCAGATCAATGTCGA
TCGTGGCTGGCTCGAAGATACCTGCAAGAATGTCATTGCGCTGCCATTCTCCAAATTGCAGTTCGCGCTTAGCTG
GATAACGCCACGGAATGATGTCGTCGTGCACAACAATGGTGACTTCTACAGCGCGGAGAATCTCGCTCTCTCCAG
GGGAAGCCGAAGTTTCCAAAAGGTCGTTGATCAAAGCTCGCCGCGTTGTTTCATCAAGCCTTACGGTCACCGTAA
CCAGCAAATCAATATCACTGTGTGGCTTCAGGCCGCCATCCACTGCGGAGCCGTACAAATGTACGGCCAGCAACG
TCGGTTCGAGATGGCGCTCGATGACGCCAACTACCTCTGATAGTTGAGTCGATACTTCGGCGATCACCGCTTCCC
CCATGATGTTTAACTTTGTTTTAGGGCGACTGCCCTGCTGCGTAACATCGTTGCTGCTCCATAACATCAAACATC
GACCCACGGCGTAACGCGCTTGCTGCTTGGATGCCCGAGGCATAGACTGTACCCCAAAAAAACATGTCATAACAA
GAAGCCATGAAAACCGCCACTGCGCCGTTACCACCGCTGCGTTCGGTCAAGGTTCTGGACCAGTTGCGTGACGGC
AGTTACGCTACTTGCATTACAGCTTACGAACCGAACGAGGCTTATGTCCACTGGGTTCGTGCCCGAATTGATCAC
AGGCAGCAACGCTCTGTCATCGTTACAATCAACATGCTACCCTCCGCGAGATCATCCGTGTTTCAAACCCGGCAG
CTTAGTTGCCGTTCTTCCGAATAGCATCGGTAACATGAGCAAAGTCTGCCGCCTTACAACGGCTCTCCCGCTGAC
GCCGTCCCGGACTGATGGGCTGCCTGTATCGAGTGGTGATTTTGTGCCGAGCTGCCGGTCGGGGAGCTGTTGGCT
GGCTGGTGGCAGGATATATTGTGGTGTAAACAAATTGACGCTTAGACAACTTAATAACACATTGCGGACGTTTTT
AATGTACTGAATTAACGCCGAATTGAATTATCAGCTTGCATGCCGGTCGATCTAGTAACATATAGATGACACCGC
GCGCGATAATTTATCCTAGTTTGCGCGCTATATTTTGTTTTCTATCGCGTATTAAATGTATAATTGCGGGACTCT
AATCATAAAAACCCATCTCATAAATAACGTCATGCATTACATGTTAATTATTACATGCTTAACGTAATTCAACAG
AAATTATATGATAATCATCGCAAGACCGGCAACAGGATTCAATCTTAAGAAACTTTATTGCCAAATGTTTGAACG

ATCTGCTTGACTCTAGGGGTCATCAGATTTCGGTGACGGGCAGGACCGGACGGGGCGGCACCGGCAGGCTGAAGT
CCAGCTGCCAGAAACCCACGTCATGCCAGTTCCCGTGCTTGAAGCCGGCCGCCCGCAGCATGCCGCGGGGGGCAT
ATCCGAGCGCCTCGTGCATGCGCACGCTCGGGTCGTTGGGCAGCCCGATGACAGCGACCACGCTCTTGAAGCCCT
GTGCCTCCAGGGACTTCAGCAGGTGGGTGTAGAGCGTGGAGCCCAGTCCCGTCCGCTGGTGGCGGGGGGAGACGT
ACACGGTCGACTCGGCCGTCCAGTCGTAGGCGTTGCGTGCCTTCCAGGGACCCGCGTAGGCGATGCCGGCGACCT
CGCCGTCCACCTCGGCGACGAGCCAGGGATAGCGCTCCCGCAGACGGACGAGGTCGTCCGTCCACTCCTGCGGTT
CCTGCGGCTCGGTACGGAAGTTGACCGTGCTTGTCTCGATGTAGTGGTTGACGATGGTGCAGACCGCCGGCATGT
CCGCCTCGGTGGCACGGCGGATGTCGGCCGGGCGTCGTTCTGGGCTCATGGTAGATCCCCTCGATCGAGTTGAGA
GTGAATATGAGACTCTAATTGGATACCGAGGGGAATTTATGGAACGTCAGTGGAGCATTTTTGACAAGAAATATT
TGCTAGCTGATAGTGACCTTAGGCGACTTTTGAACGCGCAATAATGGTTTCTGACGTATGTGCTTAGCTCATTAA
ACTCCAGAAACCCGCGGCTCAGTGGCTCCTTCAACGTTGCGGTTCTGTCAGTTCCAAACGTAAAACGGCTTGTCC
CGCGTCATCGGCGGGGGTCATAACGTGACTCCCTTAATTCTCATGTAT

#### pB-DLO1promLUC

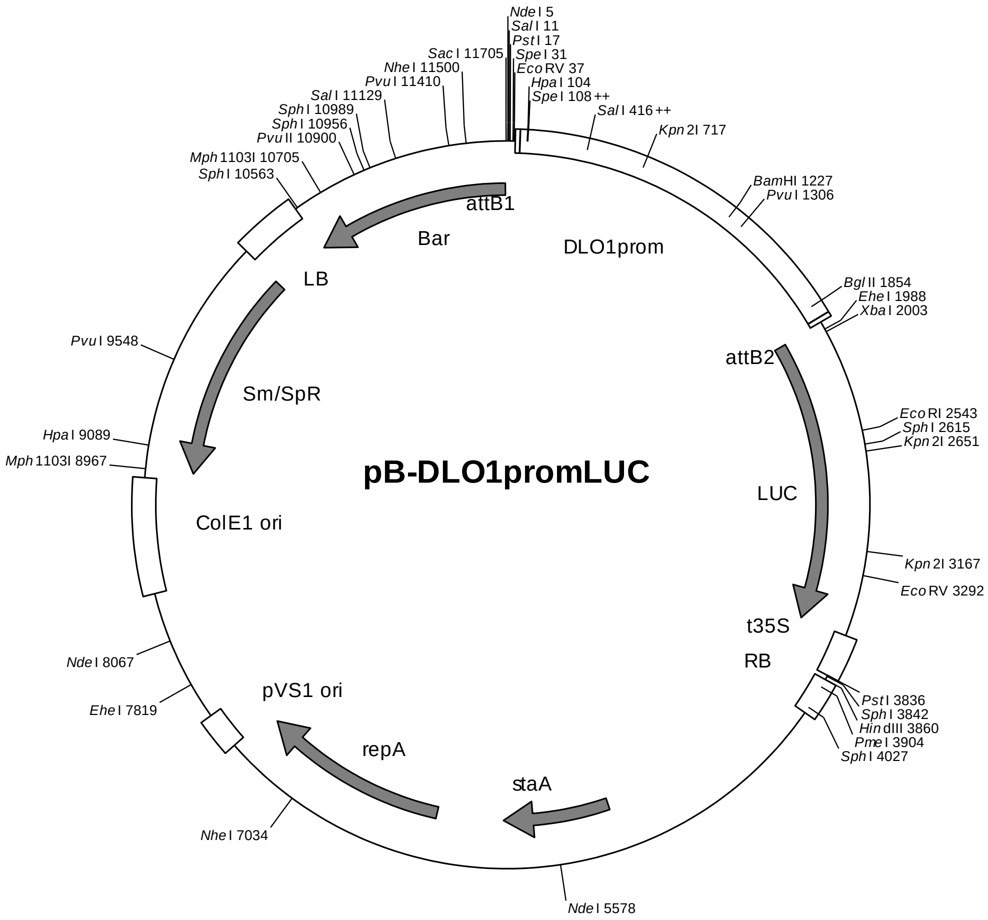

attB1: 43-67
*DLO1* promoter (-1777 to +77): 68-1921
A-box: 114-119
*TGACG* motif: 1773-1780
attB2: 1922-1946
firefly luciferase coding region (*LUC*): 1956-3608
35S terminator: 3612-3837
Right border: 3861-4060
pVS1 replicon (*sta*A, *rep*A, ori): 5246-7630
ColE1 ori: 8921-8322
Spectinomycin resistance cassette: 10213-8964
Left border: 10219-10551
Basta resistance cassette: 11700-10555

Position relative to transcriptional start site +1:

| | | | |
|---|---|---|---|
| *A-box* | -1731 | TACGTA | -1726 |
| *A-box* mutated | | TTTTTC | |

| | | | |
|---|---|---|---|
| *TGACGTCA* | -72 | TGACGTCA | -65 |
| *TGACGTCA* mutated | | TTTTTTCA | |

sequence (wild-type):
CTCCCATATGGTCGACCTGCAGGCGGCCGCACTAGTGATATCACAAGTTTGTACAAAAAAGCAGGCTAACTAATT
TACGTGTTCTCCACCATAGGTTATATCTGTTAACTAGTTACGTACGTCGACCAAAAACATATTACTTTATCGTGG
TTTTATCGTTGAATTATTCACTTTCATGTTGGGATATCAAGATTTGAATCTTTCAATCATGATTTTAATTATAGT
TGTCAAATGTCATATCAAAAATATCTATTTTCTTATAGTTTTTAACTGATTAGTATGAGTTTCGATTTGGAATAT
TTCAGATATGAACATACAACTAAGGAACAAAATATTGATATTAGTTTCCTATATCTTTTTATTTATTTTAAATAC
AATAGCAGATCACATGTATATATTTTAACTCAATAGTTACGTCGACCAAAAACATATAACTTTAAATATCGTGGT
TTCATTGTTGATTTATTCACTTTCAGGTTGGGACATCGGAATTTGAATTTTTCAATACGAATTTTAAACATAGTT
AAATATTTTCTTATAGTTCTTACTTCTTAACGATTAGTATGAGTTTAGATTTGGAATATTTTAAATATGAACATA
TAGTTAAGGAACGAAAGGCGATCACATGTTTGTAGTTCACCTAAGTATTGTAGTTATAATCTCATCCAGTACCTG
GAGCAGTTGAGCTAATCTTTAGATTAAGCTTGGGCAAAAAATCCGGACTAAAATACTTGAACCGAAACCGATCCA
AAAAATCGGTTTGGGACGAGTACGGATATTACCTATTTACTTTATTGAGTCTTGAATTGTCGTACCCGTGAGTAT
AGGGTAAAATCCAACCCGAACCTGATACTCGAACATGTATCCGCTTACCCGAAATTACATATACTTATTCATAGA
TTTATATATATAAAATCTTATTTTATATTTAATATTTGAACTAAATACATTTATATCCAAATATCTAAAAATAAT
TACCAAAAACTAAATACATTTGTAAATACTTAAGACTAAATATTTTTATACGCAAGATACTTATAAGTTTTTTGG
ATATTGGGACAAAATTTCGGTTATGTGGGTAAAATTCAGATATTTGGGTAAAATATTCAGGTATTTAGATAAAAT
TTTAGATATTTGGCTATAAAATACCTGAATCTCAAAGATACCCATTTTTTTGGTATTTAGCGGTTCTAAAATTAT
ATATCCGAACCGACCCGAACCCCATTGGATCCAAACCGAAACAAAACAGAAATTTTAGATTTATCTTATTGGGTC
CTAAACTTTTCTACCCAAAAAATTCTTATCCGATCGAATTTTACCCGAATATTCCGACCCACATACCTGAATGCC
CAACCGTACTTTAGGGAATTACATCCGATGCACAAAAGAACTAACAATTTTGTACTATCATTAACACCACCCATT
TGCAACTCCGAAAGACGATCACATATTGTAGATTTAAAATGCAATCTTAATCACATTTTCACTAAAAGTTTTCCT
TAACGTATTTGGTGGGCATGATTTAACAGCTCAATTTGGCAGGTCAACGAATCAACCGTGCTTTAAACGTTGTAA
TGTTTTTCAACTTAAGATAAGACTCGAAATCAATTAAAAAAACAAACAAAAATCATAGTCATTGTTTTTCTTTGT
TAATTCATATACACTCATACAAAACATTGACAACTTATCTATTGTTTAATCTAAATAATACAATTATGTACAAGT
GTTACTTTAGTAATACTGCCGTCCATATAACGGCACAAAACGGAGAATGACGTCACCATTGTTGATGTCACACCT
CCATGTAACTTCTGTTATAAACGCATTAAACGTTACACTTGTGTATATTATATAGATCTCTGTCCCTTTATATTT
CATCTTTTGACATTACAAAATTACATTACCAAACACATTAAATGGCACCCAGCTTTCTTGTACAAAGTGGTGATA
AAAAAATGGAAGACGCCAAAAACATAAAGAAAGGCCCGGCGCCATTCTATCCTCTAGAGGATGGAACCGCTGGAG
AGCAACTGCATAAGGCTATGAAGAGATACGCCCTGGTTCCTGGAACAATTGCTTTTACAGATGCACATATCGAGG
TGAACATCACGTACGCGGAATACTTCGAAATGTCCGTTCGGTTGGCAGAAGCTATGAAACGATATGGGCTGAATA
CAAATCACAGAATCGTCGTATGCAGTGAAAACTCTCTTCAATTCTTTATGCCGGTGTTGGGCGCGTTATTTATCG
GAGTTGCAGTTGCGCCCGCGAACGACATTTATAATGAACGTGAATTGCTCAACAGTATGAACATTTCGCAGCCTA
CCGTAGTGTTTGTTTCCAAAAAGGGGTTGCAAAAAATTTTGAACGTGCAAAAAAAATTACCAATAATCCAGAAAA
TTATTATCATGGATTCTAAAACGGATTACCAGGGATTTCAGTCGATGTACACGTTCGTCACATCTCATCTACCTC
CCGGTTTTAATGAATACGATTTTGTACCAGAGTCCTTTGATCGTGACAAAACAATTGCACTGATAATGAATTCCT
CTGGATCTACTGGGTTACCTAAGGGTGTGGCCCTTCCGCATAGAACTGCCTGCGTCAGATTCTCGCATGCCAGAG
ATCCTATTTTTGGCAATCAAATCATTCCGGATACTGCGATTTTAAGTGTTGTTCCATTCCATCACGGTTTTGGAA
TGTTTACTACACTCGGATATTTGATATGTGGATTTCGAGTCGTCTTAATGTATAGATTTGAAGAAGAGCTGTTTT
TACGATCCCTTCAGGATTACAAAATTCAAAGTGCGTTGCTAGTACCAACCCTATTTTCATTCTTCGCCAAAAGCA
CTCTGATTGACAAATACGATTTATCTAATTTACACGAAATTGCTTCTGGGGGCGCACCTCTTTCGAAAGAAGTCG
GGGAAGCGGTTGCAAAACGCTTCCATCTTCCAGGGATACGACAAGGATATGGGCTCACTGAGACTACATCAGCTA
TTCTGATTACACCCGAGGGGGATGATAAACCGGGCGCGGTCGGTAAAGTTGTTCCATTTTTTGAAGCGAAGGTTG
TGGATCTGGATACCGGGAAAACGCTGGGCGTTAATCAGAGAGGCGAATTATGTGTCAGAGGACCTATGATTATGT
CCGGTTATGTAAACAATCCGGAAGCGACCAACGCCTTGATTGACAAGGATGGATGGCTACATTCTGGAGACATAG
CTTACTGGGACGAAGACGAACACTTCTTCATAGTTGACCGCTTGAAGTCTTTAATTAAATACAAAGGATATCAGG
TGGCCCCCGCTGAATTGGAATCGATATTGTTACAACACCCCAACATCTTCGACGCGGGCGTGGCAGGTCTTCCCG
ACGATGACGCCGGTGAACTTCCCGCCGCCGTTGTTGTTTTGGAGCACGGAAAGACGATGACGGAAAAAGAGATCG
TGGATTACGTCGCCAGTCAAGTAACAACCGCGAAAAAGTTGCGCGGAGGAGTTGTGTTTGTGGACGAAGTACCGA
AAGGTCTTACCGGAAAACTCGACGCAAGAAAAATCAGAGAGATCCTCATAAAGGCCAAGAAGGGCGGAAAGTCCA
AATTGTAACCGCGGCCATGCTAGAGTCCGCAAAAATCACCAGTCTCTCTCTACAAATCTATCTCTCTCTATTTTT
CTCCAGAATAATGTGTGAGTAGTTCCCAGATAAGGGAATTAGGGTTCTTATAGGGTTTCGCTCATGTGTTGAGCA
TATAAGAAACCCTTAGTATGTATTTGTATTTGTAAAATACTTCTATCAATAAAATTTCTAATTCCTAAAACCAAA
ATCCAGTGACCTGCAGGCATGCGACGTCGGGCCCAAGCTTAGCTTGAGCTTGGATCAGATTGTCGTTTCCCGCCT
TCAGTTTAAACTATCAGTGTTTGACAGGATATATTGGCGGGTAAACCTAAGAGAAAAGAGCGTTTATTAGAATAA
CGGATATTTAAAAGGGCGTGAAAAGGTTTATCCGTTCGTCCATTTGTATGTGCATGCCAACCACAGGGTTCCCCT
CGGGATCAAAGTACTTTGATCCAACCCCTCCGCTGCTATAGTGCAGTCGGCTTCTGACGTTCAGTGCAGCCGTCT
TCTGAAAACGACATGTCGCACAAGTCCTAAGTTACGCGACAGGCTGCCGCCCTGCCCTTTTCCTGGCGTTTTCTT
GTCGCGTGTTTTAGTCGCATAAAGTAGAATACTTGCGACTAGAACCGGAGACATTACGCCATGAACAAGAGCGCC
GCCGCTGGCCTGCTGGGCTATGCCCGCGTCAGCACCGACGACCAGGACTTGACCAACCAACGGGCCGAACTGCAC
GCGGCCGGCTGCACCAAGCTGTTTTCCGAGAAGATCACCGGCACCAGGCGCGACCGCCCGGAGCTGGCCAGGATG
CTTGACCACCTACGCCCTGGCGACGTTGTGACAGTGACCAGGCTAGACCGCCTGGCCCGCAGCACCCGCGACCTA
CTGGACATTGCCGAGCGCATCCAGGAGGCCGGCGCGGGCCTGCGTAGCCTGGCAGAGCCGTGGGCCGACACCACC

ACGCCGGCCGGCCGCATGGTGTTGACCGTGTTCGCCGGCATTGCCGAGTTCGAGCGTTCCCTAATCATCGACCGC
ACCCGGAGCGGGCGCGAGGCCGCCAAGGCCCGAGGCGTGAAGTTTGGCCCCCGCCCTACCCTCACCCCGGCACAG
ATCGCGCACGCCCGCGAGCTGATCGACCAGGAAGGCCGCACCGTGAAAGAGGCGGCTGCACTGCTTGGCGTGCAT
CGCTCGACCCTGTACCGCGCACTTGAGCGCAGCGAGGAAGTGACGCCCACCGAGGCCAGGCGGCGCGGTGCCTTC
CGTGAGGACGCATTGACCGAGGCCGACGCCCTGGCGGCCGCCGAGAATGAACGCCAAGAGGAACAAGCATGAAAC
CGCACCAGGACGGCCAGGACGAACCGTTTTTCATTACCGAAGAGATCGAGGCGGAGATGATCGCGGCCGGGTACG
TGTTCGAGCCGCCCGCGCACGTCTCAACCGTGCGGCTGCATGAAATCCTGGCCGGTTTGTCTGATGCCAAGCTGG
CGGCCTGGCCGGCCAGCTTGGCCGCTGAAGAAACCGAGCGCCGCCGTCTAAAAAGGTGATGTGTATTTGAGTAAA
ACAGCTTGCGTCATGCGGTCGCTGCGTATATGATGCGATGAGTAAATAAACAAATACGCAAGGGGAACGCATGAA
GGTTATCGCTGTACTTAACCAGAAAGGCGGGTCAGGCAAGACGACCATCGCAACCCATCTAGCCCGCGCCCTGCA
ACTCGCCGGGGCCGATGTTCTGTTAGTCGATTCCGATCCCCAGGGCAGTGCCCGCGATTGGGCGGCCGTGCGGGA
AGATCAACCGCTAACCGTTGTCGGCATCGACCGCCCGACGATTGACCGCGACGTGAAGGCCATCGGCCGGCGCGA
CTTCGTAGTGATCGACGGAGCGCCCCAGGCGGCGGACTTGGCTGTGTCCGCGATCAAGGCAGCCGACTTCGTGCT
GATTCCGGTGCAGCCAAGCCCTTACGACATATGGGCCACCGCCGACCTGGTGGAGCTGGTTAAGCAGCGCATTGA
GGTCACGGATGGAAGGCTACAAGCGGCCTTTGTCGTGTCGCGGGCGATCAAAGGCACGCGCATCGGCGGTGAGGT
TGCCGAGGCGCTGGCCGGGTACGAGCTGCCCATTCTTGAGTCCCGTATCACGCAGCGCGTGAGCTACCCAGGCAC
TGCCGCCGCCGGCACAACCGTTCTTGAATCAGAACCCGAGGGCGACGCTGCCCGCGAGGTCCAGGCGCTGGCCGC
TGAAATTAAATCAAAACTCATTTGAGTTAATGAGGTAAAGAGAAAATGAGCAAAAGCACAAACACGCTAAGTGCC
GGCCGTCCGAGCGCACGCAGCAGCAAGGCTGCAACGTTGGCCAGCCTGGCAGACACGCCAGCCATGAAGCGGGTC
AACTTTCAGTTGCCGGCGGAGGATCACACCAAGCTGAAGATGTACGCGGTACGCCAAGGCAAGACCATTACCGAG
CTGCTATCTGAATACATCGCGCAGCTACCAGAGTAAATGAGCAAATGAATAAATGAGTAGATGAATTTTAGCGGC
TAAAGGAGGCGGCATGGAAAATCAAGAACAACCAGGCACCGACGCCGTGGAATGCCCCATGTGTGGAGGAACGGG
CGGTTGGCCAGGCGTAAGCGGCTGGGTTGTCTGCCGGCCCTGCAATGGCACTGGAACCCCCAAGCCCGAGGAATC
GGCGTGACGGTCGCAAACCATCCGGCCCGGTACAAATCGGCGCGGCGCTGGGTGATGACCTGGTGGAGAAGTTGA
AGGCCGCGCAGGCCGCCCAGCGGCAACGCATCGAGGCAGAAGCACGCCCCGGTGAATCGTGGCAAGCGGCCGCTG
ATCGAATCCGCAAAGAATCCCGGCAACCGCCGGCAGCCGGTGCGCCGTCGATTAGGAAGCCGCCCAAGGGCGACG
AGCAACCAGATTTTTTCGTTCCGATGCTCTATGACGTGGGCACCCGCGATAGTCGCAGCATCATGGACGTGGCCG
TTTTCCGTCTGTCGAAGCGTGACCGACGAGCTGGCGAGGTGATCCGCTACGAGCTTCCAGACGGGCACGTAGAGG
TTTCCGCAGGGCCGGCCGGCATGGCCAGTGTGTGGGATTACGACCTGGTACTGATGGCGGTTTCCCATCTAACCG
AATCCATGAACCGATACCGGGAAGGGAAGGGAGACAAGCCCGGCCGCGTGTTCCGTCCACACGTTGCGGACGTAC
TCAAGTTCTGCCGGCGAGCCGATGGCGGAAAGCAGAAAGACGACCTGGTAGAAACCTGCATTCGGTTAAACACCA
CGCACGTTGCCATGCAGCGTACGAAGAAGGCCAAGAACGGCCGCCTGGTGACGGTATCCGAGGGTGAAGCCTTGA
TTAGCCGCTACAAGATCGTAAAGAGCGAAACCGGGCGGCCGGAGTACATCGAGATCGAGCTAGCTGATTGGATGT
ACCGCGAGATCACAGAAGGCAAGAACCCGGACGTGCTGACGGTTCACCCCGATTACTTTTTGATCGATCCCGGCA
TCGGCCGTTTTCTCTACCGCCTGGCACGCCGCGCCGCAGGCAAGGCAGAAGCCAGATGGTTGTTCAAGACGATCT
ACGAACGCAGTGGCAGCGCCGGAGAGTTCAAGAAGTTCTGTTTCACCGTGCGCAAGCTGATCGGGTCAAATGACC
TGCCGGAGTACGATTTGAAGGAGGAGGCGGGGCAGGCTGGCCCGATCCTAGTCATGCGCTACCGCAACCTGATCG
AGGGCGAAGCATCCGCCGGTTCCTAATGTACGGAGCAGATGCTAGGGCAAATTGCCCTAGCAGGGGAAAAAGGTC
GAAAAGGTCTCTTTCCTGTGGATAGCACGTACATTGGGAACCCAAAGCCGTACATTGGGAACCGGAACCCGTACA
TTGGGAACCCAAAGCCGTACATTGGGAACCGGTCACACATGTAAGTGACTGATATAAAAGAGAAAAAAGGCGATT
TTTCCGCCTAAAACTCTTTAAAACTTATTAAAACTCTTAAAACCCGCCTGGCCTGTGCATAACTGTCTGGCCAGC
GCACAGCCGAAGAGCTGCAAAAAGCGCCTACCCTTCGGTCGCTGCGCTCCCTACGCCCCGCCGCTTCGCGTCGGC
CTATCGCGGCCGCTGGCCGCTCAAAAATGGCTGGCCTACGGCCAGGCAATCTACCAGGGCGCGGACAAGCCGCGC
CGTCGCCACTCGACCGCCGGCGCCCACATCAAGGCACCCTGCCTCGCGCGTTTCGGTGATGACGGTGAAAACCTC
TGACACATGCAGCTCCCGGAGACGGTCACAGCTTGTCTGTAAGCGGATGCCGGGAGCAGACAAGCCCGTCAGGGC
GCGTCAGCGGGTGTTGGCGGGTGTCGGGGCGCAGCCATGACCCAGTCACGTAGCGATAGCGGAGTGTATACTGGC
TTAACTATGCGGCATCAGAGCAGATTGTACTGAGAGTGCACCATATGCGGTGTGAAATACCGCACAGATGCGTAA
GGAGAAAATACCGCATCAGGCGCTCTTCCGCTTCCTCGCTCACTGACTCGCTGCGCTCGGTCGTTCGGCTGCGGC
GAGCGGTATCAGCTCACTCAAAGGCGGTAATACGGTTATCCACAGAATCAGGGGATAACGCAGGAAAGAACATGT
GAGCAAAAGGCCAGCAAAAGGCCAGGAACCGTAAAAAGGCCGCGTTGCTGGCGTTTTTCCATAGGCTCCGCCCCC
CTGACGAGCATCACAAAAATCGACGCTCAAGTCAGAGGTGGCGAAACCCGACAGGACTATAAAGATACCAGGCGT
TTCCCCCTGGAAGCTCCCTCGTGCGCTCTCCTGTTCCGACCCTGCCGCTTACCGGATACCTGTCCGCCTTTCTCC
CTTCGGGAAGCGTGGCGCTTTCTCATAGCTCACGCTGTAGGTATCTCAGTTCGGTGTAGGTCGTTCGCTCCAAGC
TGGGCTGTGTGCACGAACCCCCCGTTCAGCCCGACCGCTGCGCCTTATCCGGTAACTATCGTCTTGAGTCCAACC
CGGTAAGACACGACTTATCGCCACTGGCAGCAGCCACTGGTAACAGGATTAGCAGAGCGAGGTATGTAGGCGGTG
CTACAGAGTTCTTGAAGTGGTGGCCTAACTACGGCTACACTAGAAGGACAGTATTTGGTATCTGCGCTCTGCTGA
AGCCAGTTACCTTCGGAAAAAGAGTTGGTAGCTCTTGATCCGGCAAACAAACCACCGCTGGTAGCGGTGGTTTTT
TTGTTTGCAAGCAGCAGATTACGCGCAGAAAAAAAGGATCTCAAGAAGATCCTTTGATCTTTTCTACGGGGTCTG
ACGCTCAGTGGAACGAAAACTCACGTTAAGGGATTTTGGTCATGCATGATATATCTCCCAATTTGTGTAGGGCTT
ATTATGCACGCTTAAAAATAATAAAAGCAGACTTGACCTGATAGTTTGGCTGTGAGCAATTATGTGCTTAGTGCA
TCTAATCGCTTGAGTTAACGCCGGCGAAGCGGCGTCGGCTTGAACGAATTTCTAGCTAGACATTATTTGCCGACT
ACCTTGGTGATCTCGCCTTTCACGTAGTGGACAAATTCTTCCAACTGATCTGCGCGCGAGGCCAAGCGATCTTCT
TCTTGTCCAAGATAAGCCTGTCTAGCTTCAAGTATGACGGGCTGATACTGGGCCGGCAGGCGCTCCATTGCCCAG

TCGGCAGCGACATCCTTCGGCGCGATTTTGCCGGTTACTGCGCTGTACCAAATGCGGGACAACGTAAGCACTACA
TTTCGCTCATCGCCAGCCCAGTCGGGCGGCGAGTTCCATAGCGTTAAGGTTTCATTTAGCGCCTCAAATAGATCC
TGTTCAGGAACCGGATCAAAGAGTTCCTCCGCCGCTGGACCTACCAAGGCAACGCTATGTTCTCTTGCTTTTGTC
AGCAAGATAGCCAGATCAATGTCGATCGTGGCTGGCTCGAAGATACCTGCAAGAATGTCATTGCGCTGCCATTCT
CCAAATTGCAGTTCGCGCTTAGCTGGATAACGCCACGGAATGATGTCGTCGTGCACAACAATGGTGACTTCTACA
GCGCGGAGAATCTCGCTCTCTCCAGGGGAAGCCGAAGTTTCCAAAAGGTCGTTGATCAAAGCTCGCCGCGTTGTT
TCATCAAGCCTTACGGTCACCGTAACCAGCAAATCAATATCACTGTGTGGCTTCAGGCCGCCATCCACTGCGGAG
CCGTACAAATGTACGGCCAGCAACGTCGGTTCGAGATGGCGCTCGATGACGCCAACTACCTCTGATAGTTGAGTC
GATACTTCGGCGATCACCGCTTCCCCCATGATGTTTAACTTTGTTTTAGGGCGACTGCCCTGCTGCGTAACATCG
TTGCTGCTCCATAACATCAAACATCGACCCACGGCGTAACGCGCTTGCTGCTTGGATGCCCGAGGCATAGACTGT
ACCCCAAAAAAACATGTCATAACAAGAAGCCATGAAAACCGCCACTGCGCCGTTACCACCGCTGCGTTCGGTCAA
GGTTCTGGACCAGTTGCGTGACGGCAGTTACGCTACTTGCATTACAGCTTACGAACCGAACGAGGCTTATGTCCA
CTGGGTTCGTGCCCGAATTGATCACAGGCAGCAACGCTCTGTCATCGTTACAATCAACATGCTACCCTCCGCGAG
ATCATCCGTGTTTCAAACCCGGCAGCTTAGTTGCCGTTCTTCCGAATAGCATCGGTAACATGAGCAAAGTCTGCC
GCCTTACAACGGCTCTCCCGCTGACGCCGTCCCGGACTGATGGGCTGCCTGTATCGAGTGGTGATTTTGTGCCGA
GCTGCCGGTCGGGGAGCTGTTGGCTGGCTGGTGGCAGGATATATTGTGGTGTAAACAAATTGACGCTTAGACAAC
TTAATAACACATTGCGGACGTTTTTAATGTACTGAATTAACGCCGAATTGAATTATCAGCTTGCATGCCGGTCGA
TCTAGTAACATAGATGACACCGCGCGCGATAATTTATCCTAGTTTGCGCGCTATATTTTGTTTTCTATCGCGTAT
TAAATGTATAATTGCGGGACTCTAATCATAAAAACCCATCTCATAAATAACGTCATGCATTACATGTTAATTATT
ACATGCTTAACGTAATTCAACAGAAATTATATGATAATCATCGCAAGACCGGCAACAGGATTCAATCTTAAGAAA
CTTTATTGCCAAATGTTTGAACGATCTGCTTGACTCTAGGGGTCATCAGATTTCGGTGACGGGCAGGACCGGACG
GGGCGGCACCGGCAGGCTGAAGTCCAGCTGCCAGAAACCCACGTCATGCCAGTTCCCGTGCTTGAAGCCGGCCGC
CCGCAGCATGCCGCGGGGGGCATATCCGAGCGCCTCGTGCATGCGCACGCTCGGGTCGTTGGGCAGCCCGATGAC
AGCGACCACGCTCTTGAAGCCCTGTGCCTCCAGGGACTTCAGCAGGTGGGTGTAGAGCGTGGAGCCCAGTCCCGT
CCGCTGGTGGCGGGGGGAGACGTACACGGTCGACTCGGCCGTCCAGTCGTAGGCGTTGCGTGCCTTCCAGGGACC
CGCGTAGGCGATGCCGGCGACCTCGCCGTCCACCTCGGCGACGAGCCAGGGATAGCGCTCCCGCAGACGGACGAG
GTCGTCCGTCCACTCCTGCGGTTCCTGCGGCTCGGTACGGAAGTTGACCGTGCTTGTCTCGATGTAGTGGTTGAC
GATGGTGCAGACCGCCGGCATGTCCGCCTCGGTGGCACGGCGGATGTCGGCCGGGCGTCGTTCTGGGCTCATGGT
AGATCCCCTCGATCGAGTTGAGAGTGAATATGAGACTCTAATTGGATACCGAGGGGAATTTATGGAACGTCAGTG
GAGCATTTTTGACAAGAAATATTTGCTAGCTGATAGTGACCTTAGGCGACTTTTGAACGCGCAATAATGGTTTCT
GACGTATGTGCTTAGCTCATTAAACTCCAGAAACCCGCGGCTCAGTGGCTCCTTCAACGTTGCGGTTCTGTCAGT
TCCAAACGTAAAACGGCTTGTCCCGCGTCATCGGCGGGGGTCATAACGTGACTCCCTTAATTCTCATGTATGATA
ATTCGAGCT

### pB-HAgTGA1

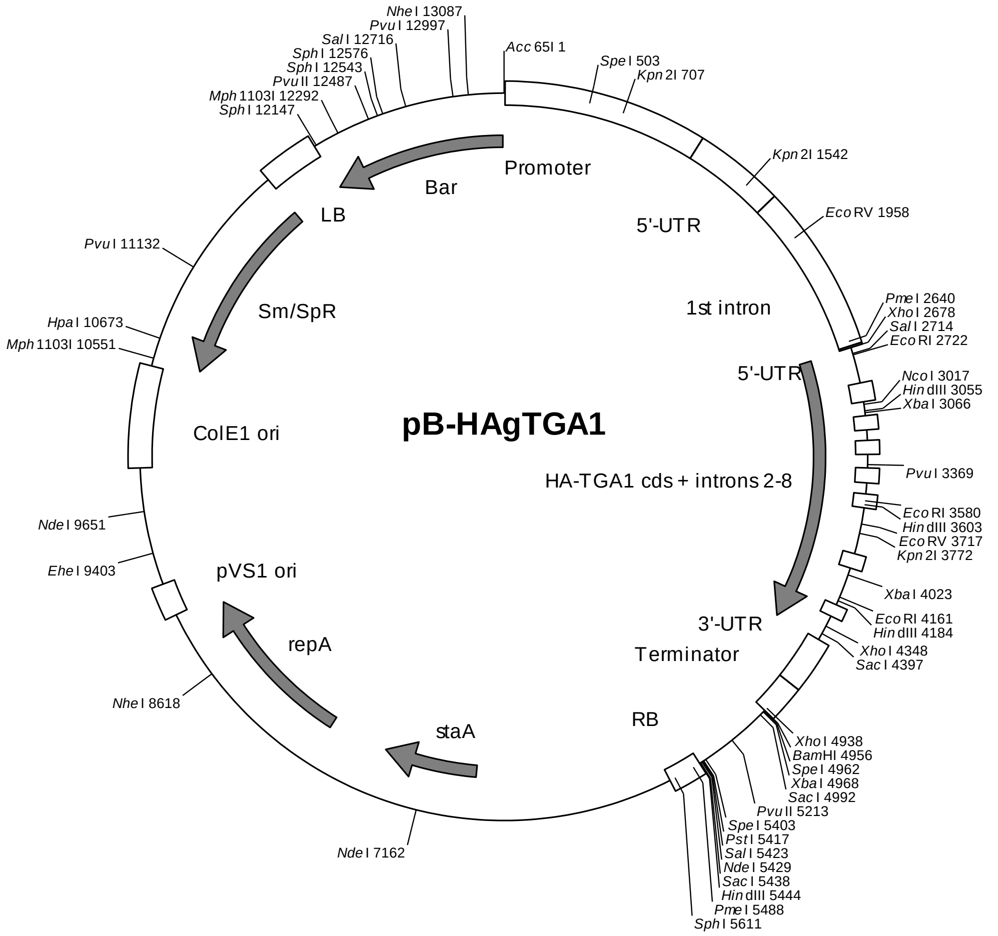

*TGA1* promoter (-1174 to -1): 7-1180
*TGA1* 5’-UTR: 1181-1701
*TGA1* intron 1: 1702-2667
*TGA1* 5’-UTR: 2668-2678
1xHA-tag: 2684-2719
*TGA1* coding region: 2720-2890
*TGA1* intron 2: 2891-3002
*TGA1* coding region: 3003-3083
*TGA1* intron 3: 3084-3167
*TGA1* coding region: 3168-3227
*TGA1* intron 4: 3228-3309
*TGA1* coding region: 3310-3387
*TGA1* intron 5: 3388-3475
*TGA1* coding region: 3476-3539
*TGA1* intron 6: 3540-3621
*TGA1* coding region: 3622-3893
*TGA1* intron 7: 3894-3981
*TGA1* coding region: 3982-4209
*TGA1* intron 8: 4210-4283
*TGA1* coding region: 4284-4436
*TGA1* 3'-UTR: 4437-4739
*TGA1* terminator: 4740-4956
Right border: 5445-5644
pVS1 replicon (*sta*A, *rep*A, ori): 6830-9214
ColE1 ori: 10505-9906
Spectinomycin resistance cassette: 11797-10548
Left border: 11803-12135
Basta resistance cassette: 13287-12139

sequence:
GGTACCTATGTCTCCTATGGAGCCTGAAGAGCCAAGCTGTTTTGATACATTAACTGGCTGCACACATATTAAATA
GGTGATTCCTTCACCTAATAAACTCTACTAACATTAGCCTGTTTAGACTAATTCCAAAGACTAATCAGTTTGTTT
AAAATTATGCTCGTATTATATAGCTACGATTTATATTATAAACAAGAAGAATATATGCAACTTTCTTGCAATATA
AATTAATGTTATCATCTTTACACTAAAATACTAACTATATTCCACTAATTAATACAAATCCGGTTAGCTATGAAC
AAAATGATCAATATCTTATTGAATTTTAGAATAAATAAACGTTGCTTGTTTGAACTGTTCATTCAGGTCGGTCGC
TACTCGCTAATTTGCTTAACCAATATTATTATATCAGCATGTAATAATATATTATAGCTAGTTGTTAAAAAAAAG
AATTATATATAGTTTTGTTTTCTTATACTAAATTCTACACTTTTTTTTGTCAACTAGTCTGATTTTTTTTAATGA
GACAGCTATGATATTTGGATAAACATCAGTATTTTTTTTTAGGTTCTCTGCATTTATATGTTTTAAAAACAGGTT
GATGAGATAGAAAAAAATGAAATATTGATTACAGATTAGATTAGTAAGATTACTTAATTATTTATTTAGAAATAG
CCGTTACTACGTCACCAGAATCGAAGCAGACTCCGGAGTTCTCTAATACTCCATTTGGCTACGCCGCGAGTAGTA
AACACGCGCAATATTTATCCTCTTTAGACGCTAAATTCCTTCGCTTGGAAGTATGCCGGCGTTTAGTGTTTACAT
TAGGCGCGTGTATTTACGCTTGCCGTCGAAACGAATTTTATGTTATCGCCGACGAATGCGCAGGTTAGAAGCAGA
CATGCGCAAGATAGAAAATATAAAATTATAAGGAAAAGAAATTATAAATTGGAGAGAATTTAGAAGAGAATAAAA
AAATGGATCTACCTGGTTCGCACATCAGCGTGCTAGGTAGTGGAGACATGACAATTGAATTTTTTAAAAACGAAT
AACTAAATATTCATAATCAAATTATTATCTTAAAAAAAATATTAAGATTCGATTGGTATCAGTGGATCATACATA
TTAAGTCGCTGCTAATCAAATTTCCTTTTCACTAATTTATTTCCTGCTTTTTTCCTTAGCACCATCAAACAAAAA
GTTTATGTGAAAAAACGTATTTTTTCCCACAAACAGATAATTAGTCCGGTGTGTTTTGTATATAGAAAGTGGAAG
AAAAAAATAGTATAGAAAAATGTAATAATTGGTTTTGACTCACTGTAGCTGAGTTATGCGTGAAGACAAAAATCA
CCGAGAAACCACAGAAACAAATAGAAAAAATAGTTAAATATATAAAACAAATATAAAATAACTGCGTCAAAGGTC
TCATTTAAGTTGTCGTGATTCTTCTTCCTTCGTCTACTTGAAGATGATGAAACCTGAGTTTCTGAGACAGAAGGA
AGAGCTTTTAACATCCCTTCTGTTTATGGTCGGATTCCAACTCCGGACTTCTTGGCTTTAACCCTTTACGTCCGA
TCTTGTCGCTGAAATCTTTCTTTCTCCTTTTACATGGTTTTCACGACGACAAAAGTTTGCCTTTTTCTAGTTGCT
GCCTTGACTTCAGATAGCATCACAAGGTTCGGCTTCAGATAACATCACAAGGTTTGATTGACTTTGCTCTGTTCT
AATCTAATCTTGTTAATATCTTAAAAGCCTAAAACTTTTCTCTTATTGATTGTTTTGGTTAATTTTGTTTCCCCT
TTTTTTTGTTGTATTTACTCGATTGATTACTTCAATATGATTTTCTGGTTTGCTTGGAATTTAAGCTGGAAGGAA
CTGATTCGAATATTTCTTGCATAAGAAGATGTTTCAATCTACCTATCTATTGACTTCATGATTGGCTTTTGCTTT
TATTGTTGATATCTCTTACTATCTTGATTTCACCTTTCGTTACCATTTGCGTACAAAAATTTAACAAAGCAATGA
ATGGGAATTTCTTATTTTTTGTATAGTCGTTTTAACTGCTAAACTATGTTAAGTTGGTCCACACAATTAGGTGCT
GAAAATAAAGTGACATCCTAATAAGTGGAAACTTGACTTATGGAGCTTTTCGTCTGCTTAACATTAATTTTTATC
ATCTTATCTATTTGACACCTTCCAAACACTATTGGGGAAATGGCTTCTTCTCTTTTTTCCGGTGTAGGTTTAAGG
TTTTAGATTTGAAGGCGGATGGTGAGGAGTTTGTGTTGATGAAATGGGTGATTTGAAGTTAAATCCTTTAAGTTC
TTTTAGCCTTAAGTTCTTTTAGCCTGTCATTACAATATATGTTTCAGTTGTGATCTTTTGGTGCTCTCCAAGGTC
TTAGGGAATCTCCGTGTCCCCTCTGGTTTCTTTCTTTGTATAAAGTCTCTGGAATTTAAAGATGATTTTGTCCTG
TTTTCTTCGAAGAATTTGGCGAAGAAACATCAAAAGTTCACTATTTGCTTTACGTGTTATGCTTGTTTGACTTTG
TGTGATCTAACTTTTGGTTGAAAACTAATGCTCATCTTTTGTTCTTTGGAATGTCTTGATCTGCTTGATTTTGAA
TGCTTTGTCACATTGTTTAAACAATTTGTTCTTTGTTTTCAGTTGAGGAAAACTCGAGATGTACCCATACGATGT
TCCAGACTACGCTGTCGACATGAATTCGACATCGACACATTTTGTGCCACCGAGAAGAGTTGGTATATACGAACC
TGTCCATCAATTCGGTATGTGGGGGGAGAGTTTCAAAAGCAATATTAGCAATGGGACTATGAACACACCAAACCA
CATAATAATACCGAATAATCAGAAACTAGACAACAACGTGGTACTTTTTTACTTCATTGCTCTTCAACCTCCTGA
TGATAAATTTGTTTGGTCTGTTTTCAGAAATTTAAACTAGCTTTCATTTCTTGACAATCTTTTTTTTTTTTTTAT
AGTCAGAGGATACTTCCCATGGAACAGCAGGAACTCCTCACATGTTCGATCAAGAAGCTTCAACGTCTAGACATC
CCGATAAGGTTAGCTATTCAAATACTTCTCTATCAGTTTCTTAATCATAAATTTCTACGCATCATATAGACTAAA
TCGTTATATCGTTACAGATACAAAGACGGCTTGCTCAAAACCGCGAGGCTGCTAGGAAAAGTCGCTTGCGCAAGA
AGGTATGACTTCATTTTCTTCCCCTTGGTAAACTGTTTCTGTTTGCCTAATGAGTTGAGAGATTTGATTGTTTTT
TTTTTGCAGGCTTATGTTCAGCAACTGGAAACAAGCAGGTTGAAGCTAATTCAATTAGAGCAAGAACTCGATCGT
GCTAGACAACAGGTTAATGAAATGAAATCTAAATATATATTCGCTTTCTTTCGAGTTCCAAAATATAATGCAATT
TCTGATTGAATATGAATGTTTTCAGGGATTCTATGTAGGAAACGGAATAGATACTAATTCTCTCGGTTTTTCGGA
AACCATGAATCCAGGTTTTGAATCAAGAACAATGAGTCTTTTGGAAAAAATATAGAATTCAAGATTTTGATTGGT
GGAAGCTTATTTGATTTGCAGGGATTGCTGCATTTGAAATGGAATATGGACATTGGGTTGAAGAACAGAACAGAC
AGATATGTGAACTAAGAACAGTTTTACACGGACACATTAACGATATCGAGCTTCGTTCGCTAGTCGAAAACGCCA
TGAAACATTACTTTGAGCTTTTCCGGATGAAATCGTCTGCTGCCAAAGCCGATGTCTTCTTCGTCATGTCAGGGA
TGTGGAGAACTTCAGCAGAACGATTCTTCTTATGGATTGGCGGATTTCGACCCTCCGATCTTCTCAAGGTATAAA
ACAAATAACAATCTAGCGTAATGTTTAATGAATAGAAGGAAACAAGCAAGATTACAATGTTTCAAATGTTTTTTT
TTGTAGGTTCTTTTGCCACATTTTGATGTCTTGACGGATCAACAACTTCTAGATGTATGCAATCTAAAACAATCG
TGTCAGCAAGCAGAAGACGCGTTGACTCAAGGTATGGAGAAGCTGCAACACACCCTTGCGGACTGCGTTGCAGCG
GGACAACTCGGTGAAGGAAGTTACATTCCTCAGGTGAATTCTGCTATGGATAGATTAGAAGCTTTGGTCAGTTTC
GTAAATCAGGTAAAATATATATATTCAAACTCATTTGAAAATACTATAGTATCAGTTTCTTGATTTTGTTTGTTC
ATTCTTAGGCTGATCACTTGAGACATGAAACATTGCAACAAATGTATCGGATATTGACAACGCGACAAGCGGCTC
GAGGATTATTAGCTCTTGGTGAGTATTTTCAACGGCTTAGAGCCTTGAGCTCAAGTTGGGCAACTCGACATCGTG
AACCAACGTAGGTTTGAGTTATTTTGTAACAACCAAATGAAGAAAATGGAAAGACCTCAAAAAATGAAGAATGAG
TGCATCTGAAAACAGAGGACTACTCTGAATAAATAGAGGGGTTGCTGCTGATATTTATTTTTACTCTGCGGCGGA

ATTAGAAAATTTGAAAAACATCATGTATTGATAAGTTGTAAATATCAGAAAAAGGTGGGGGTGCAAAAATTTGTA
CTTTTTAGCTTTTGAAAGAGGCAAGTTTTTCGAATGTTTGTTTGATTTGTAAACAATTTCAGAATTATATAAACT
TGGTTCCAAATCCCCTGTAATAATGTCGAGCTATCTGCAATTTGAAAACTATAGGGGCTTTACTTAATTTTACGT
TTTGTGTGACACCTATTTGATTTTTTTTGATACGTTTGGACCAGTTTCGTAATTAGAGGTAAATAAAGGGGCAAA
TAGTAAGATATAAGAAAAGCCTTATCTCATAGAGTTTCTTCTCTTCATTATCCTCAAAATCGCTCGAGAGTGAGA
ACTCTGGATCCACTAGTTCTAGAGCGGCCGCCACCGCGGTGGAGCTCCAGCTTTTGTTCCCTTTAGTGAGGGTTA
ATTCCGAGCTTGGCGTAATCATGGTCATAGCTGTTTCCTGTGTGAAATTGTTATCCGCTCACAATTCCACACAAC
ATACGAGCCGGAAGHCATAAAGTGTAAAGCCTGGGGTGCCTAATGAGTGAGCTAACTCACATTAATTGCGTTGCG
CTCACTGCCCGCTTTCCAGTCGGGAAACCTGTCGTGCCAGCTGCATTAATGAATCGGCCAACGCGCGGGGAGAGG
CGGTTTGCGTATTGGGCGCTCTTCCGCTTCCTCGCTCACTGACTCGCTGCGCTCGGTCGTTCGGCTGCGGCGAGC
GGTATCAGCTCACTCAAAGGCGGTAATACGGTTATCCACAGAATCAGGGGATAACGCAGGAAAGAACATGAAGGA
TCACTAGTGCGGCCGCCTGCAGGTCGACCATATGGGAGAGCTCAAGCTTAGCTTGAGCTTGGATCAGATTGTCGT
TTCCCGCCTTCAGTTTAAACTATCAGTGTTTGACAGGATATATTGGCGGGTAAACCTAAGAGAAAAGAGCGTTTA
TTAGAATAACGGATATTTAAAAGGGCGTGAAAAGGTTTATCCGTTCGTCCATTTGTATGTGCATGCCAACCACAG
GGTTCCCCTCGGGATCAAAGTACTTTGATCCAACCCCTCCGCTGCTATAGTGCAGTCGGCTTCTGACGTTCAGTG
CAGCCGTCTTCTGAAAACGACATGTCGCACAAGTCCTAAGTTACGCGACAGGCTGCCGCCCTGCCCTTTTCCTGG
CGTTTTCTTGTCGCGTGTTTTAGTCGCATAAAGTAGAATACTTGCGACTAGAACCGGAGACATTACGCCATGAAC
AAGAGCGCCGCCGCTGGCCTGCTGGGCTATGCCCGCGTCAGCACCGACGACCAGGACTTGACCAACCAACGGGCC
GAACTGCACGCGGCCGGCTGCACCAAGCTGTTTTCCGAGAAGATCACCGGCACCAGGCGCGACCGCCCGGAGCTG
GCCAGGATGCTTGACCACCTACGCCCTGGCGACGTTGTGACAGTGACCAGGCTAGACCGCCTGGCCCGCAGCACC
CGCGACCTACTGGACATTGCCGAGCGCATCCAGGAGGCCGGCGCGGGCCTGCGTAGCCTGGCAGAGCCGTGGGCC
GACACCACCACGCCGGCCGGCCGCATGGTGTTGACCGTGTTCGCCGGCATTGCCGAGTTCGAGCGTTCCCTAATC
ATCGACCGCACCCGGAGCGGGCGCGAGGCCGCCAAGGCCCGAGGCGTGAAGTTTGGCCCCCGCCCTACCCTCACC
CCGGCACAGATCGCGCACGCCCGCGAGCTGATCGACCAGGAAGGCCGCACCGTGAAAGAGGCGGCTGCACTGCTT
GGCGTGCATCGCTCGACCCTGTACCGCGCACTTGAGCGCAGCGAGGAAGTGACGCCCACCGAGGCCAGGCGGCGC
GGTGCCTTCCGTGAGGACGCATTGACCGAGGCCGACGCCCTGGCGGCCGCCGAGAATGAACGCCAAGAGGAACAA
GCATGAAACCGCACCAGGACGGCCAGGACGAACCGTTTTTCATTACCGAAGAGATCGAGGCGGAGATGATCGCGG
CCGGGTACGTGTTCGAGCCGCCCGCGCACGTCTCAACCGTGCGGCTGCATGAAATCCTGGCCGGTTTGTCTGATG
CCAAGCTGGCGGCCTGGCCGGCCAGCTTGGCCGCTGAAGAAACCGAGCGCCGCCGTCTAAAAAGGTGATGTGTAT
TTGAGTAAAACAGCTTGCGTCATGCGGTCGCTGCGTATATGATGCGATGAGTAAATAAACAAATACGCAAGGGGA
ACGCATGAAGGTTATCGCTGTACTTAACCAGAAAGGCGGGTCAGGCAAGACGACCATCGCAACCCATCTAGCCCG
CGCCCTGCAACTCGCCGGGGCCGATGTTCTGTTAGTCGATTCCGATCCCCAGGGCAGTGCCCGCGATTGGGCGGC
CGTGCGGGAAGATCAACCGCTAACCGTTGTCGGCATCGACCGCCCGACGATTGACCGCGACGTGAAGGCCATCGG
CCGGCGCGACTTCGTAGTGATCGACGGAGCGCCCCAGGCGGCGGACTTGGCTGTGTCCGCGATCAAGGCAGCCGA
CTTCGTGCTGATTCCGGTGCAGCCAAGCCCTTACGACATATGGGCCACCGCCGACCTGGTGGAGCTGGTTAAGCA
GCGCATTGAGGTCACGGATGGAAGGCTACAAGCGGCCTTTGTCGTGTCGCGGGCGATCAAAGGCACGCGCATCGG
CGGTGAGGTTGCCGAGGCGCTGGCCGGGTACGAGCTGCCCATTCTTGAGTCCCGTATCACGCAGCGCGTGAGCTA
CCCAGGCACTGCCGCCGCCGGCACAACCGTTCTTGAATCAGAACCCGAGGGCGACGCTGCCCGCGAGGTCCAGGC
GCTGGCCGCTGAAATTAAATCAAAACTCATTTGAGTTAATGAGGTAAAGAGAAAATGAGCAAAAGCACAAACACG
CTAAGTGCCGGCCGTCCGAGCGCACGCAGCAGCAAGGCTGCAACGTTGGCCAGCCTGGCAGACACGCCAGCCATG
AAGCGGGTCAACTTTCAGTTGCCGGCGGAGGATCACACCAAGCTGAAGATGTACGCGGTACGCCAAGGCAAGACC
ATTACCGAGCTGCTATCTGAATACATCGCGCAGCTACCAGAGTAAATGAGCAAATGAATAAATGAGTAGATGAAT
TTTAGCGGCTAAAGGAGGCGGCATGGAAAATCAAGAACAACCAGGCACCGACGCCGTGGAATGCCCCATGTGTGG
AGGAACGGGCGGTTGGCCAGGCGTAAGCGGCTGGGTTGTCTGCCGGCCCTGCAATGGCACTGGAACCCCCAAGCC
CGAGGAATCGGCGTGACGGTCGCAAACCATCCGGCCCGGTACAAATCGGCGCGGCGCTGGGTGATGACCTGGTGG
AGAAGTTGAAGGCCGCGCAGGCCGCCCAGCGGCAACGCATCGAGGCAGAAGCACGCCCCGGTGAATCGTGGCAAG
CGGCCGCTGATCGAATCCGCAAAGAATCCCGGCAACCGCCGGCAGCCGGTGCGCCGTCGATTAGGAAGCCGCCCA
AGGGCGACGAGCAACCAGATTTTTTCGTTCCGATGCTCTATGACGTGGGCACCCGCGATAGTCGCAGCATCATGG
ACGTGGCCGTTTTCCGTCTGTCGAAGCGTGACCGACGAGCTGGCGAGGTGATCCGCTACGAGCTTCCAGACGGGC
ACGTAGAGGTTTCCGCAGGGCCGGCCGGCATGGCCAGTGTGTGGGATTACGACCTGGTACTGATGGCGGTTTCCC
ATCTAACCGAATCCATGAACCGATACCGGGAAGGGAAGGGAGACAAGCCCGGCCGCGTGTTCCGTCCACACGTTG
CGGACGTACTCAAGTTCTGCCGGCGAGCCGATGGCGGAAAGCAGAAAGACGACCTGGTAGAAACCTGCATTCGGT
TAAACACCACGCACGTTGCCATGCAGCGTACGAAGAAGGCCAAGAACGGCCGCCTGGTGACGGTATCCGAGGGTG
AAGCCTTGATTAGCCGCTACAAGATCGTAAAGAGCGAAACCGGGCGGCCGGAGTACATCGAGATCGAGCTAGCTG
ATTGGATGTACCGCGAGATCACAGAAGGCAAGAACCCGGACGTGCTGACGGTTCACCCCGATTACTTTTTGATCG
ATCCCGGCATCGGCCGTTTTCTCTACCGCCTGGCACGCCGCGCCGCAGGCAAGGCAGAAGCCAGATGGTTGTTCA
AGACGATCTACGAACGCAGTGGCAGCGCCGGAGAGTTCAAGAAGTTCTGTTTCACCGTGCGCAAGCTGATCGGGT
CAAATGACCTGCCGGAGTACGATTTGAAGGAGGAGGCGGGGCAGGCTGGCCCGATCCTAGTCATGCGCTACCGCA
ACCTGATCGAGGGCGAAGCATCCGCCGGTTCCTAATGTACGGAGCAGATGCTAGGGCAAATTGCCCTAGCAGGGG
AAAAAGGTCGAAAAGGTCTCTTTCCTGTGGATAGCACGTACATTGGGAACCCAAAGCCGTACATTGGGAACCGGA
ACCCGTACATTGGGAACCCAAAGCCGTACATTGGGAACCGGTCACACATGTAAGTGACTGATATAAAAGAGAAAA
AAGGCGATTTTTCCGCCTAAAACTCTTTAAAACTTATTAAAACTCTTAAAACCCGCCTGGCCTGTGCATAACTGT
CTGGCCAGCGCACAGCCGAAGAGCTGCAAAAAGCGCCTACCCTTCGGTCGCTGCGCTCCCTACGCCCCGCCGCTT

CGCGTCGGCCTATCGCGGCCGCTGGCCGCTCAAAAATGGCTGGCCTACGGCCAGGCAATCTACCAGGGCGCGGAC
AAGCCGCGCCGTCGCCACTCGACCGCCGGCGCCCACATCAAGGCACCCTGCCTCGCGCGTTTCGGTGATGACGGT
GAAAACCTCTGACACATGCAGCTCCCGGAGACGGTCACAGCTTGTCTGTAAGCGGATGCCGGGAGCAGACAAGCC
CGTCAGGGCGCGTCAGCGGGTGTTGGCGGGTGTCGGGGCGCAGCCATGACCCAGTCACGTAGCGATAGCGGAGTG
TATACTGGCTTAACTATGCGGCATCAGAGCAGATTGTACTGAGAGTGCACCATATGCGGTGTGAAATACCGCACA
GATGCGTAAGGAGAAAATACCGCATCAGGCGCTCTTCCGCTTCCTCGCTCACTGACTCGCTGCGCTCGGTCGTTC
GGCTGCGGCGAGCGGTATCAGCTCACTCAAAGGCGGTAATACGGTTATCCACAGAATCAGGGGATAACGCAGGAA
AGAACATGTGAGCAAAAGGCCAGCAAAAGGCCAGGAACCGTAAAAAGGCCGCGTTGCTGGCGTTTTTCCATAGGC
TCCGCCCCCCTGACGAGCATCACAAAAATCGACGCTCAAGTCAGAGGTGGCGAAACCCGACAGGACTATAAAGAT
ACCAGGCGTTTCCCCCTGGAAGCTCCCTCGTGCGCTCTCCTGTTCCGACCCTGCCGCTTACCGGATACCTGTCCG
CCTTTCTCCCTTCGGGAAGCGTGGCGCTTTCTCATAGCTCACGCTGTAGGTATCTCAGTTCGGTGTAGGTCGTTC
GCTCCAAGCTGGGCTGTGTGCACGAACCCCCCGTTCAGCCCGACCGCTGCGCCTTATCCGGTAACTATCGTCTTG
AGTCCAACCCGGTAAGACACGACTTATCGCCACTGGCAGCAGCCACTGGTAACAGGATTAGCAGAGCGAGGTATG
TAGGCGGTGCTACAGAGTTCTTGAAGTGGTGGCCTAACTACGGCTACACTAGAAGGACAGTATTTGGTATCTGCG
CTCTGCTGAAGCCAGTTACCTTCGGAAAAAGAGTTGGTAGCTCTTGATCCGGCAAACAAACCACCGCTGGTAGCG
GTGGTTTTTTTGTTTGCAAGCAGCAGATTACGCGCAGAAAAAAAGGATCTCAAGAAGATCCTTTGATCTTTTCTA
CGGGGTCTGACGCTCAGTGGAACGAAAACTCACGTTAAGGGATTTTGGTCATGCATGATATATCTCCCAATTTGT
GTAGGGCTTATTATGCACGCTTAAAAATAATAAAAGCAGACTTGACCTGATAGTTTGGCTGTGAGCAATTATGTG
CTTAGTGCATCTAATCGCTTGAGTTAACGCCGGCGAAGCGGCGTCGGCTTGAACGAATTTCTAGCTAGACATTAT
TTGCCGACTACCTTGGTGATCTCGCCTTTCACGTAGTGGACAAATTCTTCCAACTGATCTGCGCGCGAGGCCAAG
CGATCTTCTTCTTGTCCAAGATAAGCCTGTCTAGCTTCAAGTATGACGGGCTGATACTGGGCCGGCAGGCGCTCC
ATTGCCCAGTCGGCAGCGACATCCTTCGGCGCGATTTTGCCGGTTACTGCGCTGTACCAAATGCGGGACAACGTA
AGCACTACATTTCGCTCATCGCCAGCCCAGTCGGGCGGCGAGTTCCATAGCGTTAAGGTTTCATTTAGCGCCTCA
AATAGATCCTGTTCAGGAACCGGATCAAAGAGTTCCTCCGCCGCTGGACCTACCAAGGCAACGCTATGTTCTCTT
GCTTTTGTCAGCAAGATAGCCAGATCAATGTCGATCGTGGCTGGCTCGAAGATACCTGCAAGAATGTCATTGCGC
TGCCATTCTCCAAATTGCAGTTCGCGCTTAGCTGGATAACGCCACGGAATGATGTCGTCGTGCACAACAATGGTG
ACTTCTACAGCGCGGAGAATCTCGCTCTCTCCAGGGGAAGCCGAAGTTTCCAAAAGGTCGTTGATCAAAGCTCGC
CGCGTTGTTTCATCAAGCCTTACGGTCACCGTAACCAGCAAATCAATATCACTGTGTGGCTTCAGGCCGCCATCC
ACTGCGGAGCCGTACAAATGTACGGCCAGCAACGTCGGTTCGAGATGGCGCTCGATGACGCCAACTACCTCTGAT
AGTTGAGTCGATACTTCGGCGATCACCGCTTCCCCCATGATGTTTAACTTTGTTTTAGGGCGACTGCCCTGCTGC
GTAACATCGTTGCTGCTCCATAACATCAAACATCGACCCACGGCGTAACGCGCTTGCTGCTTGGATGCCCGAGGC
ATAGACTGTACCCCAAAAAAACATGTCATAACAAGAAGCCATGAAAACCGCCACTGCGCCGTTACCACCGCTGCG
TTCGGTCAAGGTTCTGGACCAGTTGCGTGACGGCAGTTACGCTACTTGCATTACAGCTTACGAACCGAACGAGGC
TTATGTCCACTGGGTTCGTGCCCGAATTGATCACAGGCAGCAACGCTCTGTCATCGTTACAATCAACATGCTACC
CTCCGCGAGATCATCCGTGTTTCAAACCCGGCAGCTTAGTTGCCGTTCTTCCGAATAGCATCGGTAACATGAGCA
AAGTCTGCCGCCTTACAACGGCTCTCCCGCTGACGCCGTCCCGGACTGATGGGCTGCCTGTATCGAGTGGTGATT
TTGTGCCGAGCTGCCGGTCGGGGAGCTGTTGGCTGGCTGGTGGCAGGATATATTGTGGTGTAAACAAATTGACGC
TTAGACAACTTAATAACACATTGCGGACGTTTTTAATGTACTGAATTAACGCCGAATTGAATTATCAGCTTGCAT
GCCGGTCGATCTAGTAACATAGTAGATGACACCGCGCGCGATAATTTATCCTAGTTTGCGCGCTATATTTTGTTT
TCTATCGCGTATTAAATGTATAATTGCGGGACTCTAATCATAAAAACCCATCTCATAAATAACGTCATGCATTAC
ATGTTAATTATTACATGCTTAACGTAATTCAACAGAAATTATATGATAATCATCGCAAGACCGGCAACAGGATTC
AATCTTAAGAAACTTTATTGCCAAATGTTTGAACGATCTGCTTGACTCTAGGGGTCATCAGATTTCGGTGACGGG
CAGGACCGGACGGGGCGGCACCGGCAGGCTGAAGTCCAGCTGCCAGAAACCCACGTCATGCCAGTTCCCGTGCTT
GAAGCCGGCCGCCCGCAGCATGCCGCGGGGGGCATATCCGAGCGCCTCGTGCATGCGCACGCTCGGGTCGTTGGG
CAGCCCGATGACAGCGACCACGCTCTTGAAGCCCTGTGCCTCCAGGGACTTCAGCAGGTGGGTGTAGAGCGTGGA
GCCCAGTCCCGTCCGCTGGTGGCGGGGGGAGACGTACACGGTCGACTCGGCCGTCCAGTCGTAGGCGTTGCGTGC
CTTCCAGGGACCCGCGTAGGCGATGCCGGCGACCTCGCCGTCCACCTCGGCGACGAGCCAGGGATAGCGCTCCCG
CAGACGGACGAGGTCGTCCGTCCACTCCTGCGGTTCCTGCGGCTCGGTACGGAAGTTGACCGTGCTTGTCTCGAT
GTAGTGGTTGACGATGGTGCAGACCGCCGGCATGTCCGCCTCGGTGGCACGGCGGATGTCGGCCGGGCGTCGTTC
TGGGCTCATGGTAGATCCCCTCGATCGAGTTGAGAGTGAATATGAGACTCTAATTGGATACCGAGGGGAATTTAT
GGAACGTCAGTGGAGCATTTTTGACAAGAAATATTTGCTAGCTGATAGTGACCTTAGGCGACTTTTGAACGCGCA
ATAATGGTTTCTGACGTATGTGCTTAGCTCATTAAACTCCAGAAACCCGCGGCTCAGTGGCTCCTTCAACGTTGC
GGTTCTGTCAGTTCCAAACGTAAAACGGCTTGTCCCGCGTCATCGGCGGGGGTCATAACGTGACTCCCTTAATTC
TCATGTATGATAATTCGAG

#### Mutant *gTGA1(C172N/C260N/C266S/C287S)*

pB-HAgTGA1, bps 3651-3710

```
3651  CAT TGG GTT GAA GAA CAG AAC AGA CAG ATA TGT GAA CTA AGA ACA GTT TTA CAC GGA CAC
      His Trp Val Glu Glu Gln Asn Arg Gln Ile Cys Glu Leu Arg Thr Val Leu His Gly His
```

pB-HAgTGA1(C172N/C260N/C266S/C287S), bps 3651-3710

```
3651  CAT TGG GTT GAA GAA CAG AAC AGA CAG ATA AAT GAA CTA AGA ACA GTT TTA CAC GGA CAC
      His Trp Val Glu Glu Gln Asn Arg Gln Ile Asn Glu Leu Arg Thr Val Leu His Gly His
```

pB-HAgTGA1, bps 4021-4080

```
4021  CTT CTA GAT GTA TGC AAT CTA AAA CAA TCG TGT CAG CAA GCA GAA GAC GCG TTG ACT CAA
      Leu Leu Asp Val Cys Asn Leu Lys Gln Ser Cys Gln Gln Ala Glu Asp Ala Leu Thr Gln
```

pB-HAgTGA1(C172N/C260N/C266S/C287S), bps 4021-4080

```
4021  CTT CTA GAT GTA AAC AAT CTA AAA CAA TCG TCT CAG CAA GCA GAA GAC GCG TTG ACT CAA
      Leu Leu Asp Val Asn Asn Leu Lys Gln Ser Ser Gln Gln Ala Glu Asp Ala Leu Thr Gln
```

pB-HAgTGA1, bps 4081-4140

```
4081  GGT ATG GAG AAG CTG CAA CAC ACC CTT GCG GAC TGC GTT GCA GCG GGA CAA CTC GGT GAA
      Gly Met Glu Lys Leu Gln His Thr Leu Ala Asp Cys Val Ala Ala Gly Gln Leu Gly Glu
```

pB-HAgTGA1(C172N/C260N/C266S/C287S), bps 4081-4140

```
4081  GGT ATG GAG AAG CTG CAA CAC ACC CTT GCG GAC TCC GTT GCA GCG GGA CAA CTC GGT GAA
      Gly Met Glu Lys Leu Gln His Thr Leu Ala Asp Ser Val Ala Ala Gly Gln Leu Gly Glu
```

### Control plasmid pB-HA

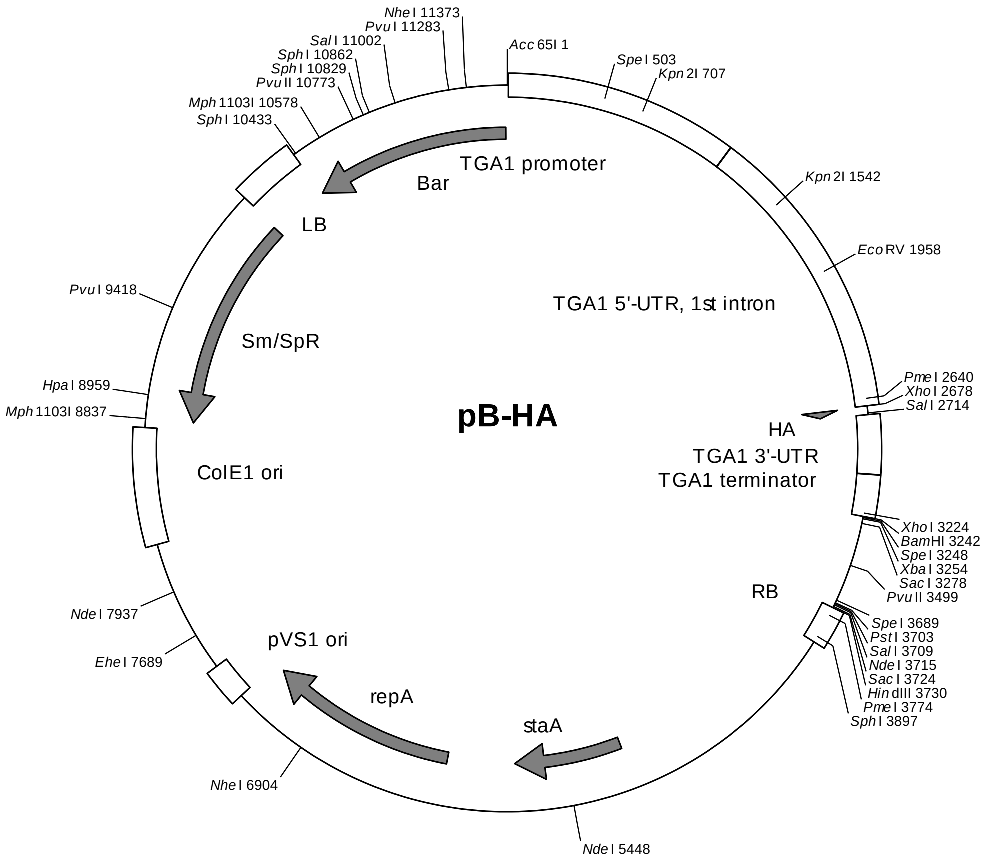

*TGA1* promoter (-1174 to -1): 7-1180
*TGA1* 5'-UTR, intron 1: 1181-2678
1xHA-tag: 2684-2722
*TGA1* 3'-UTR: 2723-3025
*TGA1* terminator: 3026-3242
Right border: 3731-3930
pVS1 replicon (*sta*A, *rep*A, ori): 5116-7500
ColE1 ori: 8791-8192
Spectinomycin resistance cassette: 10083-8834
Left border: 10089-10421
Basta resistance cassette: 11573-10425

sequence:

```
GGTACCTATGTCTCCTATGGAGCCTGAAGAGCCAAGCTGTTTTGATACATTAACTGGCTGCACACATATTAAATA
GGTGATTCCTTCACCTAATAAACTCTACTAACATTAGCCTGTTTAGACTAATTCCAAAGACTAATCAGTTTGTTT
AAAATTATGCTCGTATTATATAGCTACGATTTATATTATAAACAAGAAGAATATATGCAACTTTCTTGCAATATA
AATTAATGTTATCATCTTTACACTAAAATACTAACTATATTCCACTAATTAATACAAATCCGGTTAGCTATGAAC
AAAATGATCAATATCTTATTGAATTTTAGAATAAATAAACGTTGCTTGTTTGAACTGTTCATTCAGGTCGGTCGC
TACTCGCTAATTTGCTTAACCAATATTATTATATCAGCATGTAATAATATATTATAGCTAGTTGTTAAAAAAAAG
AATTATATATAGTTTTGTTTTCTTATACTAAATTCTACACTTTTTTTTGTCAACTAGTCTGATTTTTTTTAATGA
GACAGCTATGATATTTGGATAAACATCAGTATTTTTTTTTAGGTTCTCTGCATTTATATGTTTTAAAAACAGGTT
GATGAGATAGAAAAAAATGAAATATTGATTACAGATTAGATTAGTAAGATTACTTAATTATTTATTTAGAAATAG
CCGTTACTACGTCACCAGAATCGAAGCAGACTCCGGAGTTCTCTAATACTCCATTTGGCTACGCCGCGAGTAGTA
AACACGCGCAATATTTATCCTCTTTAGACGCTAAATTCCTTCGCTTGGAAGTATGCCGGCGTTTAGTGTTTACAT
TAGGCGCGTGTATTTACGCTTGCCGTCGAAACGAATTTTATGTTATCGCCGACGAATGCGCAGGTTAGAAGCAGA
CATGCGCAAGATAGAAAATATAAAATTATAAGGAAAAGAAATTATAAATTGGAGAGAATTTAGAAGAGAATAAAA
```

AAATGGATCTACCTGGTTCGCACATCAGCGTGCTAGGTAGTGGAGACATGACAATTGAATTTTTTAAAAACGAAT
AACTAAATATTCATAATCAAATTATTATCTTAAAAAAAAATATTAAGATTCGATTGGTATCAGTGGATCATACATA
TTAAGTCGCTGCTAATCAAATTTCCTTTTCACTAATTTATTTCCTGCTTTTTTCCTTAGCACCATCAAACAAAAA
GTTTATGTGAAAAAACGTATTTTTTCCCACAAACAGATAATTAGTCCGGTGTGTTTTGTATATAGAAAGTGGAAG
AAAAAAATAGTATAGAAAAATGTAATAATTGGTTTTGACTCACTGTAGCTGAGTTATGCGTGAAGACAAAAATCA
CCGAGAAACCACAGAAACAAATAGAAAAAATAGTTAAATATATAAAACAAATATAAAATAACTGCGTCAAAGGTC
TCATTTAAGTTGTCGTGATTCTTCTTCCTTCGTCTACTTGAAGATGATGAAACCTGAGTTTCTGAGACAGAAGGA
AGAGCTTTTAACATCCCTTCTGTTTATGGTCGGATTCCAACTCCGGACTTCTTGGCTTTAACCCTTTACGTCCGA
TCTTGTCGCTGAAATCTTTCTTTCTCCTTTTACATGGTTTCACGACGACAAAAGTTTGCCTTTTTCTAGTTGCT
GCCTTGACTTCAGATAGCATCACAAGGTTCGGCTTCAGATAACATCACAAGGTTTGATTGACTTTGCTCTGTTCT
AATCTAATCTTGTTAATATCTTAAAAGCCTAAAACTTTTCTCTTATTGATTGTTTTGGTTAATTTTGTTTCCCCT
TTTTTTTGTTGTATTTACTCGATTGATTACTTCAATATGATTTTCTGGTTTGCTTGGAATTTAAGCTGGAAGGAA
CTGATTCGAATATTTCTTGCATAAGAAGATGTTTCAATCTACCTATCTATTGACTTCATGATTGGCTTTTGCTTT
TATTGTTGATATCTCTTACTATCTTGATTTCACCTTTCGTTACCATTTGCGTACAAAAATTTAACAAAGCAATGA
ATGGGAATTTCTTATTTTTTGTATAGTCGTTTTAACTGCTAAACTATGTTAAGTTGGTCCACACAATTAGGTGCT
GAAAATAAAGTGACATCCTAATAAGTGGAAACTTGACTTATGGAGCTTTTCGTCTGCTTAACATTAATTTTTATC
ATCTTATCTATTTGACACCTTCCAAACACTATTGGGGAAATGGCTTCTTCTCTTTTTTCCGGTGTAGGTTTAAGG
TTTTAGATTTGAAGGCGGATGGTGAGGAGTTTGTGTTGATGAAATGGGTGATTTGAAGTTAAATCCTTTAAGTTC
TTTTAGCCTTAAGTTCTTTTAGCCTGTCATTACAATATATGTTTCAGTTGTGATCTTTTGGTGCTCTCCAAGGTC
TTAGGGAATCTCCGTGTCCCCTCTGGTTTCTTTCTTTGTATAAAGTCTCTGGAATTTAAAGATGATTTTGTCCTG
TTTTCTTCGAAGAATTTGGCGAAGAAACATCAAAAGTTCACTATTTGCTTTACGTGTTATGCTTGTTTGACTTTG
TGTGATCTAACTTTTGGTTGAAAACTAATGCTCATCTTTTGTTCTTTGGAATGTCTTGATCTGCTTGATTTTGAA
TGCTTTGTCACATTGTTTAAACAATTTGTTCTTTGTTTTCAGTTGAGGAAAACTCGAGATGTACCCATACGATGT
TCCAGACTACGCTGTCGACTAGGTTTGAGTTATTTTGTAACAACCAAATGAAGAAAATGGAAAGACCTCAAAAAA
TGAAGAATGAGTGCATCTGAAAACAGAGGACTACTCTGAATAAATAGAGGGGTTGCTGCTGATATTTATTTTTAC
TCTGCGGCGGAATTAGAAAATTTGAAAAACATCATGTATTGATAAGTTGTAAATATCAGAAAAAGGTGGGGGTGC
AAAAATTTGTACTTTTTAGCTTTTGAAAGAGGCAAGTTTTTCGAATGTTTGTTTGATTTGTAAACAATTTCAGAA
TTATATAAACTTGGTTCCAAATCCCCTGTAATAATGTCGAGCTATCTGCAATTTGAAAACTATAGGGGCTTTACT
TAATTTTACGTTTTGTGTGACACCTATTTGATTTTTTTTGATACGTTTGGACCAGTTTCGTAATTAGAGGTAAAT
AAAGGGGCAAATAGTAAGATATAAGAAAAGCCTTATCTCATAGAGTTTCTTCTCTTCATTATCCTCAAAATCGCT
CGAGAGTGAGAACTCTGGATCCACTAGTTCTAGAGCGGCCGCCACCGCGGTGGAGCTCCAGCTTTTGTTCCCTTT
AGTGAGGGTTAATTCCGAGCTTGGCGTAATCATGGTCATAGCTGTTTCCTGTGTGAAATTGTTATCCGCTCACAA
TTCCACACAACATACGAGCCGGAAGHCATAAAGTGTAAAGCCTGGGGTGCCTAATGAGTGAGCTAACTCACATTA
ATTGCGTTGCGCTCACTGCCCGCTTTCCAGTCGGGAAACCTGTCGTGCCAGCTGCATTAATGAATCGGCCAACGC
GCGGGGAGAGGCGGTTTGCGTATTGGGCGCTCTTCCGCTTCCTCGCTCACTGACTCGCTGCGCTCGGTCGTTCGG
CTGCGGCGAGCGGTATCAGCTCACTCAAAGGCGGTAATACGGTTATCCACAGAATCAGGGGATAACGCAGGAAAG
AACATGAAGGATCACTAGTGCGGCCGCCTGCAGGTCGACCATATGGGAGAGCTCAAGCTTAGCTTGAGCTTGGAT
CAGATTGTCGTTTCCCGCCTTCAGTTTAAACTATCAGTGTTTGACAGGATATATTGGCGGGTAAACCTAAGAGAA
AAGAGCGTTTATTAGAATAACGGATATTTAAAAGGGCGTGAAAAGGTTTATCCGTTCGTCCATTTGTATGTGCAT
GCCAACCACAGGGTTCCCCTCGGGATCAAAGTACTTTGATCCAACCCCTCCGCTGCTATAGTGCAGTCGGCTTCT
GACGTTCAGTGCAGCCGTCTTCTGAAAACGACATGTCGCACAAGTCCTAAGTTACGCGACAGGCTGCCGCCCTGC
CCTTTTCCTGGCGTTTTCTTGTCGCGTGTTTTAGTCGCATAAAGTAGAATACTTGCGACTAGAACCGGAGACATT
ACGCCATGAACAAGAGCGCCGCCGCTGGCCTGCTGGGCTATGCCCGCGTCAGCACCGACGACCAGGACTTGACCA
ACCAACGGGCCGAACTGCACGCGGCCGGCTGCACCAAGCTGTTTTCCGAGAAGATCACCGGCACCAGGCGCGACC
GCCCGGAGCTGGCCAGGATGCTTGACCACCTACGCCCTGGCGACGTTGTGACAGTGACCAGGCTAGACCGCCTGG
CCCGCAGCACCCGCGACCTACTGGACATTGCCGAGCGCATCCAGGAGGCCGGCGCGGGCCTGCGTAGCCTGGCAG
AGCCGTGGGCCGACACCACCACGCCGGCCGGCCGCATGGTGTTGACCGTGTTCGCCGGCATTGCCGAGTTCGAGC
GTTCCCTAATCATCGACCGCACCCGGAGCGGGCGCGAGGCCGCCAAGGCCCGAGGCGTGAAGTTTGGCCCCCGCC
CTACCCTCACCCCGGCACAGATCGCGCACGCCCGCGAGCTGATCGACCAGGAAGGCCGCACCGTGAAAGAGGCGG
CTGCACTGCTTGGCGTGCATCGCTCGACCCTGTACCGCGCACTTGAGCGCAGCGAGGAAGTGACGCCCACCGAGG
CCAGGCGGCGCGGTGCCTTCCGTGAGGACGCATTGACCGAGGCCGACGCCCTGGCGGCCGCCGAGAATGAACGCC
AAGAGGAACAAGCATGAAACCGCACCAGGACGGCCAGGACGAACCGTTTTTCATTACCGAAGAGATCGAGGCGGA
GATGATCGCGGCCGGGTACGTGTTCGAGCCGCCCGCGCACGTCTCAACCGTGCGGCTGCATGAAATCCTGGCCGG
TTTGTCTGATGCCAAGCTGGCGGCCTGGCCGGCCAGCTTGGCCGCTGAAGAAACCGAGCGCCGCCGTCTAAAAAG
GTGATGTGTATTTGAGTAAAACAGCTTGCGTCATGCGGTCGCTGCGTATATGATGCGATGAGTAAATAAACAAAT
ACGCAAGGGGAACGCATGAAGGTTATCGCTGTACTTAACCAGAAAGGCGGGTCAGGCAAGACGACCATCGCAACC
CATCTAGCCCGCGCCCTGCAACTCGCCGGGGCCGATGTTCTGTTAGTCGATTCCGATCCCCAGGGCAGTGCCCGC
GATTGGGCGGCCGTGCGGGAAGATCAACCGCTAACCGTTGTCGGCATCGACCGCCCGACGATTGACCGCGACGTG
AAGGCCATCGGCCGGCGCGACTTCGTAGTGATCGACGGAGCGCCCCAGGCGGCGGACTTGGCTGTGTCCGCGATC
AAGGCAGCCGACTTCGTGCTGATTCCGGTGCAGCCAAGCCCTTACGACATATGGGCCACCGCCGACCTGGTGGAG
CTGGTTAAGCAGCGCATTGAGGTCACGGATGGAAGGCTACAAGCGGCCTTTGTCGTGTCGCGGGCGATCAAAGGC
ACGCGCATCGGCGGTGAGGTTGCCGAGGCGCTGGCCGGGTACGAGCTGCCCATTCTTGAGTCCCGTATCACGCAG
CGCGTGAGCTACCCAGGCACTGCCGCCGCCGGCACAACCGTTCTTGAATCAGAACCCGAGGGCGACGCTGCCCGC

GAGGTCCAGGCGCTGGCCGCTGAAATTAAATCAAAACTCATTTGAGTTAATGAGGTAAAGAGAAAATGAGCAAAA
GCACAAACACGCTAAGTGCCGGCCGTCCGAGCGCACGCAGCAGCAAGGCTGCAACGTTGGCCAGCCTGGCAGACA
CGCCAGCCATGAAGCGGGTCAACTTTCAGTTGCCGGCGGAGGATCACACCAAGCTGAAGATGTACGCGGTACGCC
AAGGCAAGACCATTACCGAGCTGCTATCTGAATACATCGCGCAGCTACCAGAGTAAATGAGCAAATGAATAAATG
AGTAGATGAATTTTAGCGGCTAAAGGAGGCGGCATGGAAAATCAAGAACAACCAGGCACCGACGCCGTGGAATGC
CCCATGTGTGGAGGAACGGGCGGTTGGCCAGGCGTAAGCGGCTGGGTTGTCTGCCGGCCCTGCAATGGCACTGGA
ACCCCCAAGCCCGAGGAATCGGCGTGACGGTCGCAAACCATCCGGCCCGGTACAAATCGGCGCGGCGCTGGGTGA
TGACCTGGTGGAGAAGTTGAAGGCCGCGCAGGCCGCCCAGCGGCAACGCATCGAGGCAGAAGCACGCCCCGGTGA
ATCGTGGCAAGCGGCCGCTGATCGAATCCGCAAAGAATCCCGGCAACCGCCGGCAGCCGGTGCGCCGTCGATTAG
GAAGCCGCCCAAGGGCGACGAGCAACCAGATTTTTTCGTTCCGATGCTCTATGACGTGGGCACCCGCGATAGTCG
CAGCATCATGGACGTGGCCGTTTTCCGTCTGTCGAAGCGTGACCGACGAGCTGGCGAGGTGATCCGCTACGAGCT
TCCAGACGGGCACGTAGAGGTTTCCGCAGGGCCGGCCGGCATGGCCAGTGTGTGGGATTACGACCTGGTACTGAT
GGCGGTTTCCCATCTAACCGAATCCATGAACCGATACCGGGAAGGGAAGGGAGACAAGCCCGGCCGCGTGTTCCG
TCCACACGTTGCGGACGTACTCAAGTTCTGCCGGCGAGCCGATGGCGGAAAGCAGAAAGACGACCTGGTAGAAAC
CTGCATTCGGTTAAACACCACGCACGTTGCCATGCAGCGTACGAAGAAGGCCAAGAACGGCCGCCTGGTGACGGT
ATCCGAGGGTGAAGCCTTGATTAGCCGCTACAAGATCGTAAAGAGCGAAACCGGGCGGCCGGAGTACATCGAGAT
CGAGCTAGCTGATTGGATGTACCGCGAGATCACAGAAGGCAAGAACCCGGACGTGCTGACGGTTCACCCCGATTA
CTTTTTGATCGATCCCGGCATCGGCCGTTTTCTCTACCGCCTGGCACGCCGCGCCGCAGGCAAGGCAGAAGCCAG
ATGGTTGTTCAAGACGATCTACGAACGCAGTGGCAGCGCCGGAGAGTTCAAGAAGTTCTGTTTCACCGTGCGCAA
GCTGATCGGGTCAAATGACCTGCCGGAGTACGATTTGAAGGAGGAGGCGGGGCAGGCTGGCCCGATCCTAGTCAT
GCGCTACCGCAACCTGATCGAGGGCGAAGCATCCGCCGGTTCCTAATGTACGGAGCAGATGCTAGGGCAAATTGC
CCTAGCAGGGGAAAAAGGTCGAAAAGGTCTCTTTCCTGTGGATAGCACGTACATTGGGAACCCAAAGCCGTACAT
TGGGAACCGGAACCCGTACATTGGGAACCCAAAGCCGTACATTGGGAACCGGTCACACATGTAAGTGACTGATAT
AAAAGAGAAAAAAGGCGATTTTTCCGCCTAAAACTCTTTAAAACTTATTAAAACTCTTAAAACCCGCCTGGCCTG
TGCATAACTGTCTGGCCAGCGCACAGCCGAAGAGCTGCAAAAAGCGCCTACCCTTCGGTCGCTGCGCTCCCTACG
CCCCGCCGCTTCGCGTCGGCCTATCGCGGCCGCTGGCCGCTCAAAAATGGCTGGCCTACGGCCAGGCAATCTACC
AGGGCGCGGACAAGCCGCGCCGTCGCCACTCGACCGCCGGCGCCCACATCAAGGCACCCTGCCTCGCGCGTTTCG
GTGATGACGGTGAAAACCTCTGACACATGCAGCTCCCGGAGACGGTCACAGCTTGTCTGTAAGCGGATGCCGGGA
GCAGACAAGCCCGTCAGGGCGCGTCAGCGGGTGTTGGCGGGTGTCGGGGCGCAGCCATGACCCAGTCACGTAGCG
ATAGCGGAGTGTATACTGGCTTAACTATGCGGCATCAGAGCAGATTGTACTGAGAGTGCACCATATGCGGTGTGA
AATACCGCACAGATGCGTAAGGAGAAAATACCGCATCAGGCGCTCTTCCGCTTCCTCGCTCACTGACTCGCTGCG
CTCGGTCGTTCGGCTGCGGCGAGCGGTATCAGCTCACTCAAAGGCGGTAATACGGTTATCCACAGAATCAGGGGA
TAACGCAGGAAAGAACATGTGAGCAAAAGGCCAGCAAAAGGCCAGGAACCGTAAAAAGGCCGCGTTGCTGGCGTT
TTTCCATAGGCTCCGCCCCCCTGACGAGCATCACAAAAATCGACGCTCAAGTCAGAGGTGGCGAAACCCGACAGG
ACTATAAAGATACCAGGCGTTTCCCCCTGGAAGCTCCCTCGTGCGCTCTCCTGTTCCGACCCTGCCGCTTACCGG
ATACCTGTCCGCCTTTCTCCCTTCGGGAAGCGTGGCGCTTTCTCATAGCTCACGCTGTAGGTATCTCAGTTCGGT
GTAGGTCGTTCGCTCCAAGCTGGGCTGTGTGCACGAACCCCCCGTTCAGCCCGACCGCTGCGCCTTATCCGGTAA
CTATCGTCTTGAGTCCAACCCGGTAAGACACGACTTATCGCCACTGGCAGCAGCCACTGGTAACAGGATTAGCAG
AGCGAGGTATGTAGGCGGTGCTACAGAGTTCTTGAAGTGGTGGCCTAACTACGGCTACACTAGAAGGACAGTATT
TGGTATCTGCGCTCTGCTGAAGCCAGTTACCTTCGGAAAAAGAGTTGGTAGCTCTTGATCCGGCAAACAAACCAC
CGCTGGTAGCGGTGGTTTTTTTGTTTGCAAGCAGCAGATTACGCGCAGAAAAAAAGGATCTCAAGAAGATCCTTT
GATCTTTTCTACGGGGTCTGACGCTCAGTGGAACGAAAACTCACGTTAAGGGATTTTGGTCATGCATGATATATC
TCCCAATTTGTGTAGGGCTTATTATGCACGCTTAAAAATAATAAAAGCAGACTTGACCTGATAGTTTGGCTGTGA
GCAATTATGTGCTTAGTGCATCTAATCGCTTGAGTTAACGCCGGCGAAGCGGCGTCGGCTTGAACGAATTTCTAG
CTAGACATTATTTGCCGACTACCTTGGTGATCTCGCCTTTCACGTAGTGGACAAATTCTTCCAACTGATCTGCGC
GCGAGGCCAAGCGATCTTCTTCTTGTCCAAGATAAGCCTGTCTAGCTTCAAGTATGACGGGCTGATACTGGGCCG
GCAGGCGCTCCATTGCCCAGTCGGCAGCGACATCCTTCGGCGCGATTTTGCCGGTTACTGCGCTGTACCAAATGC
GGGACAACGTAAGCACTACATTTCGCTCATCGCCAGCCCAGTCGGGCGGCGAGTTCCATAGCGTTAAGGTTTCAT
TTAGCGCCTCAAATAGATCCTGTTCAGGAACCGGATCAAAGAGTTCCTCCGCCGCTGGACCTACCAAGGCAACGC
TATGTTCTCTTGCTTTTGTCAGCAAGATAGCCAGATCAATGTCGATCGTGGCTGGCTCGAAGATACCTGCAAGAA
TGTCATTGCGCTGCCATTCTCCAAATTGCAGTTCGCGCTTAGCTGGATAACGCCACGGAATGATGTCGTCGTGCA
CAACAATGGTGACTTCTACAGCGCGGAGAATCTCGCTCTCTCCAGGGGAAGCCGAAGTTTCCAAAAGGTCGTTGA
TCAAAGCTCGCCGCGTTGTTTCATCAAGCCTTACGGTCACCGTAACCAGCAAATCAATATCACTGTGTGGCTTCA
GGCCGCCATCCACTGCGGAGCCGTACAAATGTACGGCCAGCAACGTCGGTTCGAGATGGCGCTCGATGACGCCAA
CTACCTCTGATAGTTGAGTCGATACTTCGGCGATCACCGCTTCCCCCATGATGTTTAACTTTGTTTTAGGGCGAC
TGCCCTGCTGCGTAACATCGTTGCTGCTCCATAACATCAAACATCGACCCACGGCGTAACGCGCTTGCTGCTTGG
ATGCCCGAGGCATAGACTGTACCCCAAAAAAACATGTCATAACAAGAAGCCATGAAAACCGCCACTGCGCCGTTA
CCACCGCTGCGTTCGGTCAAGGTTCTGGACCAGTTGCGTGACGGCAGTTACGCTACTTGCATTACAGCTTACGAA
CCGAACGAGGCTTATGTCCACTGGGTTCGTGCCCGAATTGATCACAGGCAGCAACGCTCTGTCATCGTTACAATC
AACATGCTACCCTCCGCGAGATCATCCGTGTTTCAAACCCGGCAGCTTAGTTGCCGTTCTTCCGAATAGCATCGG
TAACATGAGCAAAGTCTGCCGCCTTACAACGGCTCTCCCGCTGACGCCGTCCCGGACTGATGGGCTGCCTGTATC
GAGTGGTGATTTTGTGCCGAGCTGCCGGTCGGGGAGCTGTTGGCTGGCTGGTGGCAGGATATATTGTGGTGTAAA
CAAATTGACGCTTAGACAACTTAATAACACATTGCGGACGTTTTTAATGTACTGAATTAACGCCGAATTGAATTA

TCAGCTTGCATGCCGGTCGATCTAGTAACATAGTAGATGACACCGCGCGATAATTTATCCTAGTTTGCGCGCT
ATATTTTGTTTTCTATCGCGTATTAAATGTATAATTGCGGGACTCTAATCATAAAAACCCATCTCATAAATAACG
TCATGCATTACATGTTAATTATTACATGCTTAACGTAATTCAACAGAAATTATATGATAATCATCGCAAGACCGG
CAACAGGATTCAATCTTAAGAAACTTTATTGCCAAATGTTTGAACGATCTGCTTGACTCTAGGGGTCATCAGATT
TCGGTGACGGGCAGGACCGGACGGGGCGGCACCGGCAGGCTGAAGTCCAGCTGCCAGAAACCCACGTCATGCCAG
TTCCCGTGCTTGAAGCCGGCCGCCCGCAGCATGCCGCGGGGGGCATATCCGAGCGCCTCGTGCATGCGCACGCTC
GGGTCGTTGGGCAGCCCGATGACAGCGACCACGCTCTTGAAGCCCTGTGCCTCCAGGGACTTCAGCAGGTGGGTG
TAGAGCGTGGAGCCCAGTCCCGTCCGCTGGTGGCGGGGGGAGACGTACACGGTCGACTCGGCCGTCCAGTCGTAG
GCGTTGCGTGCCTTCCAGGGACCCGCGTAGGCGATGCCGGCGACCTCGCCGTCCACCTCGGCGACGAGCCAGGGA
TAGCGCTCCCGCAGACGGACGAGGTCGTCCGTCCACTCCTGCGGTTCCTGCGGCTCGGTACGGAAGTTGACCGTG
CTTGTCTCGATGTAGTGGTTGACGATGGTGCAGACCGCCGGCATGTCCGCCTCGGTGGCACGGCGGATGTCGGCC
GGGCGTCGTTCTGGGCTCATGGTAGATCCCCTCGATCGAGTTGAGAGTGAATATGAGACTCTAATTGGATACCGA
GGGGAATTTATGGAACGTCAGTGGAGCATTTTTGACAAGAAATATTTGCTAGCTGATAGTGACCTTAGGCGACTT
TTGAACGCGCAATAATGGTTTCTGACGTATGTGCTTAGCTCATTAAACTCCAGAAACCCGCGGCTCAGTGGCTCC
TTCAACGTTGCGGTTCTGTCAGTTCCAAACGTAAAACGGCTTGTCCCGCGTCATCGGCGGGGGTCATAACGTGAC
TCCCTTAATTCTCATGTATGATAATTCGAG
